# Systematic investigation of imprinted gene expression and enrichment in the mouse brain explored at single-cell resolution

**DOI:** 10.1101/2020.07.27.222893

**Authors:** M. J. Higgs, M. J. Hill, R. M. John, A. R. Isles

## Abstract

**Background:** Although a number of imprinted genes are known to be highly expressed in the brain, and in certain brain regions in particular, whether they are truly over-represented in the brain has never been formally tested. Using thirteen single-cell RNA sequencing datasets we systematically investigated imprinted gene over-representation at the organ, brain region, and cell-specific levels.

**Results:** We established that imprinted genes are indeed over-represented in the adult brain, and in neurons particularly compared to other brain cell-types. We then examined brain-wide datasets to test enrichment within distinct brain regions and neuron subpopulations and demonstrated over-representation of imprinted genes in the hypothalamus, ventral midbrain, pons and medulla. Finally, using datasets focusing on these regions of enrichment, we identified hypothalamic neuroendocrine populations and the monoaminergic hindbrain neurons as specific hotspots of imprinted gene expression.

**Conclusions:** These analyses provide the first robust assessment of the neural systems on which imprinted genes converge. Moreover, the unbiased approach, with each analysis informed by the findings of the previous level, permits highly informed inferences about the functions on which imprinted gene expression converges. Our findings indicate the neuronal regulation of motivated behaviours such as feeding and sleep, alongside the regulation of pituitary function as functional hotspots for imprinting, thus adding statistical rigour to prior assumptions and providing testable predictions for novel neural and behavioural phenotypes associated with specific genes and imprinted gene networks. In turn, this work sheds further light on the potential evolutionary drivers of genomic imprinting in the brain.

## BACKGROUND

Imprinted genes demonstrate a preferential or exclusively monoallelic expression from either the maternal or paternal allele in an epigenetically predetermined manner (a parent-of-origin effect, POE). To date approximately 260 imprinted genes, demonstrating biased allelic expression and/or associated with a parental-specific epigenetic mark, have been identified in the mouse (~230 in humans) (9, 10). This epigenetic regulation makes genomic imprinting an evolutionary puzzle as many of these genes are effectively haploid and thereby negate many of the benefits of diploidy (11). Studying the patterns of expression and function of imprinted genes may shed light on the drivers leading to the evolution of genomic imprinting. For instance, characterisation of a number of imprinted genes points to convergence on placental function (12), in line with the predictions of early theoretical ideas (13). Outside of the placenta, the brain consistently emerges as an adult tissue with a large number of expressed imprinted genes (14–16). However, given that it is estimated that ~80% of all genes in the genome are expressed in the brain (17, 18), the question remains, *is* imprinted gene expression actually enriched in the brain compared to other adult tissues? To date this has never been formally tested.

A role for imprinted genes in the brain was initially suggested by (19), and neurological phenotypes observed in, early imprinted gene mouse models (20). In addition, behavioural deficits were seen in imprinting disorders such as Prader-Willi and Angelman syndromes (21, 22). Subsequent studies have revealed diverse roles for imprinted genes in the brain. During development, several imprinted genes are involved in the processes of neural differentiation, migration, axonal outgrowth and apoptosis (23). In the adult brain, studies of mice carrying manipulations of individual imprinted genes have suggested a wide range of behavioural roles including maternal care (24), feeding (25), social behaviour (26, 27), learning/memory (28), cognition (29, 30), and more recently, sleep and circadian activity (31).

In addition to studies on individual imprinted genes, there are a limited number of studies that take a systems level approach to characterizing the role of genomic imprinting in the brain. Early studies examining developing and adult chimeras of normal and parthenogenetic/gynogenetic (Pg/Gg - two maternal genomes) or androgenetic (Ag - two paternal genomes) cells indicated distinct regional distribution for maternally (cortex and hippocampus) and paternally (hypothalamus) expressed genes (20, 32). More recently, Gregg, Zhang (16) used the known imprinting status of 45 imprinted genes and the Allen Brain Atlas to track dichotomous expression of imprinted genes across 118 brain regions to identify brain-wide patterns of expression. Most imprinted genes were expressed in every brain region, but detectable expression of the largest number of imprinted genes was found in regions of the hypothalamus (medial preoptic area, arcuate nucleus), central amygdala, basal nuclei of the stria terminalis and the monoaminergic nuclei, suggesting some form of specialisation. Although pioneering, this study, and others identifying novel imprinted genes and/or mapping allelic expression in the brain (14, 15, 33, 34), did not test whether the expression of these genes was especially enriched in given brain regions but simply asked if they were expressed, at any level, or not.

Here we address the question of whether the brain and/or specific brain circuitry is a foci for genomic imprinting by exploiting the rapidly expanding number of single-cell RNA sequencing (scRNA-seq) datasets and systematically investigating imprinted gene enrichment and over-representation in the murine brain. We performed this by a hierarchical sequence of data analysis, using datasets that allowed a multi-organ (Level 1) comparison first, before proceeding to brain-specific (Level 2) and brain region-specific (Level 3) comparisons with the outcome of each level informing the data selection for the next one, to identify a consistent pattern of enrichment (Figure 1). We sought to provide a robust assessment of the neural systems on which imprinted genes converge, statistically validating previous assumptions, identifying neuronal domains that have received less emphasis in earlier studies, and providing testable predictions for novel neural and behavioural phenotypes associated with specific genes and imprinted gene networks.

**Figure 1.**
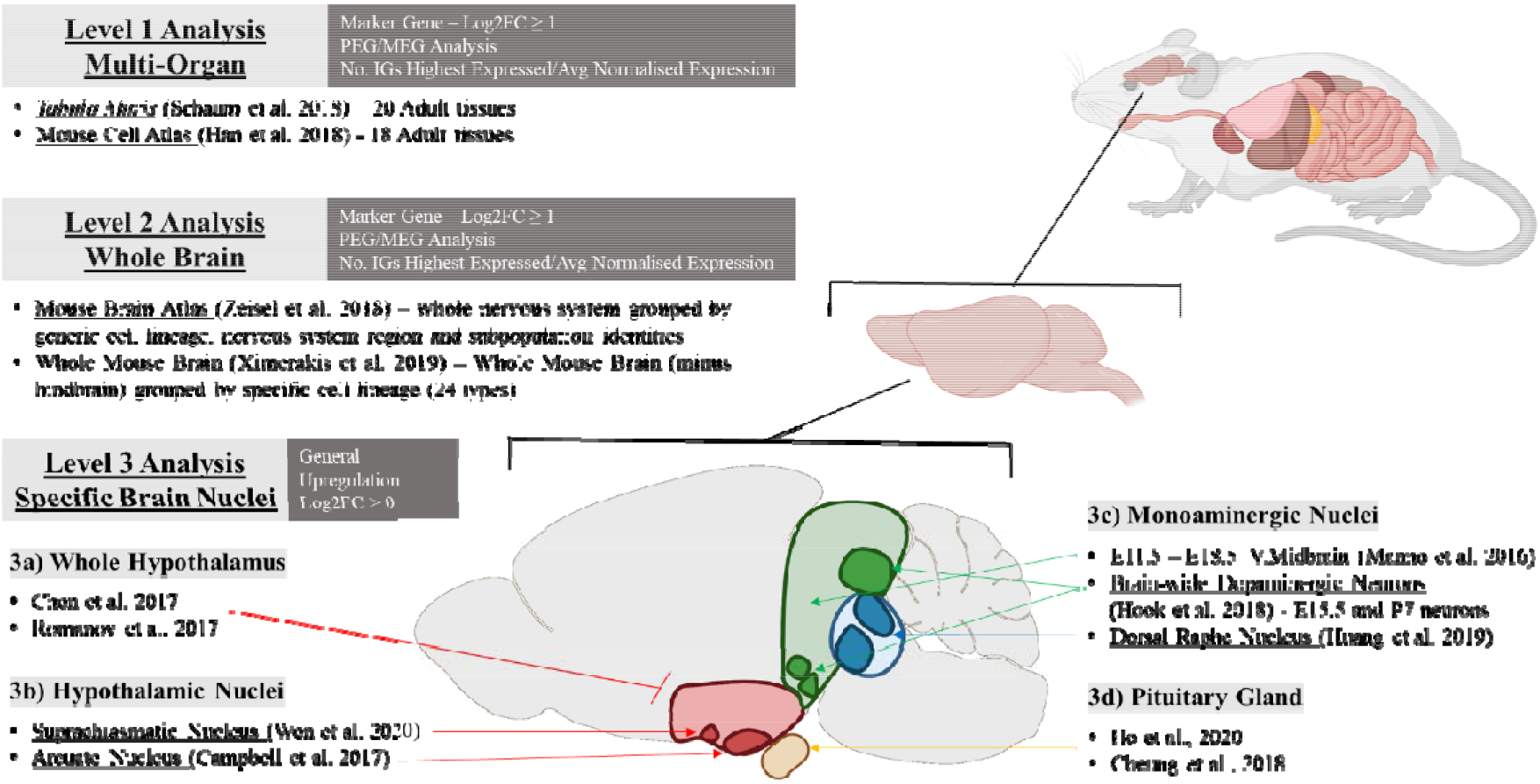
The hierarchical set of datasets in this analysis. The datasets are sorted into Level 1 (Multi-Organ), Level 2 (Whole Brain) and Level 3 (Specific Brain Nuclei) analyses. The original publication and specific tissue/s analysed are provided for each analysis. White text in dark grey box indicates specifics to the analysis at that level – whether the analysis used the ‘marker gene’ Log2FC criteria or the relaxed Log2FC > 0 criterion, whether paternally and maternally expressed gene (PEG/MEG) analysis was carried out and whether the number of IGs with highest expression in a cell population and the average normalised expression were reported for imprinted genes.

## RESULTS

### Imprinted gene expression is enriched in the brain in a multi-organ analysis (Level 1 Analysis)

The Mouse Cell Atlas (MCA) (35) and the *Tabula Muris* (TM) (36) are single cell compendiums containing ~20 overlapping, but not identical, adult mouse organs. Key overlapping organs include the bladder, brain, kidney, lung, limb muscle, and pancreas while organs included in only one dataset include the ovary, testes, uterus, stomach within the MCA, and the heart, fat, skin, trachea and diaphragm within the TM. These compendiums create a snapshot of gene expression across adult tissues to assess imprinted gene enrichment. Since this study focused on the adult body and brain, fetal tissues (including the placenta) were not assessed.

An over-representation analysis (ORA) was performed on both datasets. All data were processed according to the original published procedure, a list of upregulated genes was produced for each tissue/identity group (vs. all other tissue/identity groups) and a one-sided Fisher’s Exact test was performed using a custom list of imprinted genes (Supplemental Table S1) to identify tissues in which imprinted genes were over-represented amongst the upregulated genes for that tissue. Each dataset in this study was analysed independently which allowed us to look for convergent patterns of enrichment between datasets of similar tissues/cell-types. Across only adult tissues, imprinted genes were convergently over-represented in the pancreas, bladder and the brain in both datasets (Figure 2A). In addition, in the MCA adult tissue dataset, there was a significant over-representation in the uterus (Table 1), and in the *Tabula Muris* analysis (Table 2), there was a significant over-representation in the muscle-based tissues - diaphragm, trachea, and limb muscles. In addition to the ORA, to identify situations in which imprinted genes were in fact enriched amongst the stronger markers of a tissue/cell-type, we performed a Gene-Set Enrichment Analysis (GSEA) on tissues meeting minimum criteria (see Methods), which assessed whether imprinted genes were enriched within the top ranked upregulated genes for that tissue (ranked by Log2 Fold Change). No tissue at this level showed a significant GSEA for imprinted genes. Mean normalised expression of imprinted genes across identity groups (Supplemental Table S2) was the highest for Brain in the MCA and highest for Pancreas in the TM (Brain (Non-Myeloid) was the fourth highest).

**Table 1.** Imprinted gene over-representation in MCA adult tissues (35). *Identity –* Tissue identities for the cells used in analysis; *Up Reg* – number of upregulated genes with *q* ≤ 0.05 and Log2FC ≥ 1 (total number of genes in the dataset in brackets); *IG* – number of imprinted genes upregulated with *q* ≤ 0.05 and Log2FC ≥ 1 (total number of IGs in the dataset in brackets); *ORA p* – *p* value from over representation analysis on groups with minimum 5% of total IGs; *ORA q* – Bonferroni corrected *p* value from ORA; *Mean FC IG* – mean fold change for upregulated imprinted genes; *Mean FC Rest* – mean fold change for all other upregulated genes; *No. IGs with highest expression* – Number of IGs with highest mean expression for cells from that identity group.

| Tissue Identity | Up Reg (20,534) | IG (95) | ORA *p* | ORA *q* | Mean FC IG | Mean FC Rest | No. IGs with highest expression |
| --- | --- | --- | --- | --- | --- | --- | --- |
| Pancreas | 2737 | 42 | 1.57E-13 | **1.89E-12** | 8.74 | 10.32 | 22 |
| Brain | 3401 | 34 | 4.43E-06 | **5.31E-05** | 8.76 | 125.00 | 19 |
| Bladder | 3183 | 29 | 0.000168 | **0.002012** | 4.45 | 8.51 | 8 |
| Uterus | 2567 | 22 | 0.002827 | **0.033919** | 4.66 | 8.46 | 7 |
| Lung | 1203 | 8 | 0.192705 | 1 | 3.82 | 151.41 | 4 |
| Ovary | 2219 | 13 | 0.223666 | 1 | 7.46 | 11.27 | 5 |
| Kidney | 1714 | 10 | 0.268425 | 1 | 13.76 | 182.89 | 5 |
| Liver | 1739 | 8 | 0.560145 | 1 | 4.55 | 80.51 | 3 |
| Stomach | 1821 | 7 | 0.748590 | 1 | 4.24 | 88.60 | 3 |
| Thymus | 1805 | 6 | 0.851579 | 1 | 2.78 | 6.76 | 2 |
| Small Intestine | 1719 | 5 | 0.908008 | 1 | 7.99 | 218.64 | 2 |
| Testis | 5212 | 14 | 0.995891 | 1 | 27.04 | 5058.36 | 10 |
| Bone Marrow | 1095 | 2 | - | - | 5.31 | 4.43 | 1 |
| Mammary Gland Virgin | 902 | 4 | - | - | 3.70 | 4.03 | 0 |
| Muscle | 1127 | 4 | - | - | 8.64 | 15.05 | 3 |
| Peripheral Blood | 1146 | 3 | - | - | 3.78 | 3.57 | 0 |
| Prostate | 369 | 0 | - | - | 0.00 | 478.10 | 0 |
| Spleen | 1501 | 1 | - | - | 4.90 | 4.77 | 1 |

**Table 2.** Imprinted gene over-representation in *Tabula Muris* adult tissues (36). *GSEA p* – *p* value from Gene Set Enrichment Analysis for identity groups with 15+ IGs and Mean FC IG > Mean FC Rest; *GSEA q* – Bonferroni corrected *p* values from GSEA. All other column descriptions can be found in the legend of Table 1.

| Tissue Identity | Up Reg (20,839) | IG (107) | ORA *p* | ORA *q* | Mean FC IG | Mean FC Rest | GSEA *p* | GSEA *q* | No. IGs with highest expression |
| --- | --- | --- | --- | --- | --- | --- | --- | --- | --- |
| Diaphragm | 416 | 19 | 3.66E-13 | **4.75E-12** | 6.49 | 4.83 | 0.1898 | 0.3796 | 4 |
| Limb Muscle | 761 | 24 | 6.32E-13 | **8.22E-12** | 9.02 | 5.09 | 0.0552 | 0.1104 | 8 |
| Pancreas | 4104 | 43 | 8.31E-07 | **1.08E-05** | 12.52 | 12.60 | - | - | 29 |
| Trachea | 1979 | 25 | 1.78E-05 | **0.0002** | 3.81 | 4.57 | - | - | 5 |
| Brain (Non-Myeloid) | 3081 | 31 | 0.0001 | **0.0016** | 12.16 | 14.17 | - | - | 14 |
| Bladder | 3338 | 31 | 0.0005 | **0.0068** | 3.30 | 5.30 | - | - | 16 |
| Fat | 1263 | 12 | 0.0286 | 0.3713 | 3.46 | 3.68 | - | - | 1 |
| Heart | 1108 | 10 | 0.0585 | 0.7601 | 2.87 | 5.14 | - | - | 0 |
| Mammary Gland | 1826 | 12 | 0.2264 | 1 | 3.52 | 5.24 | - | - | 3 |
| Liver | 1808 | 7 | 0.8307 | 1 | 6.19 | 54.93 | - | - | 3 |
| Aorta | 3515 | 14 | 0.8832 | 1 | 7.47 | 16.08 | - | - | 2 |
| Tongue | 4295 | 15 | 0.9696 | 1 | 4.15 | 7.16 | - | - | 8 |
| Large Intestine | 4758 | 11 | 0.9998 | 1 | 5.95 | 12.22 | - | - | 5 |
| Brain (Myeloid) | 1024 | 5 | - | - | 3.39 | 6.80 | - | - | 2 |
| Kidney | 584 | 3 | - | - | 24.94 | 22.90 | - | - | 1 |
| Lung | 914 | 2 | - | - | 2.73 | 5.41 | - | - | 0 |
| Marrow | 1957 | 5 | - | - | 7.65 | 5.25 | - | - | 4 |
| Skin | 1612 | 4 | - | - | 4.15 | 8.36 | - | - | 1 |
| Spleen | 625 | 1 | - | - | 4.63 | 4.28 | - | - | 0 |
| Thymus | 678 | 4 | - | - | 3.46 | 7.45 | - | - | 1 |

**Figure 2.**
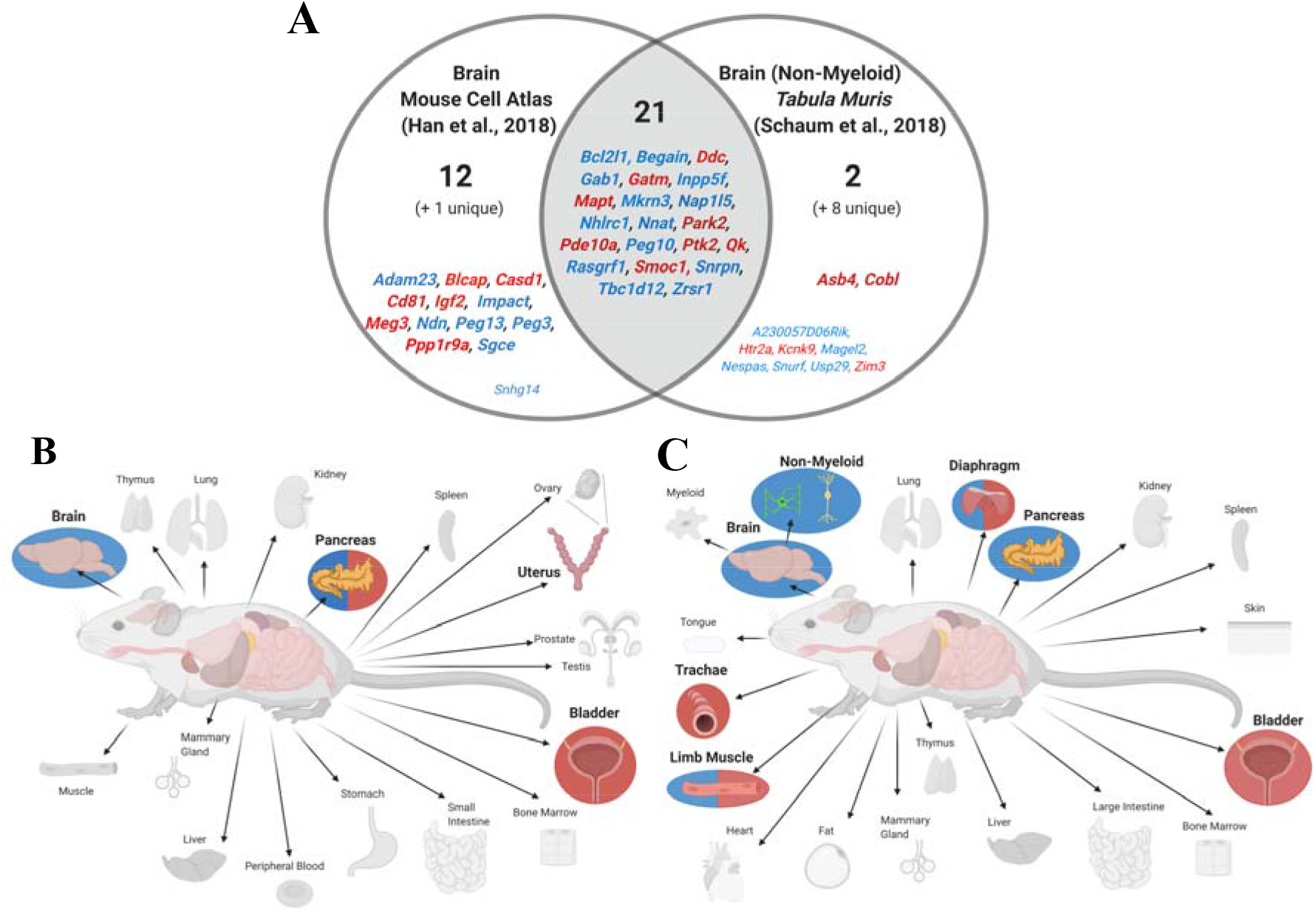
Level 1 multi-organ comparison summary graphics. (*A*) Venn diagram of upregulated imprinted genes in the brain in Mouse Cell Atlas and in the brain (non-myeloid) in the Tabula Muris. Imprinted genes are listed which show significant upregulation (*q* ≤ 0.05 and Log2FC ≥ 1) in the tissues. Although these tissues are not identical, these were the two brain associated over-representations in the enrichment analysis. Parental-bias is indicated by colour (MEG - red, PEG - blue). From the 119 imprinted genes in the gene list, only 92 were common to both analyses (i.e., successfully sequenced and passed gene quality control filters). 34 imprinted genes were upregulated in the brain in the MCA and 31 genes in the *TM*. Genes in common from the two analyses are presented in bold and totalled in each section of the Venn Diagram, while genes found upregulated in one analysis but not available in the other analysis are included in small font and the number indicated in brackets. (*B*) Tissues with over-representation in MCA. Coloured tissues with bold labels were over-represented tissues using all imprinted genes, tissues with a blue circle behind were over-represented for PEGs alone, a red circle represent the same for MEGs, and a red/blue split circle were over-represented for both PEGs and MEGs. (*C*) Tissues with over-representation in *Tabula Muris*. See 2b description for details.

Given the interest in the different functions of maternally expressed genes (MEGs) and paternally expressed genes (PEGs), we additionally ran the large-scale enrichment analyses (Levels 1 and 2) using separate lists of PEGs and MEGs. At Level 1, MEGs and PEGs (Supplemental Table S3A, S3B, S4A and S4B) revealed a similar pattern of enrichment in both datasets (Fig. 2). PEGs were over-represented in the brain in both datasets (MCA - *q* = 4.56×10^−6^, *TM - q* = 0.0005) while MEGs were not. PEGs were also over-represented in the diaphragm (*q* = 0.0007), limb muscle (*q* = 0.0001) and pancreas (MCA - *q* =1.93×10^−5^, *TM - q* = 0.0002), with a significant GSEA in the MCA pancreas (*p* = 0.02, Supplemental Fig. S1). While MEGs were over-represented in the bladder (MCA - *q* = 0.002, *TM* - *q* = 0.020), the pancreas (MCA - *q* = 1.53×10^−7^) and in the three muscular tissues of the *Tabula Muris* (diaphragm - *q* = 2.13×10^−8^, limb muscle - *q* = 2.43×10^−7^, trachea - *q* = 0.004).

### Imprinted gene expression is enriched in neurons and neuroendocrine cells of the brain (Level 2 Analysis)

We next analysed cells from the whole mouse brain (Level 2), firstly using the Ximerakis, Lipnick (6) dataset, in which cells were grouped from the whole mouse brain (minus the hindbrain) into major cell classes according to cell lineage. Imprinted genes were over-represented in neuroendocrine cells and mature neurons (Table 3).

**Table 3.**
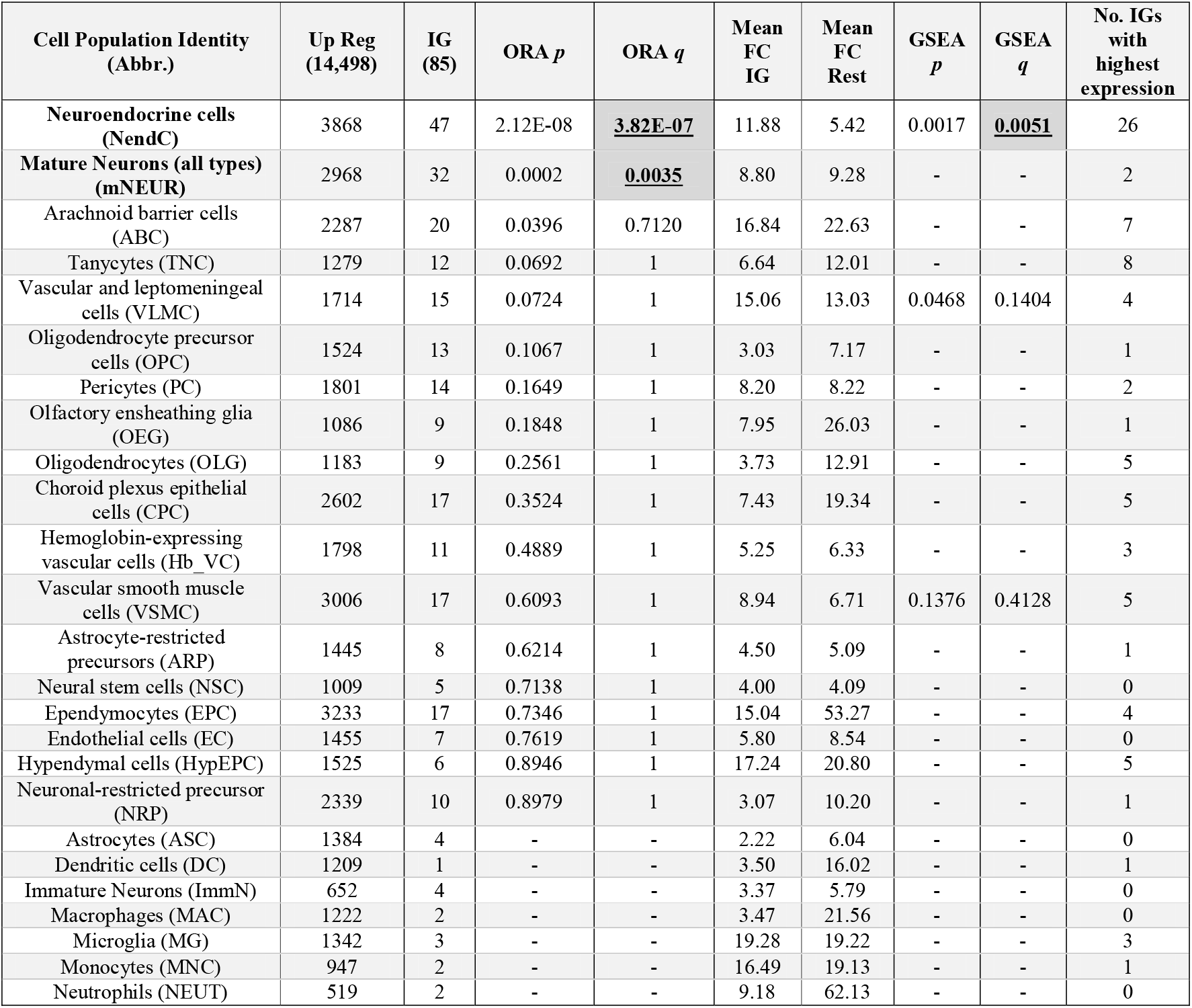
Imprinted gene over-representation in neural lineage types (6). *Identity –* Cell lineage identities for the cells used in analysis. All other column descriptions can be found in the legend of Tables 1 & 2.

| Cell Population Identity (Abbr.) | Up Reg (14,498) | IG (85) | ORA *p* | ORA *q* | Mean FC IG | Mean FC Rest | GSEA *p* | GSEA *q* | No. IGs with highest expression |
| --- | --- | --- | --- | --- | --- | --- | --- | --- | --- |
| Neuroendocrine cells (NendC) | 3868 | 47 | 2.12E-08 | **3.82E-07** | 11.88 | 5.42 | 0.0017 | **0.0051** | 26 |
| Mature Neurons (all types) (mNEUR) | 2968 | 32 | 0.0002 | **0.0035** | 8.80 | 9.28 | - | - | 2 |
| Arachnoid barrier cells (ABC) | 2287 | 20 | 0.0396 | 0.7120 | 16.84 | 22.63 | - | - | 7 |
| Tanycytes (TNC) | 1279 | 12 | 0.0692 | 1 | 6.64 | 12.01 | - | - | 8 |
| Vascular and leptomeningeal cells (VLMC) | 1714 | 15 | 0.0724 | 1 | 15.06 | 13.03 | 0.0468 | 0.1404 | 4 |
| Oligodendrocyte precursor cells (OPC) | 1524 | 13 | 0.1067 | 1 | 3.03 | 7.17 | - | - | 1 |
| Pericytes (PC) | 1801 | 14 | 0.1649 | 1 | 8.20 | 8.22 | - | - | 2 |
| Olfactory ensheathing glia (OEG) | 1086 | 9 | 0.1848 | 1 | 7.95 | 26.03 | - | - | 1 |
| Oligodendrocytes (OLG) | 1183 | 9 | 0.2561 | 1 | 3.73 | 12.91 | - | - | 5 |
| Choroid plexus epithelial cells (CPC) | 2602 | 17 | 0.3524 | 1 | 7.43 | 19.34 | - | - | 5 |
| Hemoglobin-expressing vascular cells (Hb_VC) | 1798 | 11 | 0.4889 | 1 | 5.25 | 6.33 | - | - | 3 |
| Vascular smooth muscle cells (VSMC) | 3006 | 17 | 0.6093 | 1 | 8.94 | 6.71 | 0.1376 | 0.4128 | 5 |
| Astrocyte-restricted precursors (ARP) | 1445 | 8 | 0.6214 | 1 | 4.50 | 5.09 | - | - | 1 |
| Neural stem cells (NSC) | 1009 | 5 | 0.7138 | 1 | 4.00 | 4.09 | - | - | 0 |
| Ependymocytes (EPC) | 3233 | 17 | 0.7346 | 1 | 15.04 | 53.27 | - | - | 4 |
| Endothelial cells (EC) | 1455 | 7 | 0.7619 | 1 | 5.80 | 8.54 | - | - | 0 |
| Hypendymal cells (HypEPC) | 1525 | 6 | 0.8946 | 1 | 17.24 | 20.80 | - | - | 5 |
| Neuronal-restricted precursor (NRP) | 2339 | 10 | 0.8979 | 1 | 3.07 | 10.20 | - | - | 1 |
| Astrocytes (ASC) | 1384 | 4 | - | - | 2.22 | 6.04 | - | - | 0 |
| Dendritic cells (DC) | 1209 | 1 | - | - | 3.50 | 16.02 | - | - | 1 |
| Immature Neurons (ImmN) | 652 | 4 | - | - | 3.37 | 5.79 | - | - | 0 |
| Macrophages (MAC) | 1222 | 2 | - | - | 3.47 | 21.56 | - | - | 0 |
| Microglia (MG) | 1342 | 3 | - | - | 19.28 | 19.22 | - | - | 3 |
| Monocytes (MNC) | 947 | 2 | - | - | 16.49 | 19.13 | - | - | 1 |
| Neutrophils (NEUT) | 519 | 2 | - | - | 9.18 | 62.13 | - | - | 0 |

Neuroendocrine cells were defined as a heterogeneous cluster, containing peptidergic neurons and neurosecretory cells expressing neuronal marker genes (e.g., *Syt1* and *Snap25*) alongside neuropeptide genes (e.g., *Oxt, Avp, Gal, Agrp* and *Sst*) but distinguished by Ximerakis, Lipnick (6) by the unique expression of *Baiap3* which plays an important role in the regulation of exocytosis in neuroendocrine cells (37). GSEA additionally showed that the imprinted genes were enriched in the genes with the highest fold change values for neuroendocrine cells only (Fig. 3). 26 imprinted genes had their highest expression in the neuroendocrine cells and the mean normalised expression of imprinted genes was almost twice as high for neuroendocrine cells as the next highest identity group (Supplemental Table S2). The MEG/PEG analysis (Supplemental Table S5A and S5B) for this dataset found that PEGs were over-represented in mature neurons (*q* = 0.027) and neuroendocrine cells (*q* = 8.97 × 10^−6^). MEGs were also over-represented in neuroendocrine cells (*q* = 0.047) and uniquely over-represented in Arachnoid barrier cells (*q* = 0.014). Only PEGs replicated the significant GSEA in neuroendocrine cells (*p* = 4×10^−4^, Supplemental Fig. S2).

**Figure 3.**
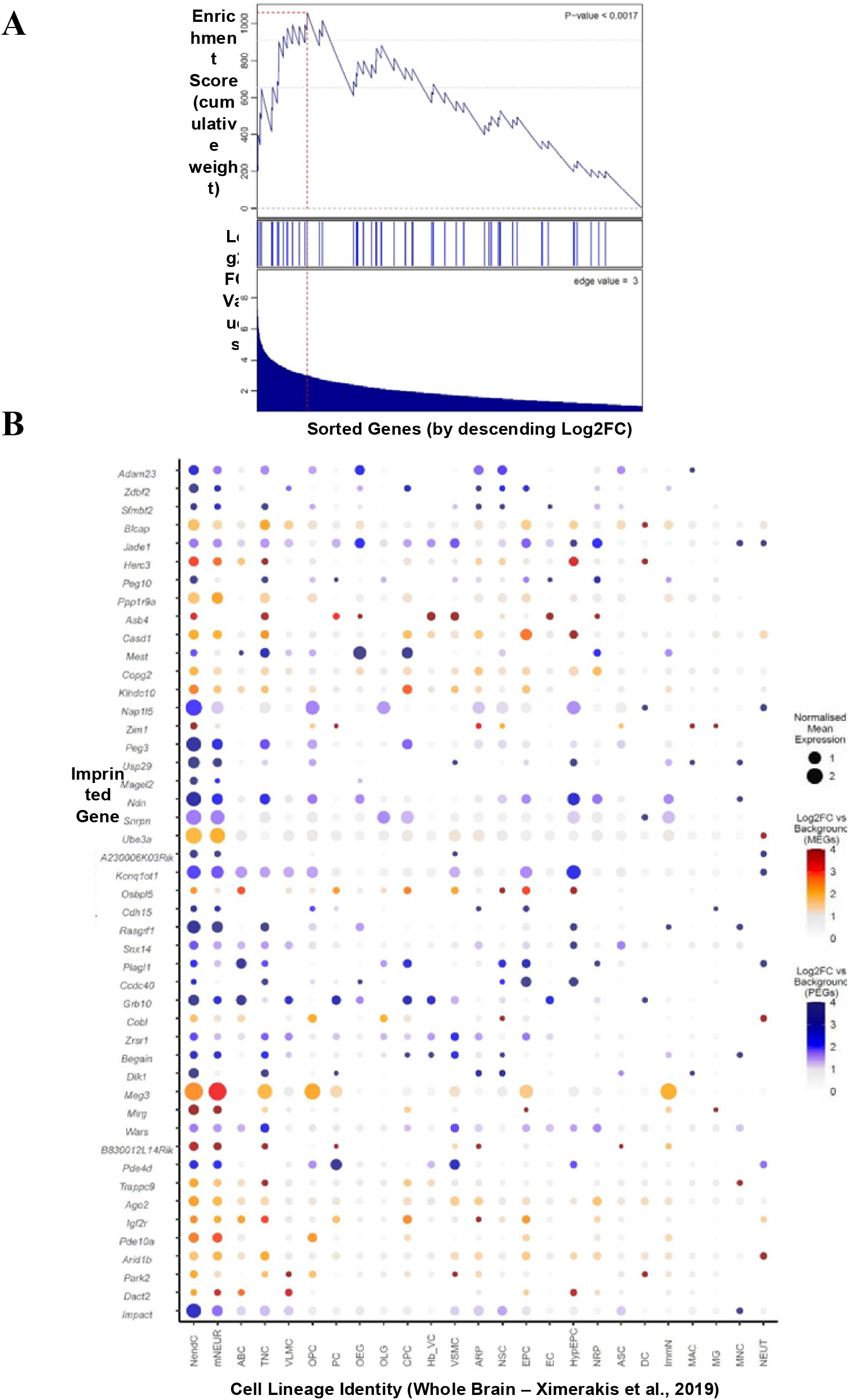
GSEA and dot plots for imprinted genes upregulated in neuroendocrine cells in the Ximerakis, Lipnick (6) whole mouse brain dataset. (*A*) GSEA for imprinted genes upregulated in the neuroendocrine cells. In the analysis, genes are sorted by strength by which they mark this neuronal cluster (sorted by Log2FC values) indicated by the bar (middle). Fold change values are displayed along the bottom of the graph. The genes are arrayed left (strongest marker) to right and blue lines mark where imprinted genes fall on this array. The vertical axis indicates an accumulating weight, progressing from left to right and increasing or decreasing depending on whether the next gene is an imprinted gene or not. The *p*-value represents the probability of observing the maximum value of the score (red dashed line) if the imprinted genes are distributed randomly along the horizontal axis. The *q*-value for this analysis was significant at 0.0036. (*B*) Dot plot of imprinted genes upregulated in the ‘Neuroendocrine cells’ plotted across all identified cell types (Abbr. in Table 3). Imprinted genes were plotted in chromosomal order. Size of points represented absolute mean expression; colour represented the size of the Log2FC value for the cell identity group (e.g., neuroendocrine cells) vs. all other cells. Unique colour scales are used for MEGs (red/orange) and PEGs (blue). Where a gene was not expressed in a cell type, this appears as a blank space in the plot

The second dataset at this level was Zeisel, Hochgerner (1) Mouse Brain Atlas (MBA) which allowed a much deeper investigation of nervous system enrichment with sequencing of the entire murine nervous system and identifying cells by both brain region and cell type. Concordant with the previous findings, primary analysis separating cells by lineage revealed over-representation of imprinted genes in neurons only (Table 4). The overlap between the upregulated imprinted genes for the over-represented neural-lineage cells from the Level 2 datasets are displayed in Figure 4. Additionally, PEGs alone demonstrated no significant over-representations in cell lineage types while MEGs demonstrated over-representation in vascular cells only (*q* = 0.0004) (Supplemental Table S6A and S6B).

**Table 4.** Imprinted gene over-representation in nervous system cell types (1). *Identity –* Cell identities for the cells used in analysis. All other column descriptions can be found in the legend of Table 1.

| Cell Population Identity | Up Reg (19,547) | IG (109) | ORA *p* | ORA *q* | Mean FC IG | Mean FC Rest | No. IGs with highest expression |
| --- | --- | --- | --- | --- | --- | --- | --- |
| Neurons | 5710 | 44 | 0.0081 | **0.0487** | 11.73 | 24.97 | 45 |
| Vascular | 2473 | 22 | 0.0171 | 0.1029 | 17.91 | 26.64 | 16 |
| Oligos | 1587 | 11 | 0.2701 | 1 | 4.64 | 11.48 | 12 |
| Peripheral Glia | 2820 | 16 | 0.5117 | 1 | 5.42 | 12.64 | 12 |
| Ependymal | 3683 | 20 | 0.5912 | 1 | 24.52 | 66.97 | 15 |
| Immune | 1564 | 7 | 0.7787 | 1 | 13.42 | 93.05 | 5 |
| Astrocytes | 1539 | 4 | - | - | 2.88 | 10.73 | 3 |

**Figure 4.**
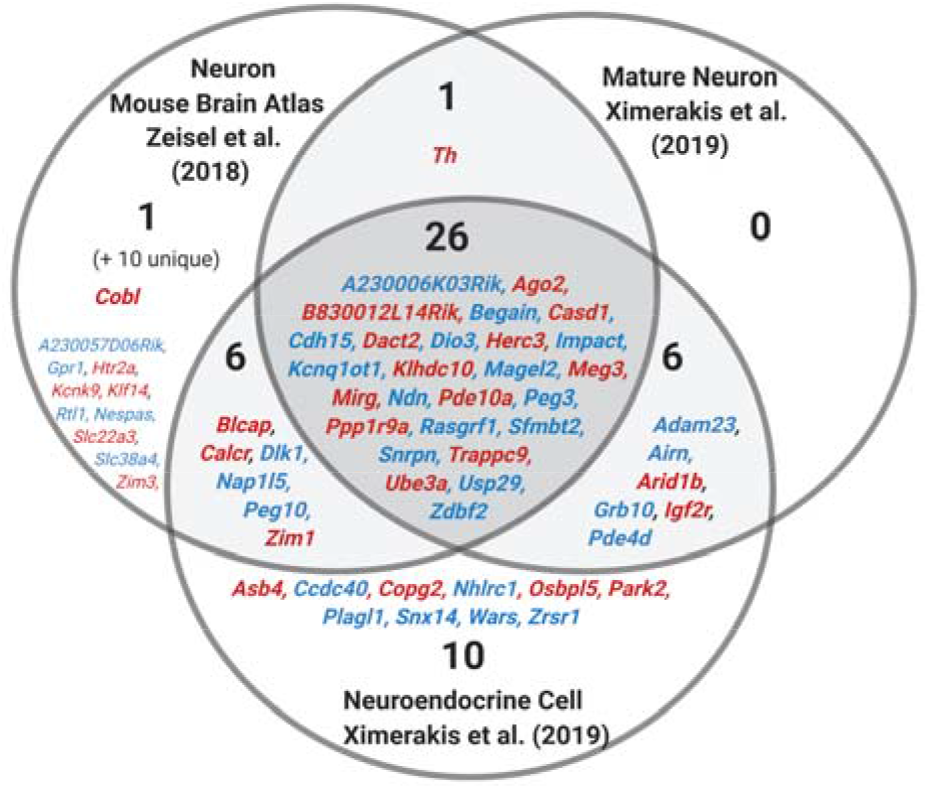
Venn diagram of upregulated imprinted genes in the mature neuronal cells in the whole brain datasets of Zeisel, Hochgerner (1) and Ximerakis, Lipnick (6). Imprinted genes are listed which show significant upregulation (*q* ≤ 0.05 and Log2FC ≥ 1) in the cells. Although these cell types are not identical, these were all mature neural lineage cells with over-representations in the enrichment analysis. Parental-bias is indicated by colour (MEG - red, PEG - blue. From the 119 imprinted genes in the gene list, only 88 were common to both analyses (i.e., successfully sequenced and passed gene quality control filters). 45 imprinted genes were upregulated in neurons in the MBA, and in Ximerakis, Lipnick (6), 33 imprinted genes were upregulated in neurons and 48 genes in neuroendocrine cells. Genes in common from the two analyses are presented in bold and totalled in each section of the Venn Diagram, while genes found upregulated in one analysis but not available in the other analysis are included in small font and the number indicated in brackets.

### The hypothalamus, ventral midbrain, pons and medulla are enriched for imprinted gene expression (Level 2 Analysis)

After confirming neuron-specific enrichment of imprinted genes in the MBA dataset, further MBA analysis was performed on cells classified as neurons and then grouped by brain/nervous system regions. Significant over-representation was seen in neurons of the hypothalamus, ventral midbrain, medulla, and pons (Table 5). The pons and medulla had the largest number, 45 and 44 respectively, of imprinted genes upregulated (Figure 5A).

**Table 5.**
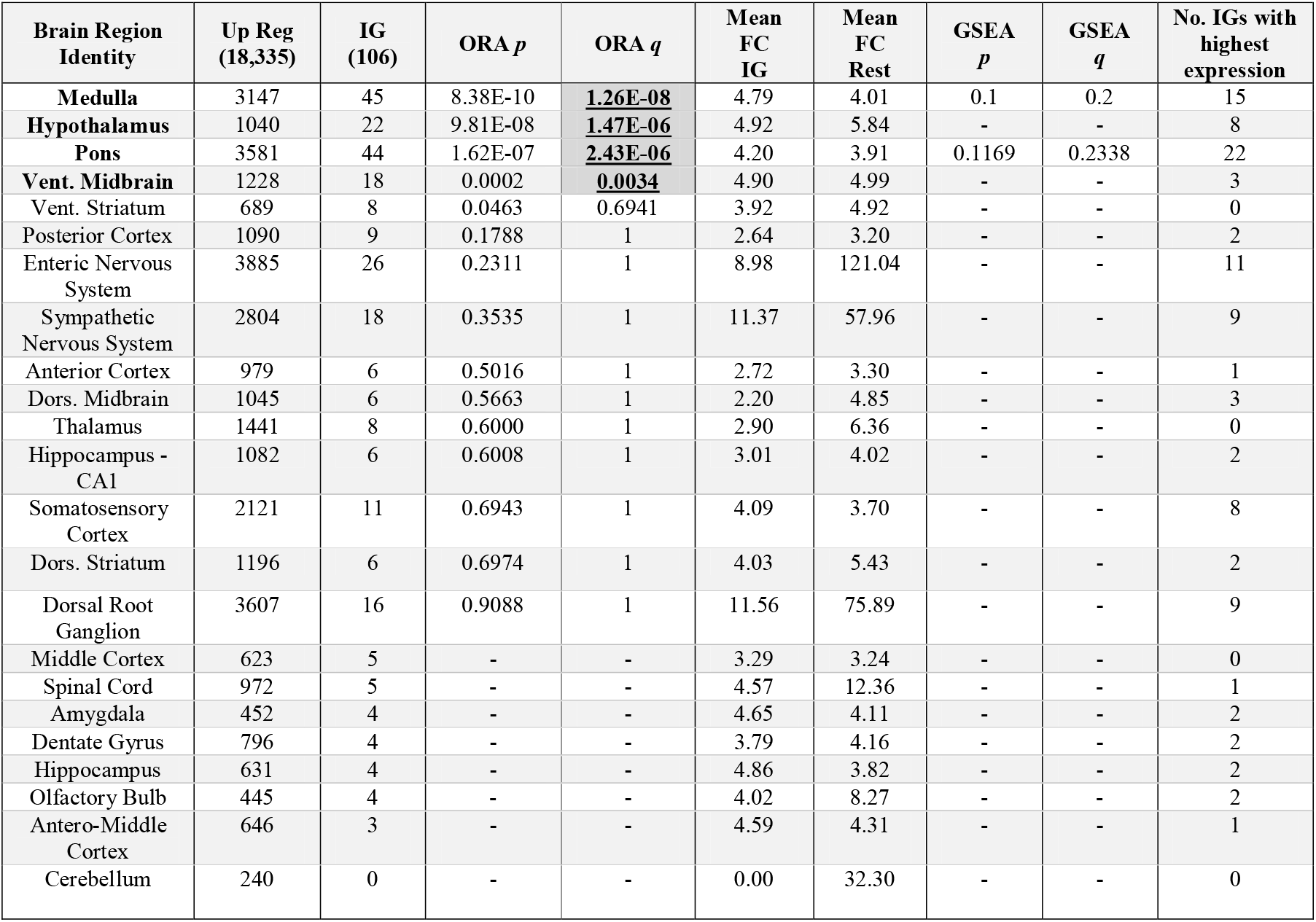
Imprinted gene over-representation in nervous system region(1). *Identity –* Nervous system regional identities for the cells used in analysis genes. All other column descriptions can be found in the legend of Tables 1 & 2.

| Brain Region Identity | Up Reg (18,335) | IG (106) | ORA $p$ | ORA $q$ | Mean FC IG | Mean FC Rest | GSEA $p$ | GSEA $q$ | No. IGs with highest expression |
| --- | --- | --- | --- | --- | --- | --- | --- | --- | --- |
| Medulla | 3147 | 45 | 8.38E-10 | **1.26E-08** | 4.79 | 4.01 | 0.1 | 0.2 | 15 |
| Hypothalamus | 1040 | 22 | 9.81E-08 | **1.47E-06** | 4.92 | 5.84 | - | - | 8 |
| Pons | 3581 | 44 | 1.62E-07 | **2.43E-06** | 4.20 | 3.91 | 0.1169 | 0.2338 | 22 |
| Vent. Midbrain | 1228 | 18 | 0.0002 | **0.0034** | 4.90 | 4.99 | - | - | 3 |
| Vent. Striatum | 689 | 8 | 0.0463 | 0.6941 | 3.92 | 4.92 | - | - | 0 |
| Posterior Cortex | 1090 | 9 | 0.1788 | 1 | 2.64 | 3.20 | - | - | 2 |
| Enteric Nervous System | 3885 | 26 | 0.2311 | 1 | 8.98 | 121.04 | - | - | 11 |
| Sympathetic Nervous System | 2804 | 18 | 0.3535 | 1 | 11.37 | 57.96 | - | - | 9 |
| Anterior Cortex | 979 | 6 | 0.5016 | 1 | 2.72 | 3.30 | - | - | 1 |
| Dors. Midbrain | 1045 | 6 | 0.5663 | 1 | 2.20 | 4.85 | - | - | 3 |
| Thalamus | 1441 | 8 | 0.6000 | 1 | 2.90 | 6.36 | - | - | 0 |
| Hippocampus - CA1 | 1082 | 6 | 0.6008 | 1 | 3.01 | 4.02 | - | - | 2 |
| Somatosensory Cortex | 2121 | 11 | 0.6943 | 1 | 4.09 | 3.70 | - | - | 8 |
| Dors. Striatum | 1196 | 6 | 0.6974 | 1 | 4.03 | 5.43 | - | - | 2 |
| Dorsal Root Ganglion | 3607 | 16 | 0.9088 | 1 | 11.56 | 75.89 | - | - | 9 |
| Middle Cortex | 623 | 5 | - | - | 3.29 | 3.24 | - | - | 0 |
| Spinal Cord | 972 | 5 | - | - | 4.57 | 12.36 | - | - | 1 |
| Amygdala | 452 | 4 | - | - | 4.65 | 4.11 | - | - | 2 |
| Dentate Gyrus | 796 | 4 | - | - | 3.79 | 4.16 | - | - | 2 |
| Hippocampus | 631 | 4 | - | - | 4.86 | 3.82 | - | - | 2 |
| Olfactory Bulb | 445 | 4 | - | - | 4.02 | 8.27 | - | - | 2 |
| Antero-Middle Cortex | 646 | 3 | - | - | 4.59 | 4.31 | - | - | 1 |
| Cerebellum | 240 | 0 | - | - | 0.00 | 32.30 | - | - | 0 |

**Figure 5.**
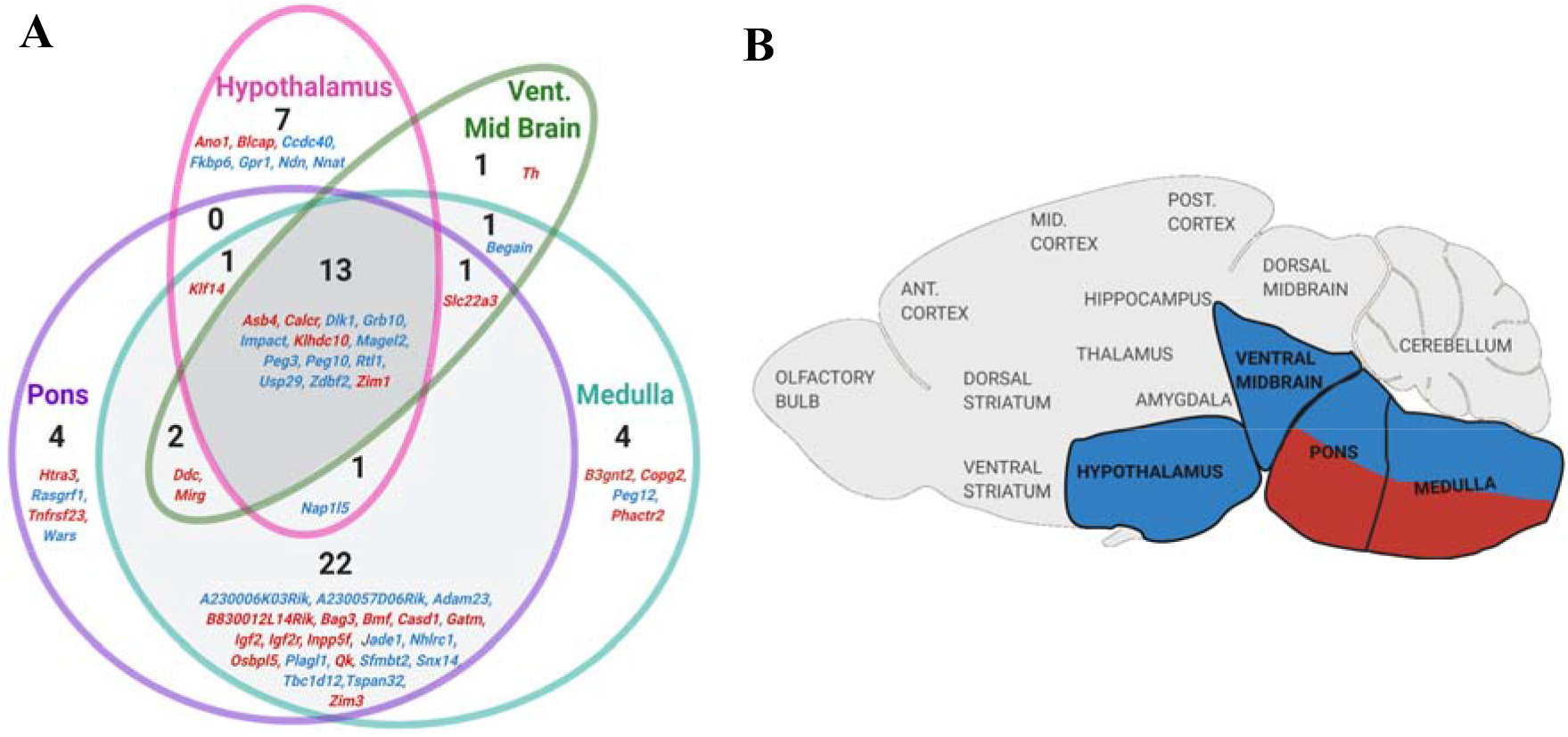
Level 2 Brain Region Analysis summary figures. (*A*) Venn diagram of upregulated imprinted genes in the neurons of enriched nervous system regions from the Mouse Brain Atlas (1). Imprinted genes are listed which show significant upregulation (*q* ≤ 0.05 and Log2FC ≥ 1) in the regions specified. The number of imprinted genes in each region of the Venn diagram are specified. Parental-bias of imprinted genes is indicated by colour (MEG - red, PEG - blue). (*B*) Brain regions enriched for imprinted gene expression via ORA or GSEA in the MBA (1). Regions over-represented for all imprinted genes are bolded. Regions over-represented for PEG expression alone are coloured blue while regions enriched for MEG expression alone are coloured red.

Regional analysis for MEGs and PEGs separately (Supplemental Table S7A and S7B), revealed that PEGs were over-represented in hypothalamus (*q* = 6.53×10^−7^), ventral midbrain (*q* = 0.018), the pons (*q* = 4.65×10^−5^) and the medulla (*q* = 4.10×10^−6^); while MEGs were only over-represented in the medulla (*q* = 0.002) but had a significant GSEA for the pons (*q* = 0.027, Supplemental Fig. S3); see Figure 5B. Neurons were then recategorized into unique subpopulations identified by marker genes (1) to uncover the specific neural populations underlying the enrichment seen in the hypothalamus, pons and medulla, and midbrain (Fig. 6; Supplemental Table S8). Each neural population was identified by its distinct gene expression and suspected location within the brain (see http://mousebrain.org/ for an online resource with detailed information on each cluster).

**Figure 6.**
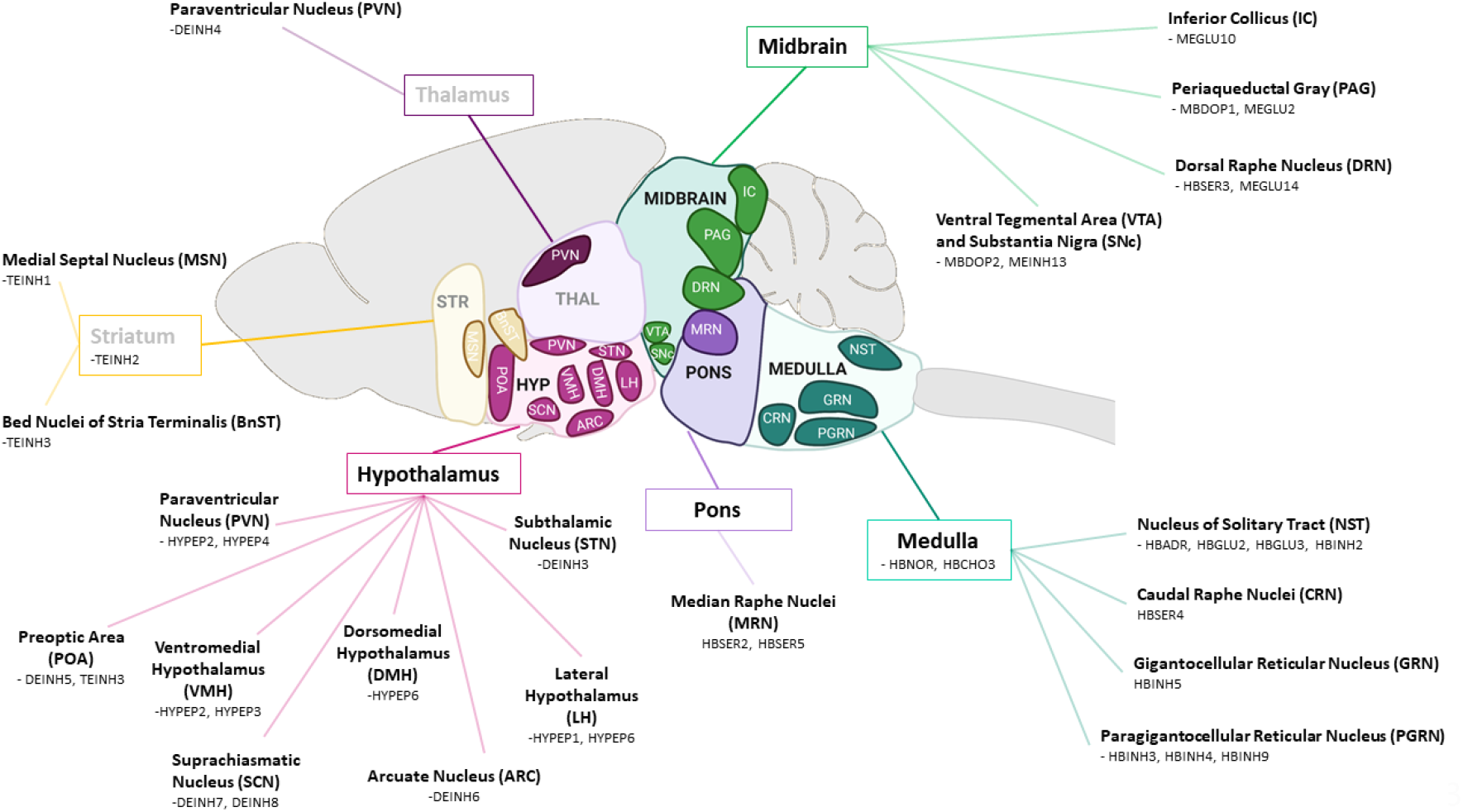
Anatomical labelling of all the neural subpopulations with a significant over-representation of imprinted genes (*q* ≤ 0.05 and Log2FC ≥ 1) in the Mouse Brain Atlas (1). The predicted brain nuclei localisation of the 32 neuronal subpopulations (out of 214 populations identified across the nervous system) specified in the MBA and enriched for imprinted genes. Brain regions that were not found to be enriched for imprinted genes are greyed out. The full Enrichment Analysis is available in Supplemental Table S8.

The hypothalamus was represented by a selection of inhibitory and peptidergic neurons. Inhibitory neurons with over-representation of imprinted genes included: a Subthalamic Nucleus population (notable genes *Lhx8, Gabrq*), two Preoptic Area/ BNST populations (*Nts, Dlk1* / *Gal, Irs4*) representing, an Arcuate nucleus population (*Agrp, Otp*), and two Suprachiasmatic nucleus populations (*Avp, Nms, Six6, Vip*). For peptidergic neurons, over-representation was seen in a ventromedial population (*Gpr101, Tac1, Baiap3*), a ventromedial/paraventricular population (*Otp, Trh, Ucn3*), a lateral hypothalamic population (*Trh, Otp, Ngb*), an oxytocin magnocellular population of the paraventricular and supraoptic nuclei (*Oxt, Otp*), and an orexin producing population of the dorsomedial/lateral hypothalamus (*Hcrt, Pdyn, Trhr*).

The midbrain, medulla and pons were represented by a number of cell groups, with over-representation seen in the medulla-based adrenergic (HBAR) and noradrenergic (HBNOR) groups and the dopaminergic neurons of the midbrain in the Periaqueductal Gray (PAG) (MBDOP1) and the Ventral Tegmental Area (VTA)/Substantia Nigra (SNc) (MBDOP2). Th ere were also several inhibitory (MEINH, HBIN) and excitatory neuron (MEGLU, HBGLU) types spread across the nuclei from the three regions (Fig. 6). The serotonergic populations of the raphe nuclei of these regions (HBSER) were particularly prominent since the pons and medulla-based serotonin neuron populations (HBSER2, HBSER4 and HBSER5) were the only neuron subpopulations out of the 214 total to have a significant GSEA for imprinted genes after correction (Supplemental Fig. S4).

Additional regions of over-representation included neurons in the pallidum and striatum and PVN neurons from the thalamus. In this comparison of 214 neuron populations, no neurons from areas such as the cortex, cerebellum or peripheral nervous system were enriched, and neither were they over-represented in the previous regional analysis. Hence, further analysis focused on those brain regions enriched in this whole brain level analysis.

### Imprinted gene expression is over-represented in specific hypothalamic neuron subtypes (Level 3A&3B Analysis)

We next sought to investigate whether those regional neuron enrichments found within the whole brain comparisons would be further clarified with enriched expression in specific neuronal subpopulations within those regions. Namely we sought to identify neural populations enriched across the whole hypothalamus and those enriched within specific hypothalamic nuclei, and also whether imprinted gene expression was enriched in the other key subpopulation identified, the ventral midbrain and hindbrain dopaminergic and serotonergic populations. Two datasets with single cell sequencing data for the adult hypothalamus existed (3, 4). Both clustered their data into neuronal subpopulations allowing us to look for convergent imprinted enrichment across major hypothalamic neuronal subtypes (Level 3A). Analysis revealed a clear neuronal bias in expression of imprinted genes (Supplemental Table S9A and S10A). Within the Romanov, Zeisel (3) data, there was a significant over-representation of imprinted genes in neurons (*q* = 0.02) and a similar observation was seen in the Chen, Wu (4) data (*q* = 0.001), and both also demonstrated a significant GSEA in neurons (Fig. 7A-D, Romanov, Zeisel (3) – *p* = 0.011, Chen, Wu (4) - *p* = 0.022).

**Figure 7.**
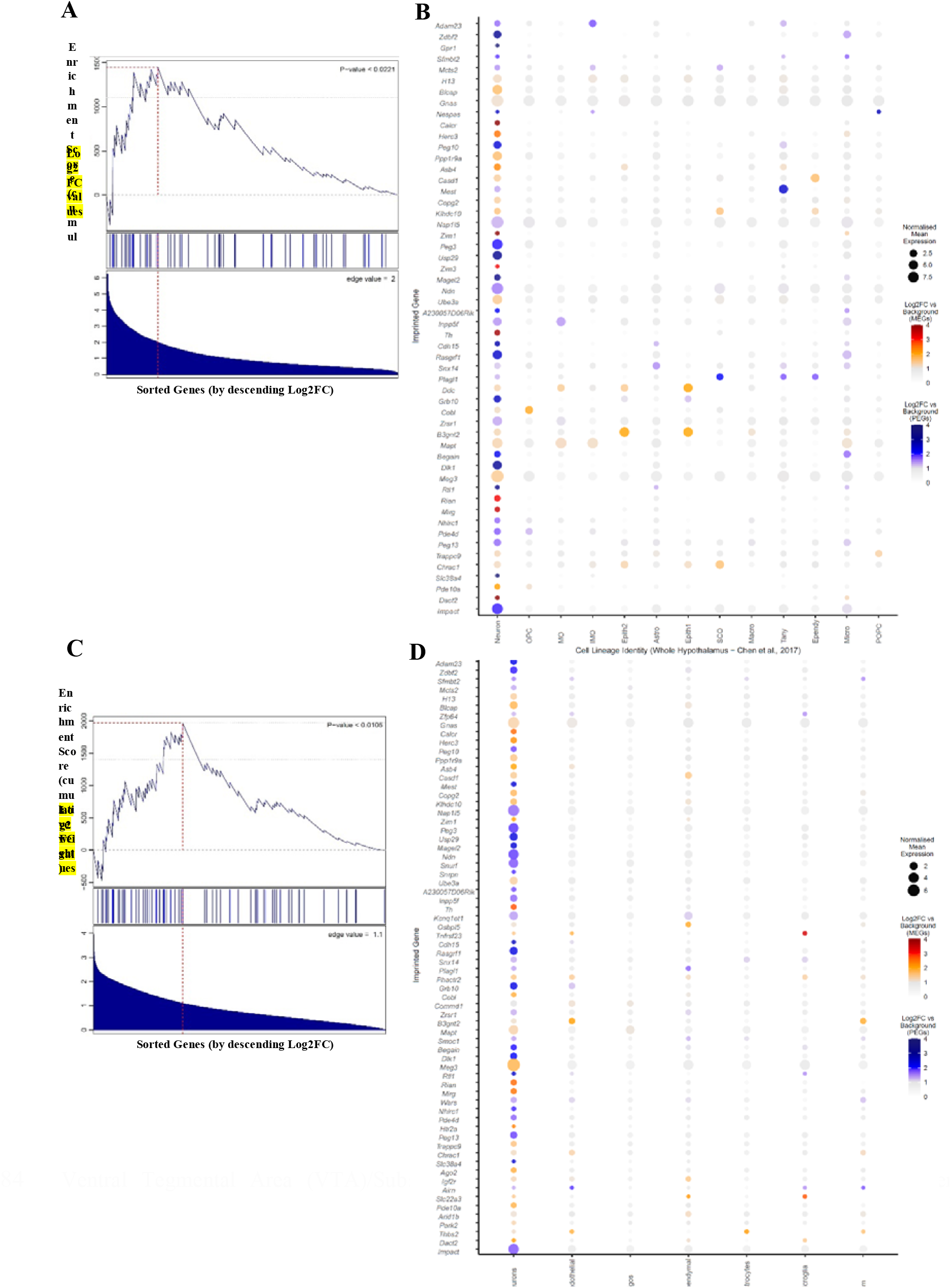
GSEA and Dot plots for imprinted genes upregulated in neurons across the whole hypothalamus. (*A*) GSEA for imprinted genes upregulated in the ‘Neuron’ cell type in the whole hypothalamic dataset of Chen, Wu (4). See legend of Figure 3A for a description of how to interpret the plot. (*B*) Dot plot of imprinted genes upregulated in the ‘Neuron’ cell type plotted across all identified cell types in the Chen, Wu (4) whole hypothalamic dataset. See legend of Figure 3B for a description of how to interpret the plot. Abbr: OPC = Oligodendrocyte Precursor Cell, MG = Myelinating Oligodendrocyte, IMG = Immature Oligodendrocyte, Astro = Astrocyte, Epith = Epithelial, Macro = Macrophage, Tany = Tanycyte, Ependy = Ependymocyte, Micro = Microglia, POPC = Proliferating Oligodendrocyte Progenitor Cell. (*C*) GSEA for imprinted genes upregulated in ‘neurons’ in the whole hypothalamic dataset of Romanov, Zeisel (3). See legend of Figure 3A for a description of how to interpret the plot. (*D*) Dot plot of imprinted genes upregulated in ‘neurons’ plotted across all identified cell types in the Romanov, Zeisel (3)whole hypothalamic dataset. See legend of Figure 3B for a description of how to interpret the plot.

Within the Chen, Wu (4) dataset, 4/33 hypothalamic neuronal subtypes had a significant over-representation of imprinted genes (Supplemental Table S9B). The four subtypes were all GABAergic neurons, specifically: Galanin neurons (*Slc18a2/Gal)* present in a several hypothalamic regions (*q* = 0.0079); a dopaminergic neuron type (*Slc6a3*) with high expression of *Th* and *Prlr* suspected to be the TIDA neurons of the arcuate nucleus (*q* = 0.0001); SCN neurons (*Vipr2)* with very high *Avp* and *Nms* expression (*q* = 0.0071); and *Agrp* feeding promoting neurons of the Arcuate Nucleus (*q* = 0.034). Within the Romanov, Zeisel (3) dataset, 3/62 subtypes had significant over-representation of imprinted gene expression (Supplemental Table S10B): A*grp*/*Npy* neurons (*q* = 0.013), the Arcuate Nucleus feeding neurons also reported in Chen, Wu (4); a *Ghrh/Th* neuronal type (*q* = 0.032), again likely corresponding to neurons from the arcuate nucleus and the top hit (*q* = 1.63×10^−6^) was a poorly segregated population (*Calcr/Lhx1)*, likely due to a deeper inner cluster heterogeneity. This cluster was interesting since the imprinted genes *Calcr* and *Asb4* were amongst its most significant marker genes, and it was notably the only cluster with high expression of all three of *Th, Slc6a3* and *Prlr*. Romanov, Zeisel (3) did not identify any of their populations as the TIDA neurons, but the above pattern of gene expression suggests that this cluster may contain these neurons. Furthermore, the suspected TIDA neurons from the Chen, Wu (4) dataset shared 21/40 upregulated genes of this unresolved cluster (see Supplemental Table S11 for full comparison);

Having consistently found well-known neurons from the arcuate nucleus (*Agrp, Ghrh*), and suprachiasmatic nucleus (*Avp, Vip*) we sought to test imprinted gene enrichment within these hypothalamic regions at a high resolution using datasets sequencing neurons purely from these hypothalamic regions (Level 3B).

### Arcuate nucleus (ARC) (2)

The first nuclei investigated was the ARC sequenced by Campbell, Macosko (2). Imprinted gene over-representation was found in 8/24 arcuate neuron types (Supplemental Table S12). These included the *Agrp*/*Sst* neuron type (with high expression of *Npy, q* = 0.003) and two *Pomc* neuron types (*Pomc/Anxa2, q* = 0.004; *Pomc/Glipr1, q* = 0.03). Pomc expressing neurons are known to work as feeding suppressants (38). Additional significant over-representation was found in the *Ghrh* neuron type (*q* = 0.009), which was also enriched in *Gal* and *Th*. Finally, a highly significant over-representation of imprinted genes was found in the *Th*/*Slc6a3* neuron type (*q* = 1.72 × 10^−8^) identified by the authors as one of the most likely candidates for the TIDA dopaminergic neuron population. Marker genes for this identity group overlapped with the TIDA candidates from the previous two datasets (e.g., *Slc6a3, Th, Lhx1, Calcr*). *Agrp* neurons, *Ghrh* neurons and these TIDA candidate neurons were identified in both whole hypothalamic datasets and at the nuclei level.

### Suprachiasmatic Nucleus (SCN) (5)

Analysis of the 10x chromium data of SCN neurons (Supplemental Table S13) revealed a significant over-representation (*q* = 1.51 × 10^−8^) and GSEA (*p* = 0.004, Supplemental Fig. S5) in the *Avp*/*Nms* neuronal cluster (out of 5 neuronal clusters). This cluster shows the strongest expression for *Oxt, Avp, Avpr1a* and *Prlr* and is one of the three neural group that Wen, Ma (5) found had robust circadian gene expression, and the only subtype with notable phase differences in circadian gene expression in the dorsal SCN. This cluster likely corresponds to the GABA8 cluster found enriched in the Chen, Wu (4) dataset. Figure 8 presents the overlapping upregulated imprinted genes from the convergently upregulated neuron subtypes in the hypothalamic analysis of Level 3a and 3b.

**Figure 8.**
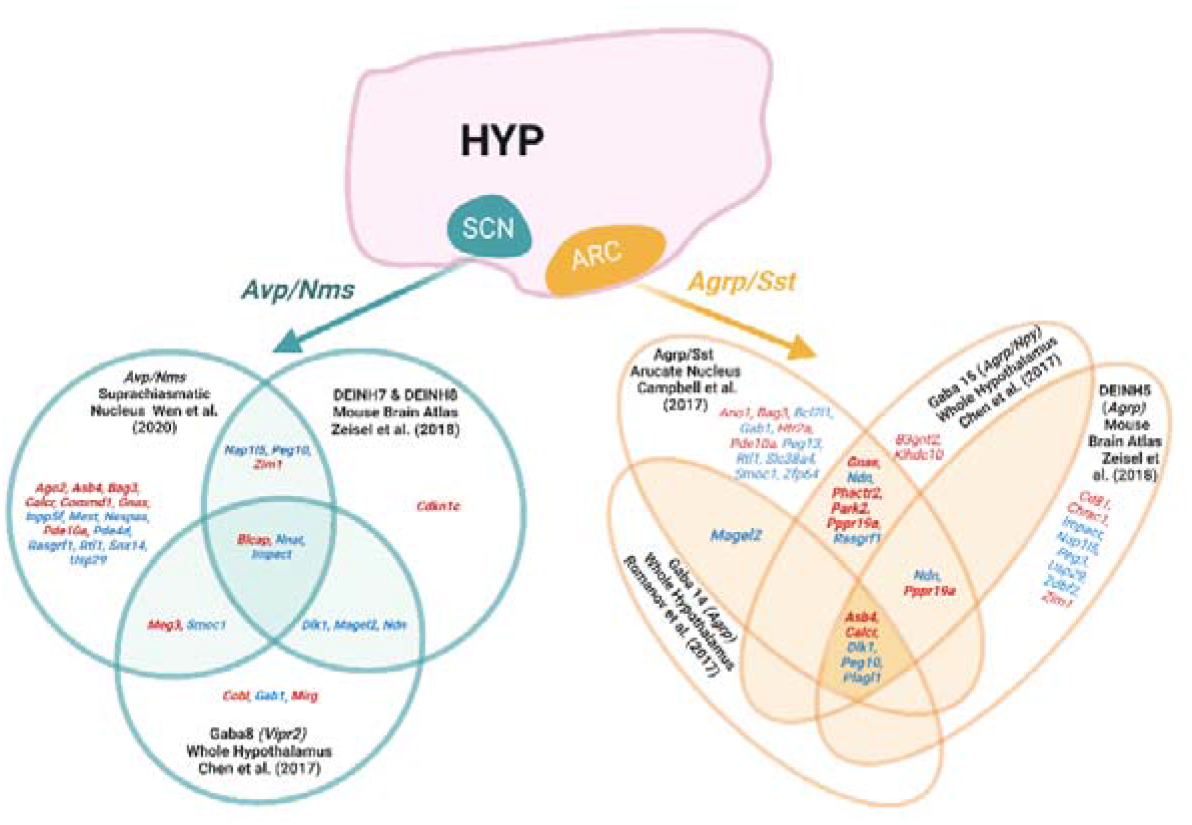
Venn diagrams of upregulated imprinted genes in the neuronal subpopulations from level 3b that were also identified in level 2 and 3a. Imprinted gene overlap was contrasted for *Agrp/Sst* neuronal populations of the Arcuate Nucleus (1–4) and *Avp/Nms* neurons from the Suprachiasmatic Nucleus (1, 4, 5) Imprinted genes are listed which show significant upregulation (*q* ≤ 0.05 and Log2FC > 0) in the subpopulation. Parental-bias is indicated by colour (MEG - red, PEG - blue).

### Imprinted gene expression is over-represented in monoaminergic nuclei of the mid- and hindbrain (Level 3C Analysis)

In the MBA, Whole Hypothalamus and Arcuate Nucleus analyses, dopaminergic clusters were consistently enriched and, to explore this further, analysis of Hook, McClymont (39) data allowed comparison for dopamine neurons across the brain (specifically from the olfactory bulb, arcuate nucleus and midbrain) at two developmental timepoints (E15.5 and Post-natal day (P) 7). The arcuate nucleus P7 dopamine neurons emerged as the clearest over-represented subgroups (Supplemental Table S14). This included the *Th*/*Slc6a3/Prlr* neurons (*q* = 1.15×10^−8^) and the *Th/Ghrh/Gal* cluster (*q* = 4.79×10^−5^) the latter of which were referred to as ‘neuroendocrine’ cells by Hook, McClymont (39), and the former a mixture of arcuate nucleus populations with *Prlr* was one of the marker genes, suggesting this includes the TIDA neurons. Additionally, P7 midbrain neurons were the other group with significant over-representation (specifically from the PAG and VTA) as well as the neuroblasts at this time point.

Although no specific adult mouse midbrain datasets exist, ventral midbrain sequencing at E11.5 - E18.5 by La Manno, Gyllborg (40) allowed us to identify imprinted enrichment within the midbrain at a timepoint when the major neuronal populations are differentiating but still identifiable (Supplemental Table S15). As anticipated, we found significant over-representation in both mature (DA1; high *Th* and *Slc6a3, q* = 0.0103), and developing (DA0, *q* = 0.0129) dopaminergic neurons, as well as the serotonergic neurons (*q* = 3.09×10^−7^), likely from the midbrain raphe nuclei.

Raphe nuclei from the midbrain/hindbrain are key serotonergic regions of the brain. Analysis of all cell types in the Dorsal Raphe Nucleus (DRN) sequenced by Huang, Ochandarena (41) revealed a clear enrichment of imprinted genes in the neuronal populations of the DRN as compared to the non-neuronal cell populations of the DRN (Supplemental Table S16A). When compared to all other cell populations, significant ORA was seen for Dopaminergic (*q* = 0.009), Serotonergic (*q* = 0.012) and Peptidergic neurons (*q* = 0.0008), however, a significant GSEA was found for all five neuronal populations (Supplemental Fig. S6). When compared against each other (i.e., serotonergic upregulation vs. the other neurons), only the serotonergic neurons of the DRN (*q* = 0.0019) were found to have a significant over-representation of imprinted genes (Supplemental Table S16B). GSEA’s were non-significant but the mean fold change for imprinted genes was markedly higher in both serotoninergic (52% higher) and dopaminergic neurons (68% higher). When contrasting neuronal subpopulations of the DRN, two of the five serotonin subpopulations had significant over-representation of imprinted genes: *Hcrtr1/Asb4* (*q* = 0.0014) and *Prkcq/Trh* (*q* = 0.007) (Supplemental Table S16C). These clusters were identified by Huang, Ochandarena (41) as the only clusters localised in the dorsal/lateral DRN and the serotonin clusters enriched in *Trh*. Huang, Ochandarena (41) hypothesised that these were the serotonin neurons that project to hypothalamic nuclei, and motor nuclei in the brainstem (as opposed to cortical/striatal projection).

### Imprinted gene expression is over-represented in lactotrophs and somatotrophs of the pituitary gland (Level 3D Analysis)

Following on from the enrichment seen above for imprinted gene expression in the dopaminergic arcuate nucleus neurons coordinating pituitary gland output, we sought to identify whether any cells in the pituitary would display matching over-representation for imprinted gene expression (Level 3D). The pituitary was not sequenced as part of the multi-organ or whole brain datasets analysed above and so two independent datasets were analysed that specifically sequencing the mouse pituitary at single cell resolution. Ho, Hu (8) recently sequenced the anterior pituitary gland of male and female C57BL/6 mice using two sequencing technologies, both 10X genomic and Drop-Seq. This identified a variety of cell types from the endocrine and non-endocrine pituitary. We analysed data from both technologies and found that imprinted gene expression was convergently over-represented in the Lactotrophs (prolactin secreting) and Somatotroph (growth hormone secreting) cells (Supplemental Table S17A & 17B). In a second independent dataset sequencing cells from male mouse pituitary glands (7), we found significant over-representation in the Somatotropes and Thyrotrope (secreting thyroid stimulating hormone). Figure 9 demonstrates the overlap in imprinted genes significantly expressed in Somatotropes and Lactotropes across the datasets since these were the only cell-types to be over-represented in more than one dataset (Supplemental Table S18). It is notable that the two cell types represented here directly match the two regulatory neurons found over-represented in the arcuate nucleus of the hypothalamus.

**Figure 9.**
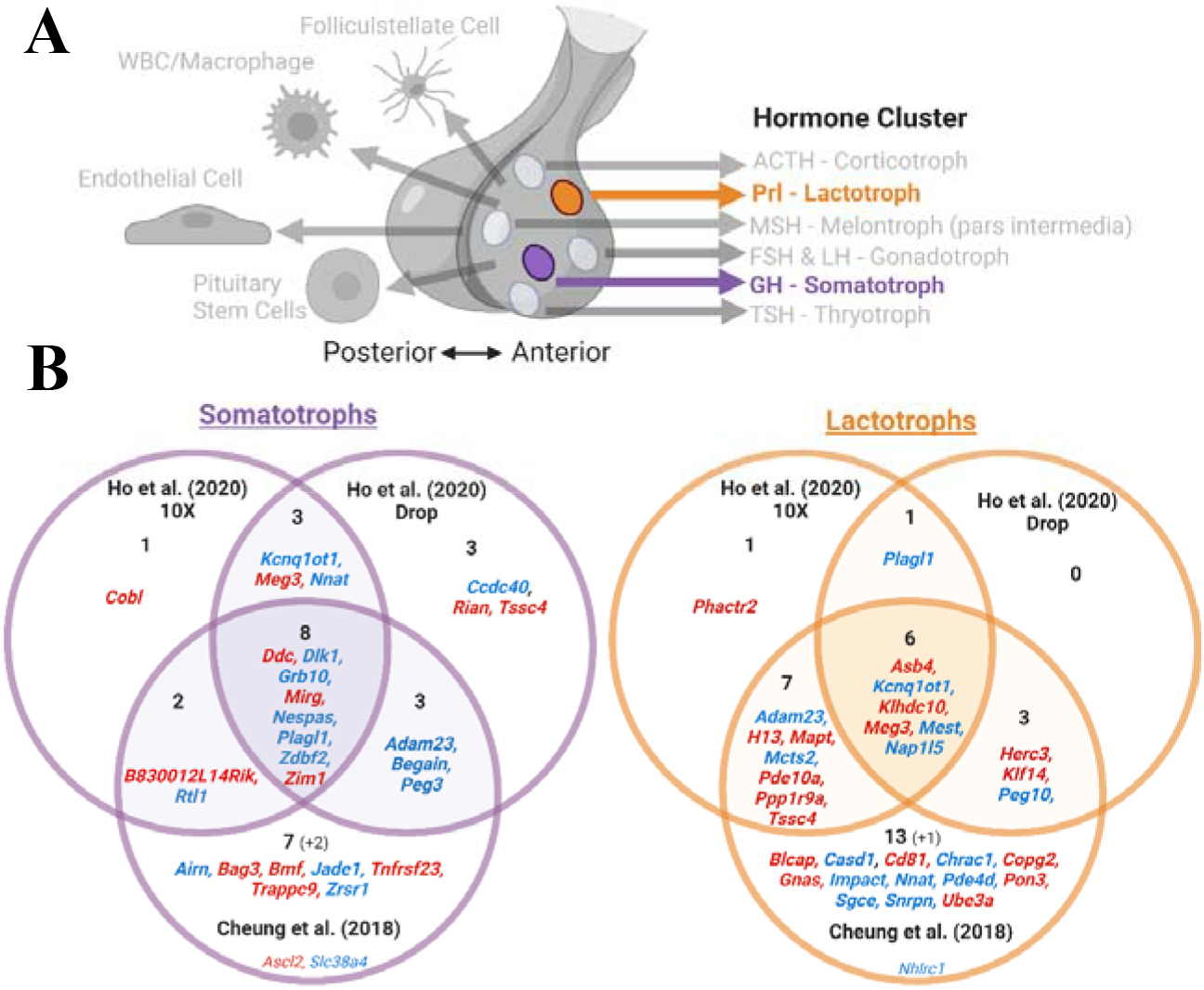
*(A)* Pituitary cell types showing over-representation for imprinted gene expression in multiple pituitary datasets. Over-represented cell types are bold and not in greyscale. The hormone/s released from the endocrine cell types are also indicated. *(B)* Venn diagram of upregulated imprinted genes in the Somatotrophs and Lactotrophs in Cheung, George (7) and Ho, Hu (8). Imprinted genes are listed which show significant upregulation (*q* ≤ 0.05 and Log2FC > 0) in the cell types. Parental bias is indicated by colour (MEG –red, PEG – blue). Genes in common from two analyses are presented in bold and totalled in each section of the Venn Diagram, while genes found upregulated in one analysis but not available in the others are included in small font and the number indicated in brackets.

## DISCUSSION

Using publicly available single cell transcriptomics data, we apply an unbiased systems biology approach to examine the enrichment of imprinted genes at the level of the brain in comparison to other adult tissues, refining this analysis to specific brain regions and then to specific neuronal populations. We confirm a significant over-representation in the brain, specifically in neurons at every level tested, with a marked enrichment in neuroendocrine cells lineages. Within-brain analyses revealed that the hypothalamus and the monoaminergic system of the mid- and hindbrain were foci for imprinted gene enrichment. While not all imprinted genes follow these patterns of expression, these findings highlight collective gene expression which is non-random in nature. As such, these analyses identify ‘expression hotspots’, which in turn suggest ‘functional hotspots’. Specifically, our results at the systems and cellular level highlight a major role for imprinted genes in the neuronal regulation of pituitary function, feeding and sleep.

Some of the earliest studies of genomic imprinting identified the brain as a key area for imprinted gene expression (20, 32). However, it is estimated that ~80% of the genome is expressed in the brain and consequently, imprinted gene expression here may not be a purposeful phenomenon. Our current analysis definitively show that imprinted genes were significantly over-represented in the brain as a whole. This over-representation was found again with PEGs alone, but not MEGs. Critically, over-representation was also found when limiting our analysis to genes that have been found to be imprinted in non-brain tissues (Supplemental Table S19). Excluding those genes imprinted in the brain alone avoids unintentionally biasing the analysis and confirms the robustness of the finding that imprinted gene expression is enriched in the brain.

Within specific brain regions, imprinted genes were over-represented in the hypothalamus, ventral midbrain, pons and medulla. This confirms some findings from studies of Pg/Gg and Ag chimera studies (20, 32) and summaries of imprinted gene expression (16). However, unlike these earlier studies, our analyses do not simply ask if imprinted genes are expressed (at any level) or not, but robustly test whether this expression is meaningful, and the expression of these genes are especially enriched in any given brain region. Additionally, in the chimera studies, Pg/Gg cells (two maternal genomes) preferentially allocated to the developing adult cortex and hippocampus, and Ag cells (two paternal genomes) preferentially allocated to the developing hypothalamus and midbrain. Our analysis does not reproduce this distinct pattern of MEG and PEG expression in the brain, and indeed we find no specific enrichment of imprinted genes in cortex or hippocampus. Although the pattern of regional enrichment seen with all imprinted genes is replicated when analysing PEGs alone, separate analysis of MEGs only shows over-representation in the pons and medulla. This difference between our findings and the Pg/Gg and Ag chimeras studies could indicate that the distribution of Pg/Gg and Ag cells in the brain is not driven by, or indeed reflective of adult PEG and MEG expression, but instead is determined by expression of specific imprinted genes during brain development (42).

At the whole brain level, mature neurons and, in particular, neural-lineage neuroendocrine cells, had disproportionately higher numbers of imprinted genes expressed, and high levels of imprinted gene expression. It is likely that this neural-lineage neuroendocrine population comprises members of the key hypothalamic populations in which the expression of imprinted genes are enriched and, when treated as their own cluster, demonstrate strong imprinted gene enrichment compared to other cell lineages of the brain, even other mature neurons.

Within the hypothalamus, a selection of informative neuronal subpopulations were over-represented. Strikingly, and suggestive of meaningful enrichment, we saw convergence across our different levels of analysis with several key neuronal types identified in the whole hypothalamus and/or hypothalamic-region-level, already having been identified against the background of general imprinted gene expression in the whole-brain-level analysis. These subpopulations are collectively associated with a few fundamental motivated behaviours. We consistently saw enriched imprinted gene expression in *Agrp* expressing neurons when contrasting neurons across the whole brain, whole hypothalamus and within the arcuate nucleus. *Agrp* neurons from the arcuate nucleus are well known feeding promotors and a few imprinted genes have previously been associated with their function (*Asb4, Magel2, Snord116*) (43, 44) but never as an enriched population. Feeding was further linked with imprinting through enrichment seen in *Pomc*+ neurons (45) as well as *Hcrt*+ and *Gal*+ neurons. Circadian processes are controlled principally by the Suprachiasmatic Nucleus and here we find strong imprinted gene enrichment in *Avp/Nms* expressing neurons (an active circadian population). Again, these were found to be enriched when contrasting neurons across the whole brain, whole hypothalamus and within the SCN. This population is of interest given the growing appreciation of the role imprinted genes play in circadian processes and the SCN suggested by studies of individual imprinted genes (46). Pituitary endocrine regulation also emerged as a key function, considering the over-representation in the dopaminergic: *Th/Slc6a3/Prlr* neuron type (top hit in the arcuate nucleus and across dopaminergic neurons of the brain) and the *Th/Ghrh* subpopulation. These neuron populations can regulate prolactin (regulating lactation, stress, weight gain, parenting and more (47, 48)) and growth hormone (promoting growth and lipid/carbohydrate metabolism) release, respectively. Remarkably, we also found a matching enrichment in the lactotroph and somatotroph cells in the pituitary. A role for imprinted genes in pituitary function is well known (49, 50), with pituitary abnormalities associated with imprinted disorders such as PWS (51) and recent sequencing work showing imprinted genes are amongst the highest expressed transcripts in the mature and developing pituitary (52). Specific genes highly expressed here, such as Dlk1 and Nnat, have been shown to alter somatotroph phenotypes (53, 54). Finally, we saw enrichment in galanin expressing neuronal populations (found enriched when contrasting neurons across the whole brain, whole hypothalamus). Galanin neurons in the hypothalamus have a diverse set of functions including subpopulations for thermoregulation, feeding, reproduction, sleep and parenting behaviour (55, 56), contributing to this consistent picture of IGs associating with neurons key for motivated behaviour.

In this analysis the hypothalamus was a clear hot spot for imprinted gene expression, in line with the prevailing view of imprinted gene and hypothalamic function (50, 57). However, outside of the hypothalamus other distinct hotspot emerged from our whole brain analysis including the monoaminergic system of the midbrain/hindbrain. Analysing data from the dorsal raphe nucleus and ventral midbrain revealed the dopaminergic and serotonergic neurons to be a foci of imprinted gene expression within this region. These midbrain dopamine neurons were enriched when contrasted to other dopamine neurons from the brain and the enriched serotonergic neurons were those that project to the subcortical regions of the brain known to be associated with feeding and other motivated behaviours (58), providing convergence with the functional hotspots seen in the hypothalamus.

Analyses of these kind are always bound by the available data and therefore there are notable limitations and caveats to this study. The aim of this study was to generate information about ‘hotspots’ of imprinted gene expression. This approach, and the use of over-representation analysis and GSEA, therefore do not provide an exhaustive list of sites of expression, and non-differentially expressed genes could still be highly expressed genes despite not contributing to this analysis. An example of a known site of expression for imprinted genes not found to be enriched in our analysis was the oxytocin neurons of the hypothalamus, since a clear oxytocin neuron phenotype has been reported in a handful of imprinted gene models (24, 59). This may be an example of a functional effect occurring below the level of over-representation, or that imprinted genes act during development and are not functionally enriched in adult oxytocin neurons, or simply that compared to other hypothalamic neuronal populations, oxytocin neurons are not a ‘hotspot’ of imprinted expression. Specific sequencing of oxytocinergic brain regions will be required to distinguish between these possibilities. A second caveat is that, due to the nature of the datasets used, not all imprinted genes were included, and our analysis was missing a significant subset of imprinted genes encoding small RNAs or isoforms from the same transcription unit. A third caveat is that we did not assess parent-of-origin expression for the 119 imprinted genes we included in the analysis. Previous expression profiling of imprinted genes have also not measured the POEs (16, 60) but have restricted their gene selection to genes with reliable imprinting status. Consequently, we only included the canonical imprinted genes and genes with more than one demonstration of a POE when looking for enrichment. Furthermore, for the vast majority of these genes, a brain-based POE effect has also already been reported (Supplemental Table S1). Although this does not replace validating the imprinting status of all 119 in the tissues and subregions examined, it does provide justification for looking at imprinted gene over-representation. To resolve this issue, scRNA-seq using tissues derived from reciprocal F1 crosses between distinct mouse lines will be key; for example, the recent work of (61) with cortical cell types provides an example of the allelic specific single-cell expression measurements necessary to confirm the enrichments found in this study.

By exploiting scRNA-seq data we have asked whether imprinted genes as a group are disproportionately represented in the brain, in specific brain regions, and in certain neuronal cell-types. In the adult brain imprinted genes were over-represented in neurons, and particularly the hypothalamic neuroendocrine populations and the monoaminergic hindbrain neurons, with the serotonergic neurons demonstrating the clearest signal. Interestingly, PEGs, but not MEGs, recreate this signal at Levels 1 and 2 - most notably only PEGs display the hypothalamic neuronal enrichment. By extension, these data also identify behaviours that are foci for the action of imprinted genes. Although there are high profile examples of individual imprinted genes expressed in the key brain regions we highlight and that have roles in feeding (*Magel2*) (62) and sleep (*Snord116*) (63), our analyses indicate that imprinted genes *as a group* are strongly linked to these behaviours and also identify other individual genes that should be explored in these domains. Conversely, there are high-profile examples of imprinted genes involved in hippocampus related learning and memory (*Ube3a*) (28), but we did not find enrichment for cell types related to this brain function. The idea that imprinted genes converge on specific physiological or behavioural processes is not unprecedented. Specialisation of function is predicted when considering why genomic imprinting evolved at all (13, 64–66). Moreover, there is increasing evidence that the imprinted genes themselves appear to be co-expressed in an imprinted gene network (IGN) and have confirmed regulatory links between each other (67–69). The idea of an IGN or, at the very least, heavily correlated and coordinated expression between imprinted genes adds further support to the idea that imprinted genes work in concert, rather than in isolation, to influence processes and that perturbating one may influence many others (70). Our findings add substance to these general ideas and highlight the neuronal regulation of pituitary function, feeding and sleep as being key functional hotspots on which imprinted genes converge which probably provides the best current basis for discerning evolutionary drivers of genomic imprinting in the brain.

## METHODS

### Data Processing

13 unique datasets were analysed across the three levels of analysis (see Fig.1) and analyses were conducted on each dataset independently. At each level of analysis, we aimed to be unbiased by using all the datasets that fitted the scope of that level, but the availability of public scRNA-seq datasets was limited, which prevented us from exploring all avenues (for example, a direct comparison of enrichment between hypothalamic nuclei). All sequencing data were acquired through publicly available resources and each dataset was filtered and normalised according to the original published procedure. Supplemental Table S20 details the basic parameters of each dataset. Once processed, each dataset was run through the same basic workflow (see below and Fig. 10), with minor adjustments laid out for each dataset detailed in the Supplemental Methods.

**Figure 10.**
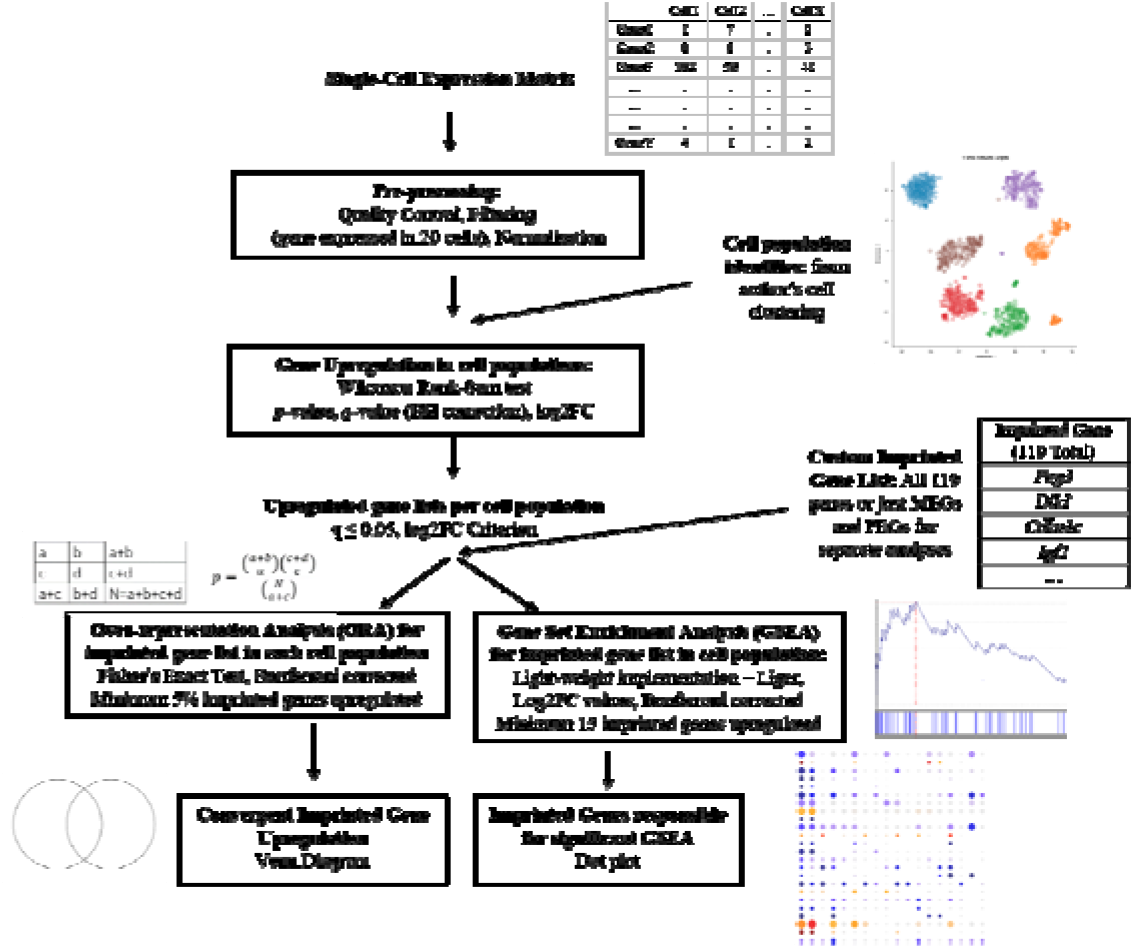
Basic workflow schematic. Single Cell Expression Matrices were acquired through publicly available depositories. Data were processed according to the author’s original specifications and all genes were required to be expressed in 20 or more cells. Cell population identities were acquired from the author’s original clustering. Positive differential gene expression was calculated via Wilcoxon Rank-Sum Test. Upregulated genes were considered as those with *q* ≤ 0.05 and a Log2FC ≥ 1 for analysis levels 1 and 2, while this criterion was relaxed to Log2FC > 0 for level 3. Our imprinted gene list was used to filter upregulated genes and two different enrichment analyses were carried out, over-

Due to the high variability in sequencing technology, mouse strain, sex and age, and processing pipeline, we have avoided doing analysis on combined datasets. Rather we chose to perform our analyses independently for each dataset and look for convergent patterns of imprinted gene enrichment between datasets on similar tissues/brain regions. As with any single-cell experiment, the identification of upregulation or over-representation of genes in a cell-type depends heavily on which other cells are included in the analysis to make up the ‘background’. Analysing separate datasets (with overlapping cell-types alongside distinct ones) and looking for convergent patterns of enrichment is one way of counteracting this limitation.

### Basic Workflow

Data were downloaded in the available form provided by the original authors (either raw or processed) and, where necessary, were processed (filtered, batch-corrected and normalized) to match the author’s original procedure. Cell quality filters were specific to each dataset and summarised in Supplemental Table S20. A consistent filter, to remove all genes expressed in fewer than 20 cells, was applied to remove genes unlikely to play a functional role due to being sparsely expressed. Datasets of the whole brain/hypothalamus were analysed both at the global cell level (neuronal and non-neuronal cells) and neuron specific level (only neurons) with genes filtered for the ≥ 20 cell expression at each level before subsequent analysis. Cell identities were supplied using the outcome of cell clustering carried out by the original authors, so that each cell included in the analysis had a cell-type or tissue-type identity. This was acquired as metadata supplied with the dataset or as a separate file primarily from the same depository as the data but occasionally acquired from personal correspondence with the authors. Cells were used from mice of both sexes when provided and all mice were aged 15 weeks or younger across all datasets. Although our focus was the adult mouse brain, embryonic data were included in some comparisons or when no alternatives were present. However, embryonic and post-natal cells were never pooled to contribute to the same cell populations.

Positive differential expression between identity groups were carried out using one-sided Wilcoxon rank-sum tests (assuming the average expression of cells within the current identity group is ‘greater’ than the average of cells from all other groups). The test was performed independently for each gene and for each identity group vs. all other groups. The large number of *p* values were corrected for multiple comparisons using a horizontal Benjamini-Hochberg correction, creating *q* values. Fold-change (FC) values, percentage expression within the identity group and percentage expressed within the rest were also calculated. We considered genes to be significantly positively differentially expressed (significantly upregulated) in a group compared to background expression if it had a *q* ≤ 0.05. In addition, for Level 1 and Level 2 analyses, the criteria for upregulated genes included demonstrating a Log2FC value of 1 or larger (i.e., 2-fold-change or larger). The datasets at these levels represented cells from a variety of organs, regions and cell-types, and in line with this cellular diversity, the aim of these analyses was to look for distinctive upregulation, akin to a marker gene. Once the analysis was restricted to cell subpopulations within a specific region of the brain (i.e., Level 3), the additional criteria for upregulation was relaxed to demonstrating just a positive Log2FC (i.e., the gene has a higher expression in this cell type than background). This was mainly because we were not expecting imprinted genes to be ‘markers’ of individual subpopulations at this level, but our aim was to identify enriched expression profiles for them. This additionally ensures consistent criteria for enrichment within levels, allowing meaningful comparison.

The same custom list of imprinted genes with reliable parent-of-origin effects (see below) was used for all analyses, and all genes were included as long as the gene passed the 20-cell filter. The first statistical analysis for enrichment was an Over-Representation Analysis (ORA) using a one-sided Fisher’s Exact Test (‘fisher.test’ function in R core package ‘stats v3.6.2’). The aim was to assess whether the number of imprinted genes considered to be upregulated as a proportion of the total number of imprinted genes in the dataset (passing the 20-cell filter) was statistically higher than would be expected by chance when compared to the total number of upregulated genes as a proportion of the overall number of genes in the dataset (passing the 20-cell filter). To limit finding over-represented identity groups with only a few upregulated imprinted genes, an identity group was required to have ≥ 5 % of the total number of imprinted genes upregulated for ORA to be conducted. Subsequent p-values for all eligible identity groups were corrected using a Bonferroni correction. This provided a measure of whether imprinted genes are expressed above expectation (as opposed to the expression pattern of any random gene selection) in particular identity groups.

Venn diagrams of the upregulated imprinted genes making up over-represented identity groups across datasets (within a level) were also reported. Full lists of upregulated imprinted genes can be found in the ‘Upregulated_IGs.csv’ file for each analysis in the Supplemental Data.

To further examine the presence of imprinted genes within tissues/cell types, and to provide a different perspective to over-representation, we conducted a Gene-Set Enrichment Analysis (GSEA) for imprinted genes amongst the upregulated genes of an identity group using a publicly available, light-weight implementation of the GSEA algorithm (71) in R (https://github.com/JEFworks/liger). This was done in a manner similar to Moffitt, Bambah-Mukku (72) since we were similarly using this computational method to identify enrichment of our gene sets *within the upregulated genes* of the different identity groups. Here, the GSEA was conducted for each individual identity group using Log2FC values to rank the upregulated genes. The GSEA acts as a more conservative measure than the ORA since it tests whether imprinted genes are enriched in the stronger markers of a group (the genes with the highest fold change for a group vs. the rest) and hence whether the imprinted genes are enriched in those genes with a high specificity to that tissue/cell type. To prevent significant results being generated from just 2 or 3 genes, identity group to be analysed were selected as having a minimum of 15 upregulated imprinted genes (i.e. the custom gene set) to measure enrichment for (a value suggested by the GSEA user guide (https://www.gsea-msigdb.org/gsea/doc/GSEAUserGuideFrame.html)) and to prevent significant results in which imprinted genes cluster at the tail, identity groups were selected as having an average fold change of the upregulated imprinted genes greater than the average fold change of the rest of the upregulated genes for that group. Again, multiple p values generated from GSEA were corrected using a Bonferroni correction. To further elucidate the genes responsible for significant GSEA’s, dot plots of the imprinted genes upregulated in that identity group were plotted across all identity groups with absolute expression and Log2FC mapped to size and colour of the dots, respectively. Graphical representations of significant GSEA’s (post-correction) are included in the main text or as supplemental figures, all other graphs, including additional dot plots not discussed in this study, can be found in the repository (https://osf.io/jx7kr/) and Supplemental Data. If no cell populations met these criteria, GSEA was not run and not included for that analysis.

For Level 1 and Level 2 analyses, we also carried out parent-of-origin specific analyses. The imprinted gene list was divided into MEGs and PEGs and the analyses detailed above were run separately for these two gene groups. For imprinted genes with known parent-of-origin variability based on tissue type (*Igf2* and *Grb10*), the parent-of-origin characterisation of these genes was changed accordingly. The absolute number of imprinted genes top-expressed in a tissue/cell-type were also reported for analyses in Level 1 and Level 2 in the tables, since these analyses included a variety of cell-types and tissues which may demonstrate meaningful clustering of the highest normalised expression values. The mean normalised expression for all imprinted genes across the series of identity groups in the datasets in Level 1 and Level 2 was also calculated alongside the mean normalised expression for the rest of the genes (Supplemental Table S2).

All graphical representations and statistical analyses were conducted using R 3.6.2 (73) in RStudio (74). Diagrams in Figures 1:2, 4:6 and 8:10 were created with BioRender.com.

### Custom Imprinted Gene List

The gene list for the analysis was based on the list of murine imprinted genes recently published in Tucci, Isles (10). Although the original list of imprinted genes was 260 genes long, only 163 genes were identified in the most comprehensive of the datasets. We further refined this list to 119 imprinted genes (Supplemental Table S1a) which excluded the X-linked genes, consisting of mostly the canonical protein-coding and long noncoding RNA imprinted genes, but the criteria for inclusion was those genes with at least two independent demonstrations of their POE status (See Supplemental Table S1b for full list of 260 imprinted genes and reasons for gene exclusion). The only exceptions to multiple independent demonstrations of a POE were four genes (*Bmf, B3gnt2, Ptk2, Gm16299*) identified by (34) where a POE was assessed across 16 brain regions and 7 adult tissues within one study. For Level 2, the MEG/PEG status of a gene was primarily based on reported allelic expression within the brain. Small non-coding RNAs such as micro-RNAs (miRs) and small nucleolar RNAs (snoRNAs), which represent ~10% of identified imprinted genes, were excluded from the analysis as their sequences were not detected/subsumed by larger transcripts in the majority of the datasets. Another caveat with short-read RNA-seq libraries is that much of the expression data for a given transcription unit cannot discriminate differentially imprinted isoforms nor do some of the technologies (e.g., Smart-Seq2) possess stranded libraries to distinguish antisense transcripts. For complex imprinting loci such as the *Gnas* locus, most reads as result map to only *Gnas* and *Nespas* ignoring several overlapping and antisense genes.

## DECLARATIONS

### Ethics approval and consent to participate

Not applicable. All samples had been collected in the context of previous studies.

### Consent for publication

Not applicable.

### Availability of data and materials

The datasets analysed during the current study were acquired from publicly available resources and are available in the following GEO repositories, Mouse Cell Atlas – <u>GSE108097</u>, *Tabula Muris* – <u>GSE109774</u>, Aging Mouse Brain – <u>GSE129788</u>, Hypothalamus (Chen) – <u>GSE87544</u>, Hypothalamus (Romanov) – <u>GSE74672</u>, Arcuate Nucleus – <u>GSE93374</u>, Suprachiasmatic Nucleus – <u>GSE132608</u>, Dopamine Neurons – <u>GSE108020</u>, Ventral Mid Brain – <u>GSE76381</u>, Dorsal Raphe Nucleus – <u>GSE134163</u>, Pituitary Gland (Ho) - <u>GSE146619</u>, Pituitary Gland (Cheung) - <u>GSE120410</u> and the following SRA repository, Mouse Brain Atlas – <u>SRP135960</u>. The data generated in this experiment is provided as Supplemental Data and in an Open Science Framework repository entitled – “Imprinted Gene Enrichment at Single-Cell Resolution” (https://osf.io/jx7kr/). Custom R scripts to analyse each dataset are provided as Supplemental Code and are available at https://github.com/MJHiggs/IG-Single-Cell-Enrichment.

## Authors’ Contributions

MJHiggs performed bioinformatic analysis, with input from MJHill; MJHiggs., and ARI contributed to project design, data interpretation, and wrote the manuscript, MJHiggs produced all Figures; all co-authors reviewed and edited the manuscript.

## Acknowledgements

We would like to thank all the research groups that carried out the single-cell RNA sequencing that made this study possible and to particularly acknowledge Dr. L. Cheung, Dr. P. Hook, Dr. A. Jackson, and Dr. S. Wen for help accessing the cell metadata for their associated studies.

