## Supplementa Materials for "Systematic investigation of imprinted gene expression and enrichment in the mouse brain explored at single-cell resolution"

**Supplemental Material**

M. J. Higgs^1^, M. J. Hill^2,3^, R. M. John^4^, and A. R. Isles^1,*^

1 Behavioural Genetics Group, MRC Centre for Neuropsychiatric, Genetics and Genomics, Neuroscience and Mental Health Research Institute, Cardiff University, Cardiff, UK

2 Division of Psychological Medicine and Clinical Neurosciences, MRC Centre for Neuropsychiatric Genetics and Genomics, School of Medicine, Cardiff University, Cardiff, UK

3 UK Dementia Research Institute, School of Medicine, Cardiff University, Cardiff, UK

4 School of Biosciences, Cardiff University, Cardiff, UK

* Corresponding author: A. R. Isles, Behavioural Genetics Group, MRC Centre for Neuropsychiatric Genetics and Genomics, Neuroscience and Mental Health Research Institute, Cardiff University, Cardiff CF24 4HQ, UK.

**Table of Contents**

Supplemental Methods	3

Specific processing steps for each unique dataset	3

1) *Level 1 - Multi-Organ Level Analyses	3*
2) *Level 2 - Whole Brain Level Analyses	4*
3) *Level 3 - Brain Region Level	5*

Supplemental Figures	10

Supplemental Figure S1.	10

Supplemental Figure S2.	11

Supplemental Figure S3.	12

Supplemental Figure S4.	13

Supplemental Figure S5.	15

Supplemental Figure S6.	16

Supplemental Tables	18

Supplemental Table S1	18

Supplemental Table S2	18

Supplemental Table S3 A & B	18

Supplemental Table S4 A & B	19

Supplemental Table S5 A & B	20

Supplemental Table S6 A & B	21

Supplemental Table S7 A & B	22

Supplemental Table S8	23

Supplemental Table S9 A & B	28

Supplemental Table S10 A & B	29

Supplemental Table S11	30

Supplemental Table S12	31

Supplemental Table S13	31

Supplemental Table S14	32

Supplemental Table S15	32

Supplemental Table S16 A, B & C	32

Supplemental Table S17 A & B	33

Supplemental Table S18	34

Supplemental Table S19	34

Supplemental Table S20	35

References for Supplemental Material	36

**Supplemental Methods**

**Specific processing steps for each unique dataset**

**1) Multi-Organ level**

Mouse cell atlas (adult and mouse – 43 tissues) (1)

The Mouse Cell Atlas (MCA) involved the collection and sequencing of 50+ tissues in an attempt to distinguish genetic signatures of tissues across a mouse’s life, of which 43 passed the author’s quality control measures. These tissues included: 21 adult tissues (from 6-10 weeks old C57BL/6J mice, including the mammary gland sequenced at four separate life stages), 11 fetal tissues (E14.5), six neonatal tissues and 4 stem cell derived populations. An Expression matrix (‘MCA_Figure2-batch-removed.txt.tar.gz’) of 60,000 cells of high quality (~1500 cells from the 43 tissues) was downloaded from Figshare (<https://figshare.com/articles/MCA_DGE_Data/5435866>) alongside the cell metadata (‘MCA_Figure2_Cell.info.xlsx’). Raw data available from GEO repository, [GSE108097](https://www.ncbi.nlm.nih.gov/geo/query/acc.cgi?acc=GSE108097). Data were filtered for the 18 unique adult tissues (including the mammary gland from virgin females only). Data were then scaled to 100,000 reads and log normalised. Tissue cell identities were acquired from the ‘Tissue’ annotations. Data were run through the core workflow once for all adult tissues.

Tabula Muris (adult mouse – 20 tissues) (2)

The Tabula Muris Consortium released the Tabula Muris soon after the MCA. The Tabula muris comprises 20 adult mouse organs with a partial overlap with the MCA but sequenced these tissues to a much greater depth and using two sequencing methods. Droplet sequencing allowed the surveying of a large number of cells at relatively low coverage and was completed for 13 organs. FACs sorting allowed characterisation of fewer cells from the organ but at much higher sensitivity and depth and was completed for all 20 organs. This was completed using 3 female mice and 4 male mice, all 10-15 weeks old and on a C57BL/6JN background. The FACs dataset was selected for analysis as it contained the largest number of tissues and the only one to include the brain (broken down as myeloid brain cells (neurons, glia, endothelial etc) and non-myeloid brain cells (microglia and macrophages)). Clustering for cells in this dataset allowed cell identification at the Tissue and cell identity level. All data were downloaded through Figshare as Robjects ([https://figshare.com/articles/Robject_files_ for_tissues _processed_by_Seurat/5821263/1](https://figshare.com/articles/Robject_files_%20for_tissues%20_processed_by_Seurat/5821263/1)). Raw data and processed data were accessible from these files and cell metadata were acquired as annotation csv’s from the respective sequencing technology Figshare page (FACS - [https://figshare.com/articles/Single-cell_RNA- seq_data_ from_Smart-seq2_sequencing_of_FACS_sorted_cells_v2_/5829687](https://figshare.com/articles/Single-cell_RNA-%20seq_data_%20from_Smart-seq2_sequencing_of_FACS_sorted_cells_v2_/5829687)). Raw data also available from GEO repository, [GSE109774](https://www.ncbi.nlm.nih.gov/geo/query/acc.cgi?acc=GSE109774). Raw data were used to identify genes expressed in 20 or more cells and processed data were then used unaltered for the analysis. Tissue cell identities were acquired from the ‘tissue’ annotations and run through the core workflow once for all adult tissues.

**2) Whole Brain level**

Mouse Brain Atlas (3)

Zeisel, Hochgerner (3) created the Mouse Brain Atlas in 2018 – a comprehensive sequencing of the adolescent mouse nervous system. This involved sequencing cells from the entire Central and the Peripheral nervous system, including the enteric nervous system. Cells were clustered into class (e.g., neuron, oligodendrocyte), region of origin (e.g., hypothalamus, pons) and specific cell type by neurotransmitter (e.g., serotonergic, dopaminergic) and most likely nuclei of origin. Data were downloaded from mousebrain.org (<http://mousebrain.org/downloads.html>) and the Level 5 data were downloaded as a one cell per column loom file which also included the cell metadata as column attributes (“l5.all.loom”). Data were then log normalised and scaled to 5,000 reads. Cell lineage labels were acquired from ‘Class’ annotations, cell tissue labels were acquired from ‘tissue’ annotations and cell identity labels were acquired from ‘ClusterName’ annotations. The core workflow as run three times, once for cell lineage with all cells, before restricting the analysis to neurons only and running it once for nervous system regions and once for the specific neuron identities across the nervous system. Marker genes to identify neuron subpopulations were acquired from combined enrichment analysis with trinaization scores carried out by the original authors located from their online resource (<http://mousebrain.org/>). Two additional marker genes were included – *Esr1* and *Prlr* – for Deinh5 neurons based on our own analyses. Raw data also available from SRA repository, [SRP135960](https://www.ncbi.nlm.nih.gov/sra/SRP135960)

Single-Cell Transcriptome profiling of the aging mouse brain (4)

To complement the analysis of the mouse brain atlas, we also analysed Ximerakis, Lipnick (4)’s single cell profile of the young and old mouse brains. Although the original aim of the study was to identify aging-related genes/pathways, the data could also be used to investigate differential expression between the different cell types within the brain, grouped by cell lineage. 16 C57BL/6 mice (8 at 2/3 months and 8 at 21/22 months) had whole brains dissected and hindbrain removed before tissue was dissociated into single cells. For this analysis we chose to only include the 2–3-month-old mouse data which was the most comparable to the age of mice from other studies used by us. Expression data and meta data were acquired from Gene Expression Omnibus through accession no. [GSE129788](https://www.ncbi.nlm.nih.gov/geo/query/acc.cgi?acc=GSE129788), scaled to 10,000 reads and log normalised. Data were run through the workflow once comparing all major CNS (minus hindbrain) cell types.

**3) Brain Region level**

***3a) Whole Hypothalamus***

**Chen, Wu (5)**

Chen, Wu (5) sequenced adult (8-10 weeks) whole hypothalamus of female B6D2F1 mice (C57B6 female × DBA2 male). Processed/normalised data were downloaded as an R object from Gene Expression Omnibus through accession no. [GSE87544](https://www.ncbi.nlm.nih.gov/geo/query/acc.cgi?acc=GSE87544) alongside count data and a metadata file with the cell identities. The original 14,437 cells were filtered to 3,319 high quality cells by the authors (containing > 2000 unique genes expressed) and this filtered number was maintained for this analysis. Cell identities included 11 non-neuronal and 34 neuronal identities identified as Snap25/Syt1-high (and split into GABA and Glut populations alongside a Hista cluster). Cells without a cluster (labelled as ‘zothers’) were discarded from the analysis leaving 2275 neurons. Marker genes supplied for these neuronal identities were identified by the original authors. Cell identities for all cells were taken from ‘SVM_clusterID’ annotations from the metadata file. The core workflow as then run twice, once for all cells and once for just the neurons.

**Romanov, Zeisel (6)**

To complement the previous study, the data from Romanov, Zeisel (6) were also analysed. The authors sequenced cells from “a central column” of the mouse hypothalamus which spanned from the posterior POA to the ARC (rostro-caudal), PVN to the VLH (dorsal-lateral). Mice were on C57BL6/N background, of both sexes and aged 14-28 days. The expression matrix and meta data was acquired from a single file available from Gene Expression Omnibus through accession no. [GSE74672](https://www.ncbi.nlm.nih.gov/geo/query/acc.cgi?acc=GSE74672). The expression matrix was prefiltered by the authors to 2,882 high quality cells (>1,500 molecules excluding rRNA) and was scaled to 10,000 reads and log2 normalised. Cell identities included 6 non-neuronal identities and 62 neuronal subtypes (clustered from 898 neurons) identified by expression of key neuropeptides (e.g., *Avp, Oxt*, DA). The ‘level1 class’ annotations were used for non-neuronal populations and the ‘level2 class (neurons only)” annotations were used for the 62 neuronal identities. The core workflow was then run twice, once for all cells and once for just the neurons.

***3b) Specific Hypothalamic Nuclei***

Several datasets focusing on individual regions of the hypothalamus were in existence prior to this investigation and each was investigated individually for imprinted enrichment.

Arcuate Nucleus (ARC) (7)

Campbell, Macosko (7) sequenced the arcuate-median eminence complex of adult C57BL/6J mice under different feeding conditions. They identified 65 distinct cellular identities of which 34 were unique neuronal identities defined by key neuropeptides and novel markers (and 24 originated from the arcuate nucleus). Raw and processed/filtered expression matrices alongside the cell metadata (including cluster annotations) were acquired from Gene Expression Omnibus through accession no. [GSE93374](https://www.ncbi.nlm.nih.gov/geo/query/acc.cgi?acc=GSE93374). Raw data were scaled to 10,000 transcripts per cell (following Timshel, Thompson (8) and log normalised. Cell labels were acquired from the “Subcluster” annotations from the metadata. Data were run through the workflow twice, once with all neurons and a second time with only ARC neurons (removing the 10 suspected non-ARC subpopulations identified by the original authors).

Suprachiasmatic Nucleus (SCN) (9)

In a spatio-temporal manner, Wen, Ma (9) used single-cell RNA sequencing to investigate the basic and circadian expression of various cell types in the SCN by micro dissecting the SCN and surrounding areas of 55 virgin 7-8 week old male C57BL/6J mice and sequencing them at 12 circadian time points (5 per timepoint). For the main experiment, conducted via Drop-seq, 8 major cell types (incl. neurons) were identified and within the neuronal cluster, 5 SCN neuronal subtypes were identified (ignoring neuronal subtypes likely from other neighbouring brain regions). Neuron identities were then reconfirmed with 10x Genomics, processing an additional 8,679 cells including 1,251 SCN neurons. These neurons were classified into the same 5 neuronal clusters as drop-seq and found a high convergence between the groups. Since the 10x data was more comprehensive for neurons, this data were chosen to look for neural specific enrichment only. The 10X Seurat files was acquired from Gene Expression Omnibus from accession no. [GSE132608](https://www.ncbi.nlm.nih.gov/geo/query/acc.cgi?acc=GSE132608) and a Seurat object was created. Cell metadata were acquired through personal communication with the authors. SCN neuron identities were acquired from the ‘cluster’ annotation. The Data were scaled by 5000 reads per cell, log-normalised and run through the workflow once.

***3c) Monoaminergic Nuclei***

In addition to the specific hypothalamic nuclei, we decided to investigate imprinted gene expression in other brain areas of interest, namely the dopamine neurons of the midbrain and the various neurons of the Dorsal Raphe Nucleus, both of which had datasets available and were regions of over-representation in the analysis of the Mouse Brain Atlas (6).

Dorsal Raphe Nucleus (10)

The Dorsal Raphe Nucleus is a critical nucleus in the CNS responsible for a significant proportion of neuromodulators, and most significantly, approximately a third of all serotonergic neurons. Huang, Ochandarena (10) sequenced the dorsal raphe nucleus and surrounding areas of 8-10-week-old C57BL/6J mice. They discovered 17 major clusters (of which 5 were major neuron classes e.g., serotonergic, dopaminergic) and the neuron classes were found to have an additional 17 subclusters. Raw data is available at GEO depository, [GSE134163](https://www.ncbi.nlm.nih.gov/geo/query/acc.cgi?acc=GSE134163). Data were downloaded as processed and annotated Seurat objects (already log normalised, scaled to 10,000 UMI) for each of the major cell types from Harvard Dataverse ([https://doi.org/10.7910/DVN/ QB5CC8](https://doi.org/10.7910/DVN/%20QB5CC8)). The workflow was run three times, once using major cell cluster identities acquired from the ‘Curated_cellTypeLabels’ annotations and once using just neurons with the major cell clusters and a final time using just the neurons with the sub cluster identities acquired from the ‘Curated_subtypeLabels’.

Brain-wide Dopamine Neurons (11)

The other key neuron type that suggested a presence of imprinted genes was the dopamine neuron population. Hook, McClymont (11) sequenced 473 dopamine neurons in the  C57BL/6J mouse brain at embryonic (E15.5) and early postnatal (P7). Midbrain and Forebrain dopamine neurons were microdissected from E15.5 mouse brains and olfactory bulb, midbrain and forebrain (arcuate nucleus) dopamine neurons from P7 mice, all identified using FACS. A total of 13 cell populations were found (four at E15.5 and nine at P7) which included clustering of dopamine neurons from the arcuate nucleus, VTA, PAG and SN individually. The raw and normalised expression matrix, pre-filtered for the 396 dopamine neurons meeting the author’s quality criteria was acquired from github ([https://github.com/ pwh124/sc-da-parkinsons](https://github.com/%20pwh124/sc-da-parkinsons)) along with accompanying cell metadata (cell cluster identities). Raw data is available at GEO repository, [GSE108020](https://www.ncbi.nlm.nih.gov/geo/query/acc.cgi?acc=GSE108020). Data were run through the workflow once.

Developing Midbrain (12)

As a final investigation into the presence of imprinted genes in the midbrain, the data from La Manno, Gyllborg (12) were analysed. The authors sequenced single cells from the CD-1 mouse ventral midbrain at time points between e11.5 – e18.5 covering dopaminergic progenitor specification and differentiation. Over this time period, 26 distinct cellular identities were identified including a variety of unique neuronal identities (e.g., serotonergic, dopaminergic, GABAergic). Data were downloaded from Gene Expression Omnibus using accession no. [GSE76381](https://www.ncbi.nlm.nih.gov/geo/query/acc.cgi?acc=GSE76381). The expression matrix was scaled to 10,000 reads and log normalised. Metadata was included as a row within the gene expression matrix and cell identities were prescribed using the ‘cell-type’ annotations. Data were run through the workflow once using all cell identities.

***3d) Pituitary Gland***

Anterior Pituitary Gland (13)

Ho, Hu (13) sequenced the anterior pituitary gland using both 10X genomics and Drop-seq platforms. The 10x genomics experiment involved 2,780 cells from 8-week-old CD1 males and females. The Drop-seq experiment sequenced cells from WT mice at 8 weeks but also 13week old and *mt*/*hGRF* transgenic mic, for consistency only the 8-week WT mice were analysed resulting in 4,663 cells. Both 10x and Drop-seq matrices were acquired from a public google drive as RDS datasets - <https://drive.google.com/drive/folders/1cbwkQl1Hh70xsToNhjOdom_ru7XGA3l->. Cell metadata for both experiments were acquired from personal correspondence with the authors and cell identities were acquired from the ‘cluster’ and ‘genotype’ annotation. Raw data are available at GEO depository, [GSE146619](https://www.ncbi.nlm.nih.gov/geo/query/acc.cgi?acc=GSE146619). Analysis pathways to filter, normalise and scale the data were acquired from <https://github.com/wulabupenn/mPit> for both datasets and data were processed until the point of pre-clustering following the author’s workflow. Additionally, genes were filtered to those expressed in 20 or more pituitary cells. Drop-seq data was additionally filtered to select only cells from 8-wk-old-WT. The workflow was run once for each dataset comparing all cell types across both sexes.

Pituitary Gland (14)

As an additional independent analysis of the pituitary gland was available through Cheung, George (14)’s sequencing of 6, 7-week-old male C57BL/6 pituitary glands. This involved 13,663 cells with many overlapping cell types with Ho et al., 2020. Files necessary to create a Seurat object were acquired from Gene Expression Omnibus through accession no. [GSE120410](https://www.ncbi.nlm.nih.gov/geo/query/acc.cgi?acc=GSE120410) and manual cell clusters were acquired from personal correspondence with the author’s. Data were scaled to 10,000 reads and log transformed. The workflow was run once, for all major cell types. Cells were run through the workflow once including all cell types.

**Supplemental Figures**

**Log2FC Values**

**Enrichment Score (cumulative weight)**

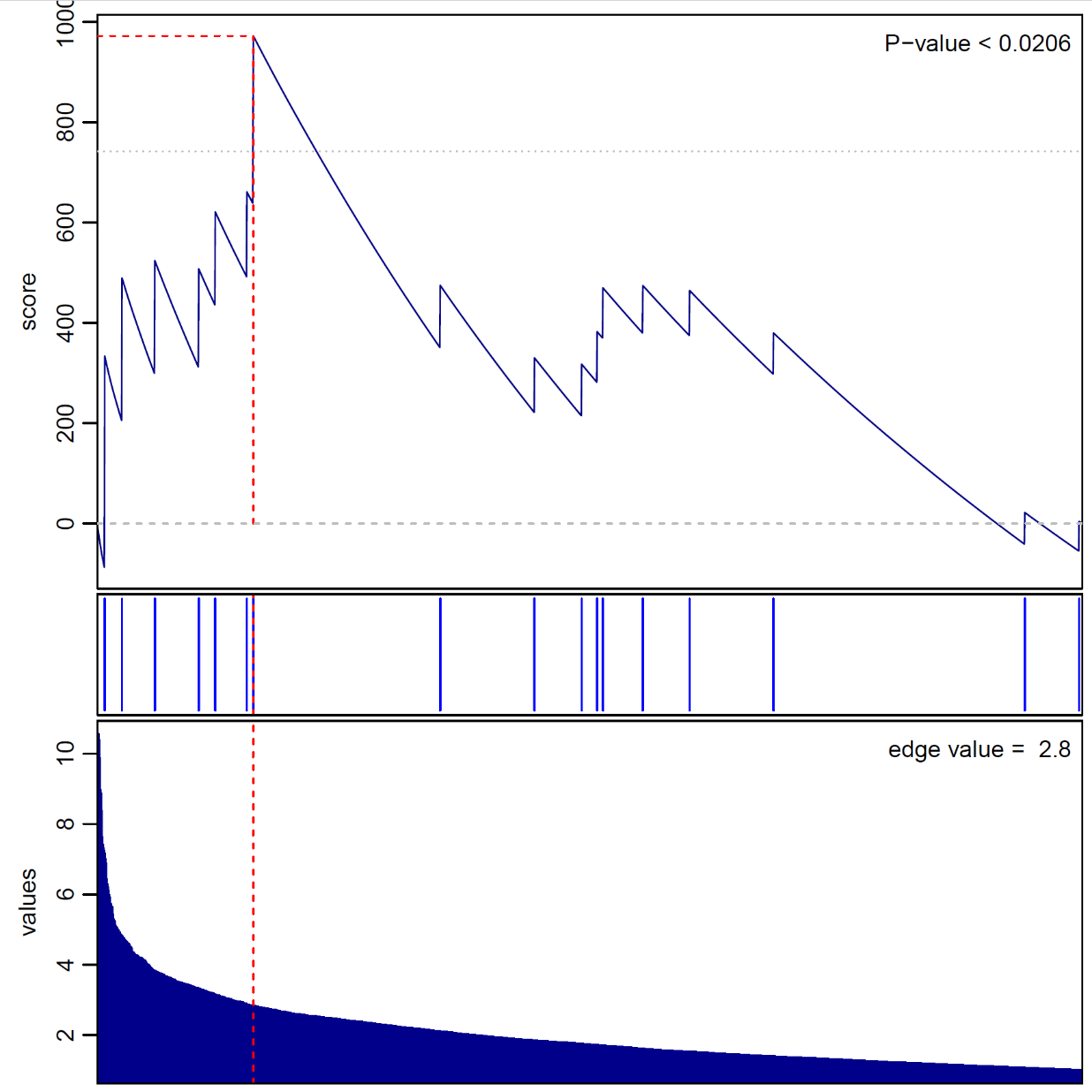

**Sorted Genes (by descending Log2FC)**

**Supplemental Figure S1.** Significant Gene Set Enrichment Analysis (GSEA) for Paternally Expressed Genes (PEGs) in the cell of the Pancreas in the Mouse Cell Atlas (1). In the GSEA analysis, genes are sorted by strength by which they mark this neuronal cluster (sorted by fold change values) indicated by the bar (middle). Fold change values are displayed along the bottom of the graph. The genes are arrayed left (strongest marker) to right and blue lines mark where imprinted genes fall on this array. The vertical axis indicates an accumulating weight, progressing from left to right and increasing or decreasing depending on whether the next gene is an imprinted gene or not. The *p*-value represents the probability of observing the maximum value of the score (red dashed line) if the imprinted genes are distributed randomly along the horizontal axis.

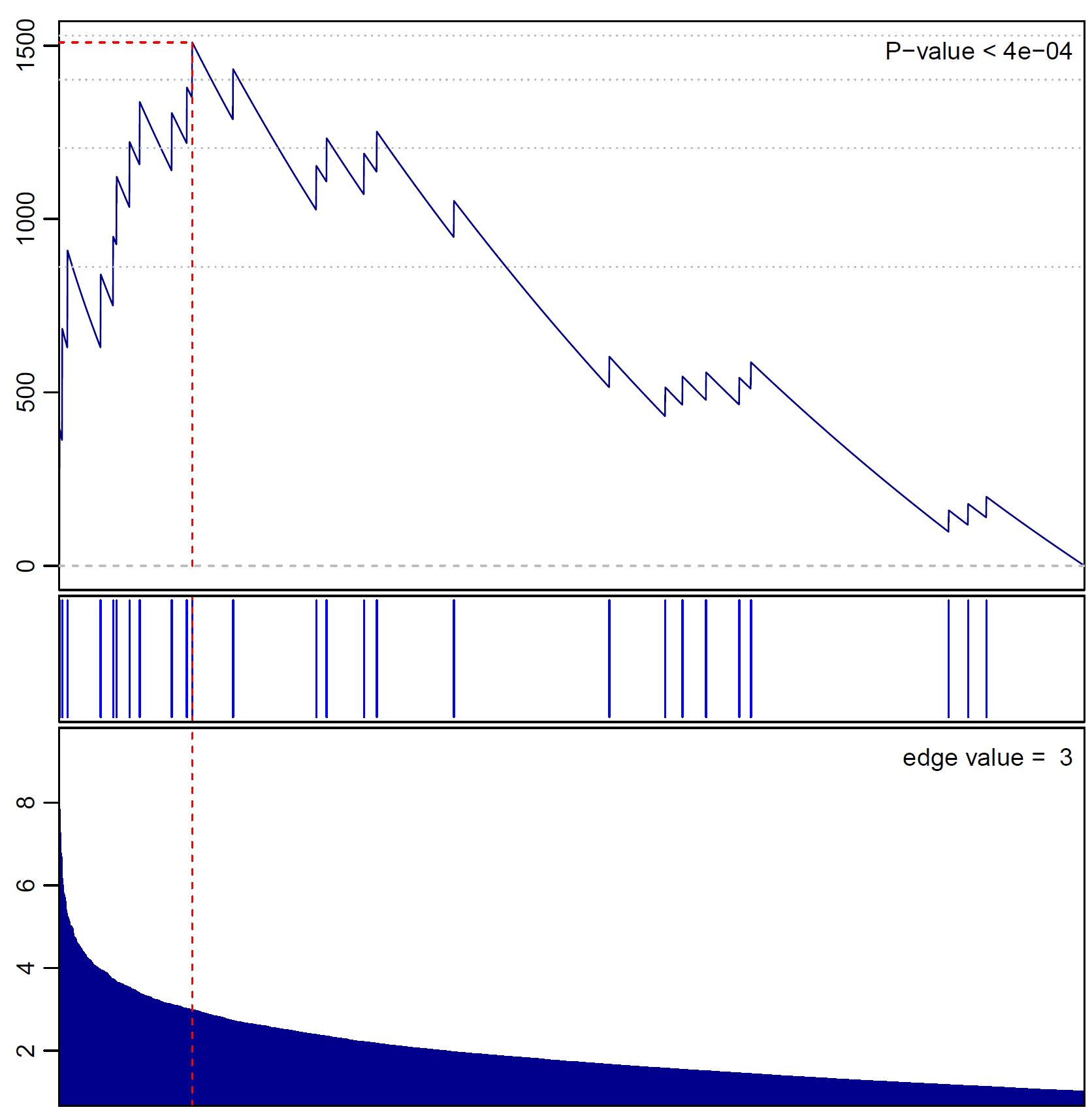

**Log2FC Values**

**Enrichment Score (cumulative weight)**

**Sorted Genes (by descending Log2FC)**

**Supplemental Figure S2.** Significant GSEA for PEGs in Neuroendocrine Cells in a Whole Mouse Brain Analysis (4). See Figure S1 legend for description on how to interpret GSEA analysis and plot.

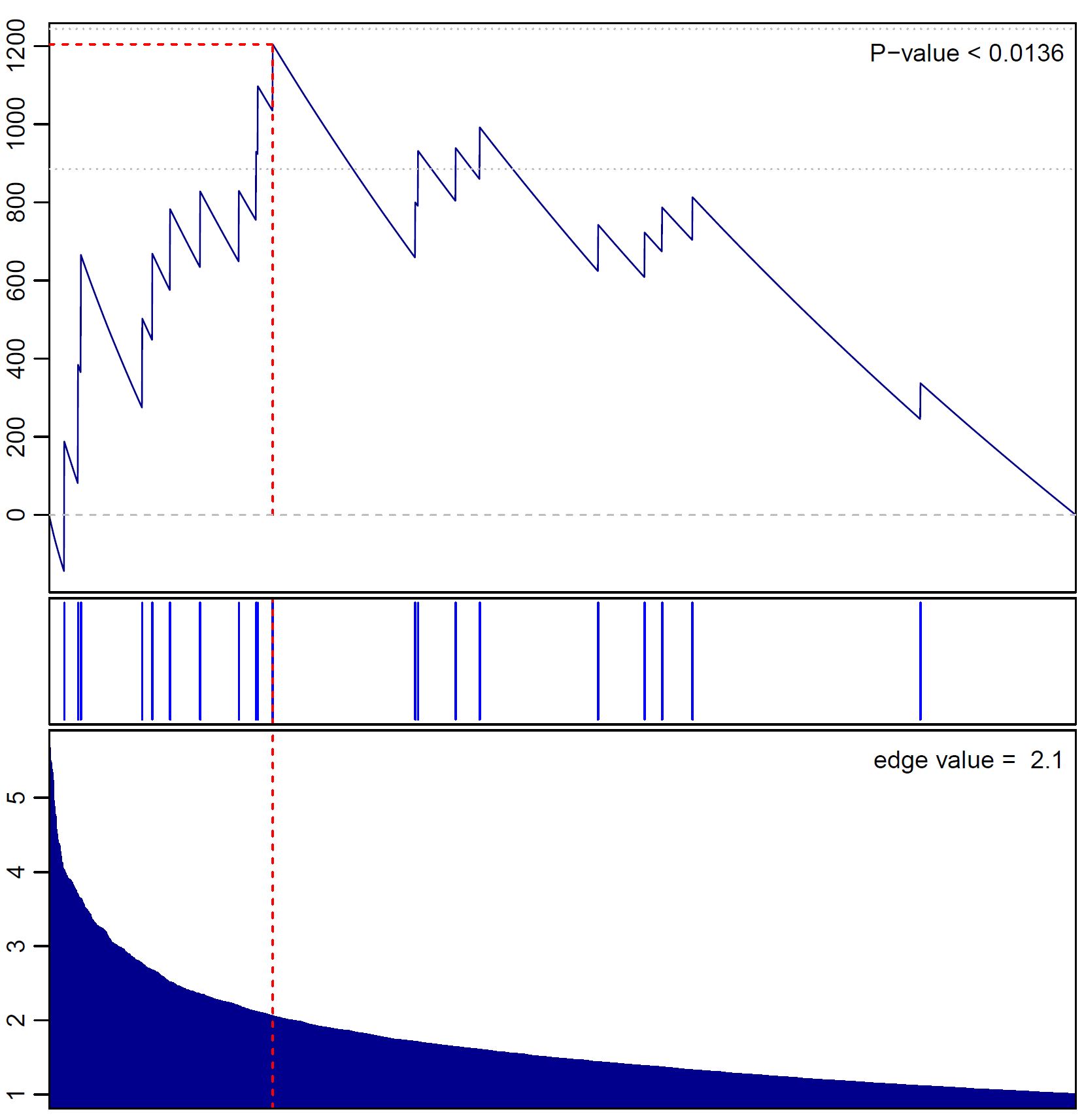

**Sorted Genes (by descending Log2FC)**

**Log2FC Values**

**Enrichment Score (cumulative weight)**

**Supplemental Figure S3.** Significant GSEA for Maternally Expressed Genes (MEGS) in the cells of the Pons in the Mouse Brain Atlas Analysis (3), *q*-value = 0.0272. See Figure S1 legend for description on how to interpret GSEA analysis and plot.

**A**

**B**

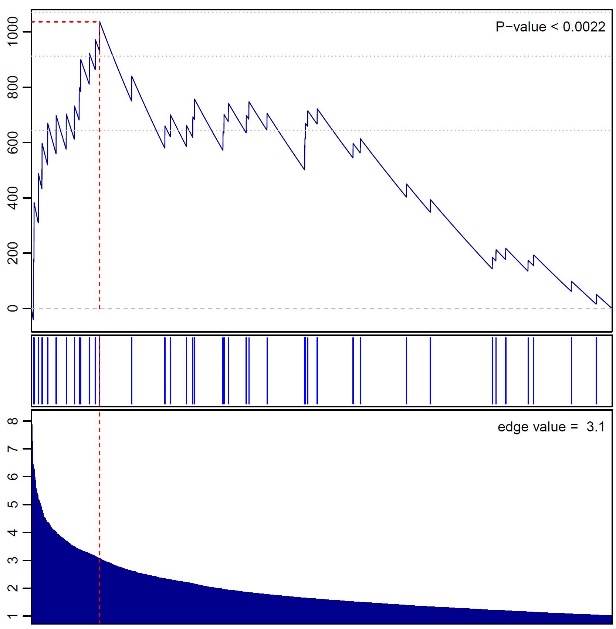
 **HBSER2 (*q* = 0.0462)**	**HBSER4 (*q* = 0.011)**

**Enrichment Score (cumulative weight)**

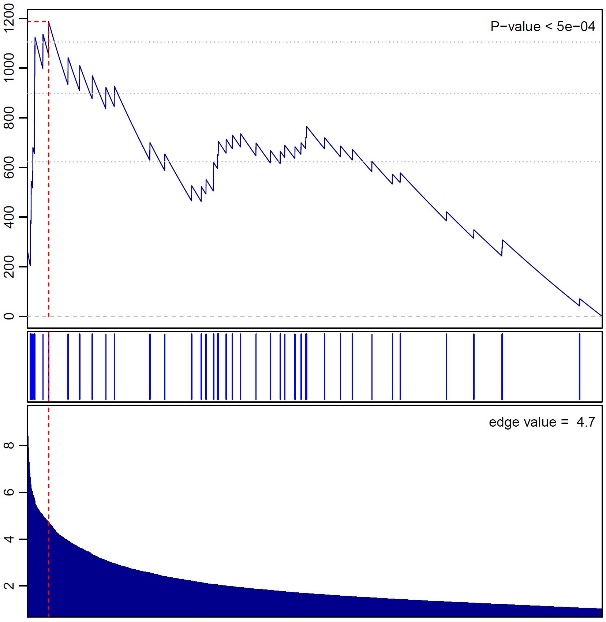

**Sorted Genes (by descending Log2FC)**

**Sorted Genes (by descending Log2FC)**

**Log2FC Values**

**Enrichment Score (cumulative weight)**

**Log2FC Values**

**C**

**C**

**HBSER5 (*q* = 0.042)**

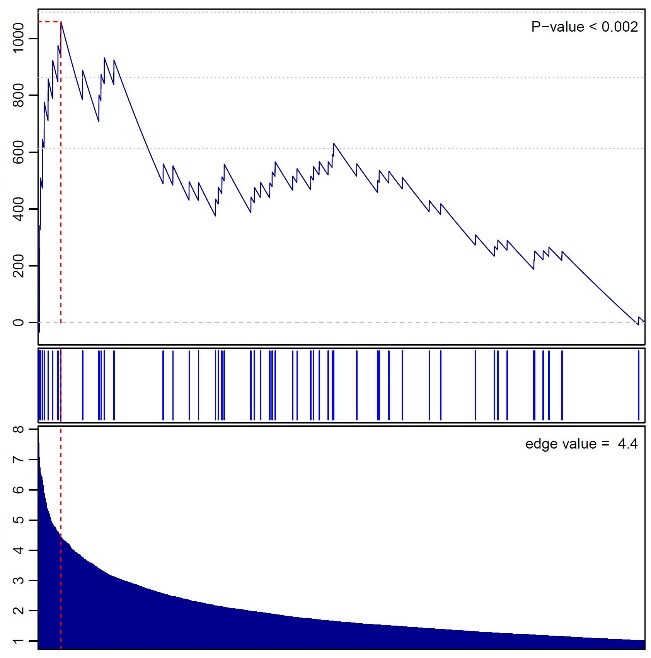

**Enrichment Score (cumulative weight)**

**Log2FC Values**

**Sorted Genes (by descending Log2FC)**

**D**

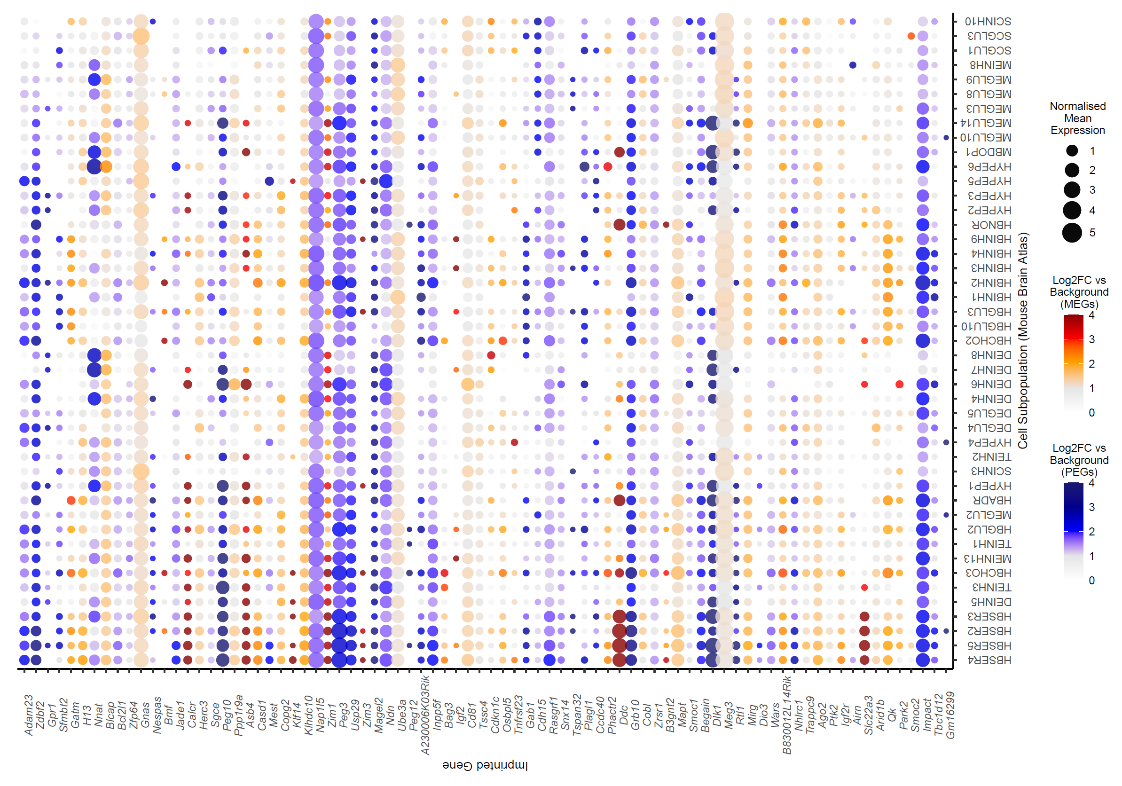

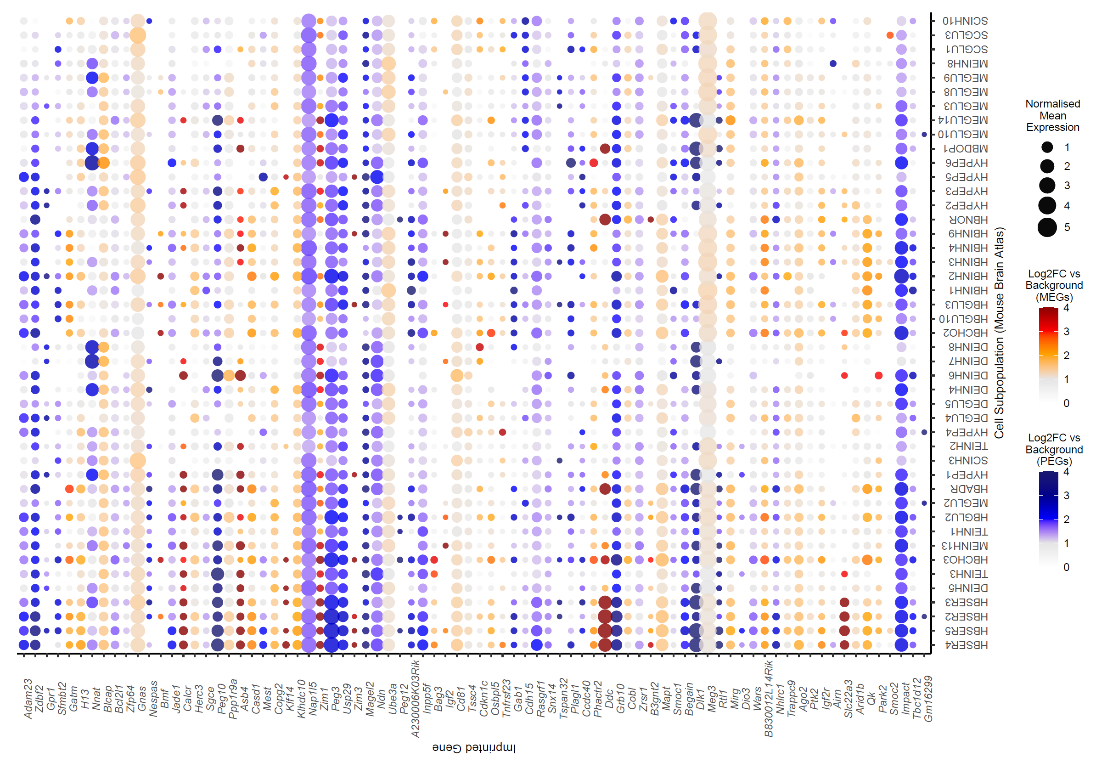

**Supplemental Figure S4.** Significant GSEA for imprinted genes in three subpopulations of Hindbrain Serotonin Neurons (HBSER2, HBSER4, HBSER5) in Mouse Brain Atlas (3). **(***A-C***) -** GSEA graph for the three enriched serotonin neuron populations. See Figure S1 legend for description on how to interpret GSEA analysis and plot. *(D)* – Dot plot of imprinted genes upregulated in any of the 3 enriched serotonin neuron populations plotted across all over-represented neuron subpopulations found in the Mouse Brain Atlas. Imprinted genes were plotted in chromosomal order. Size of points represented absolute mean expression; colour represented the size of the Log2FC value for the cell identity group (e.g., HBSER4) vs. all other cells. Unique colour scales are used for MEGs (red/orange) and PEGs (blue). Where a gene was not expressed in a cell type, this appears as a blank space in the plot

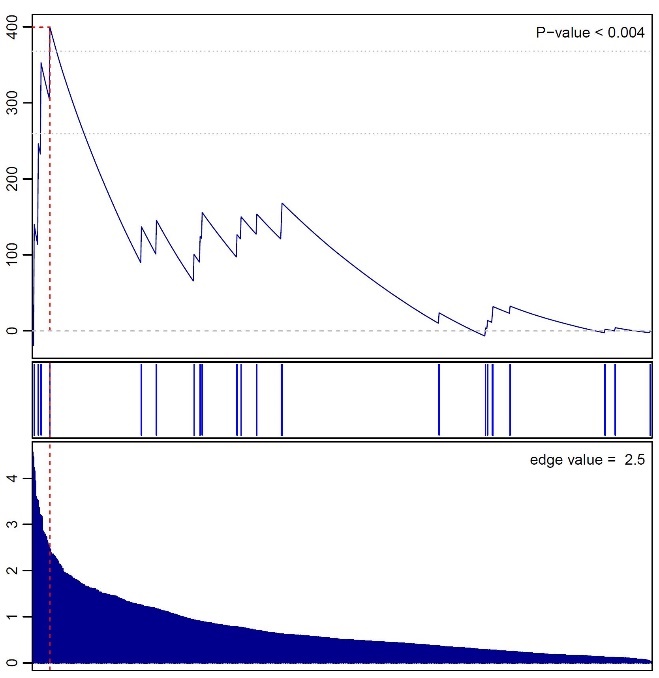

**Enrichment Score (cumulative weight)**

**A**

**Log2FC Values**

**Sorted Genes (by descending Log2FC)**

**B**

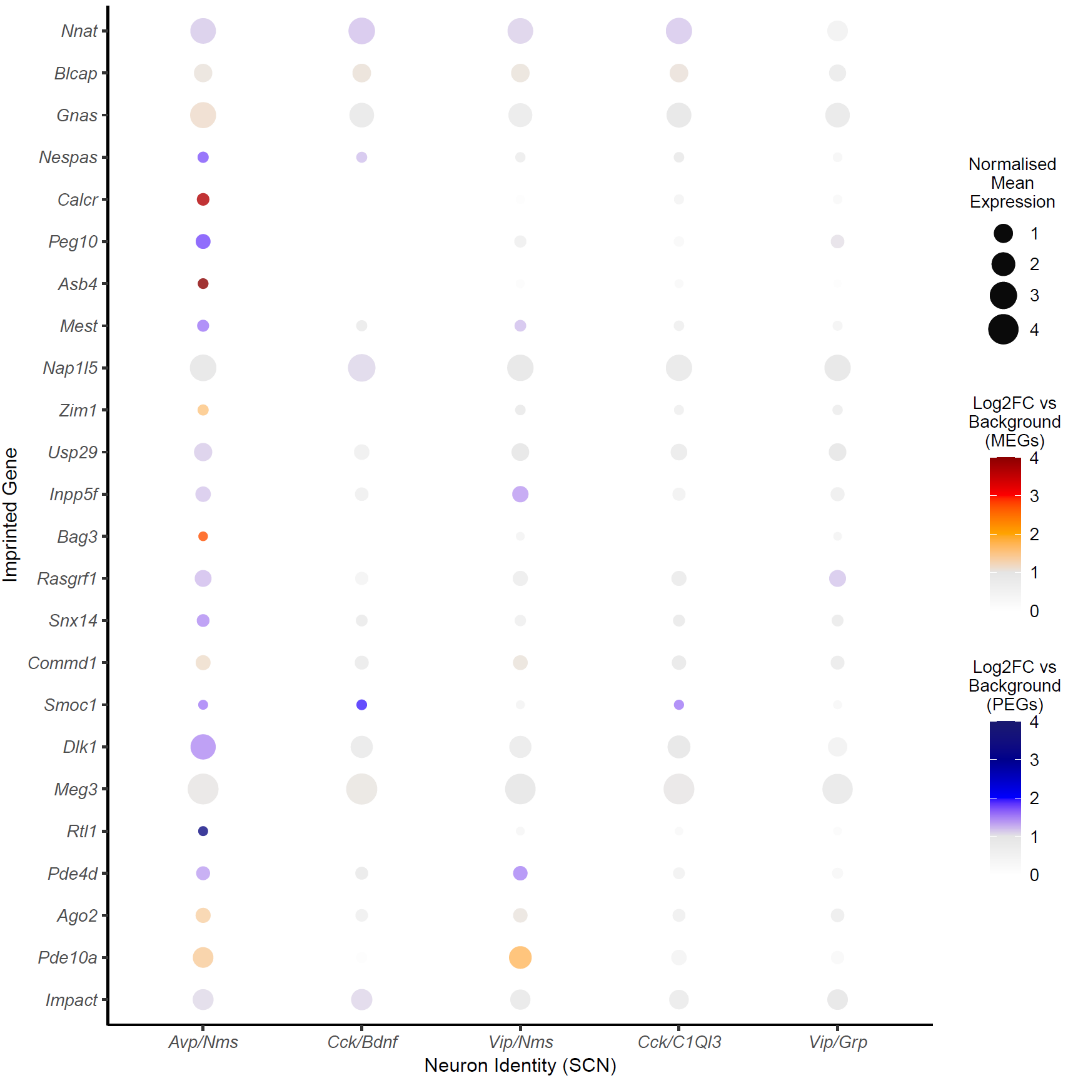

**Supplemental Figure S5.** Significant GSEA for imprinted genes in *Avp/Nms* neurons in Suprachiasmatic Nucleus (9). *(A)* GSEA graph for *Avp/Nms* neurons. See Figure S1 legend for description on how to interpret GSEA analysis and plot. *(B)* – Dot plot of imprinted genes upregulated in *Avp/Nms* neurons plotted across the five neuron types identified in the SCN. See Figure S4D legend for a description of how to interpret the dot plot.

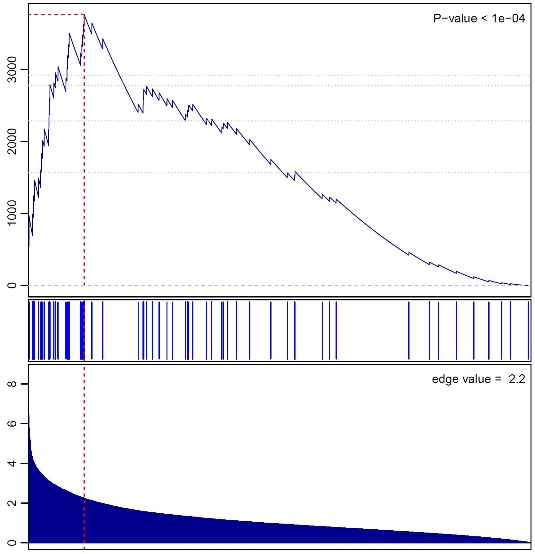

**Sorted Genes (by descending Log2FC)**

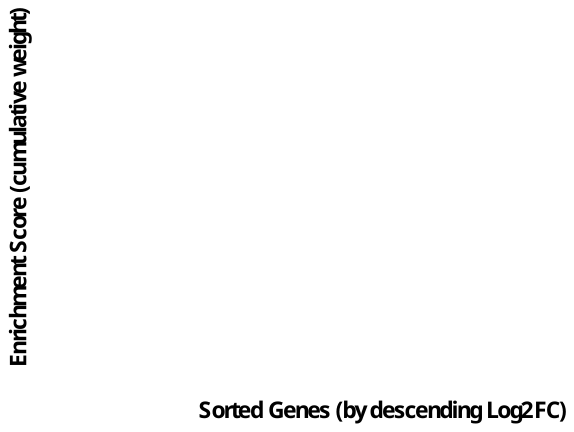

**Enrichment Score (cumulative weight)**

**Log2FC Value**

**Serotonergic Neurons (q = 0.0005)**

**Sorted Genes (by descending Log2FC)**

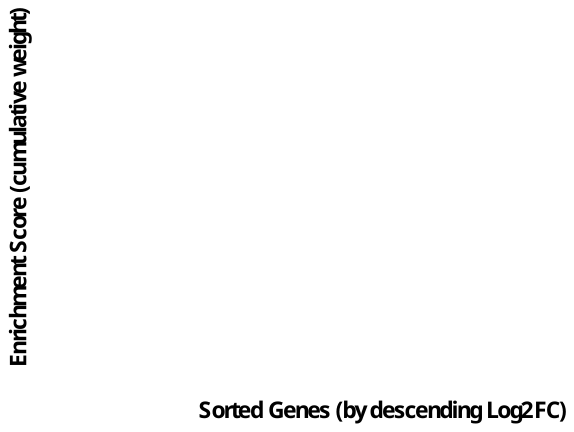

**Enrichment Score (cumulative weight)**

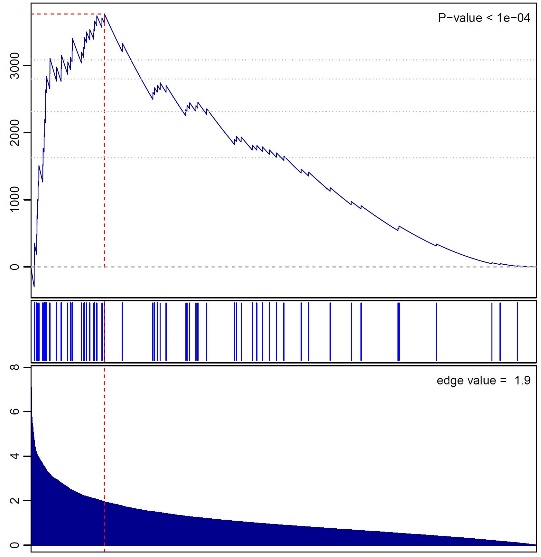

**Log2FC Value**

**Dopaminergic Neurons (*q* = 0.0005)**

**B**

**A**

**Glutamatergic Neurons (*q* = 0.0035)**

**D**

**C**

**GABAergic Neurons (*q* = 0.003)**

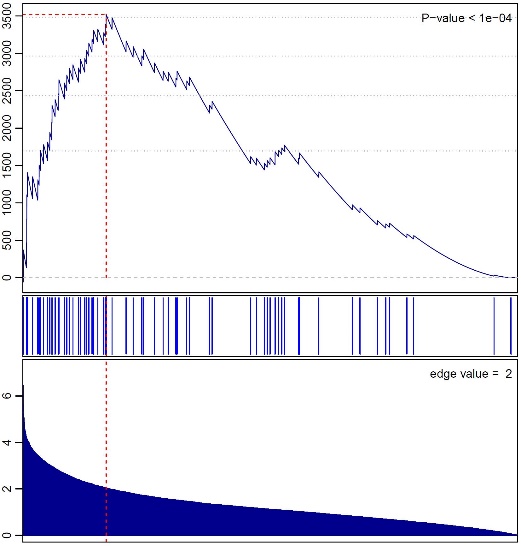

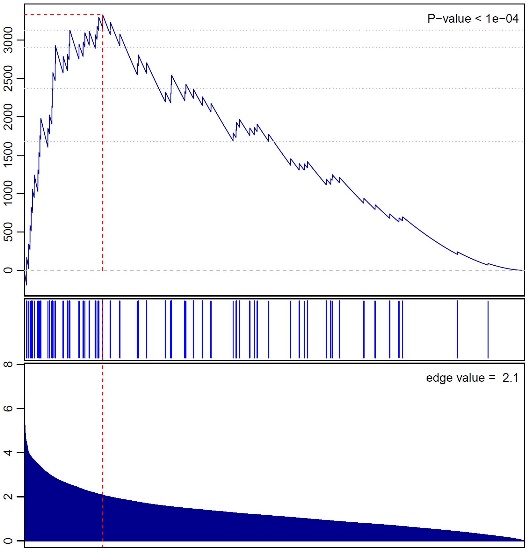

**Sorted Genes (by descending Log2FC)**

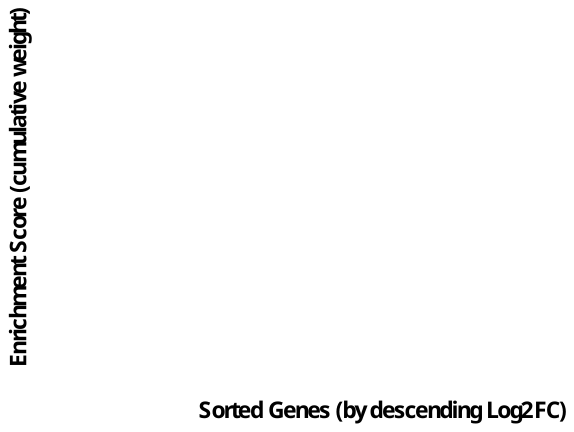

**Enrichment Score (cumulative weight)**

**Log2FC Value**

**Sorted Genes (by descending Log2FC)**

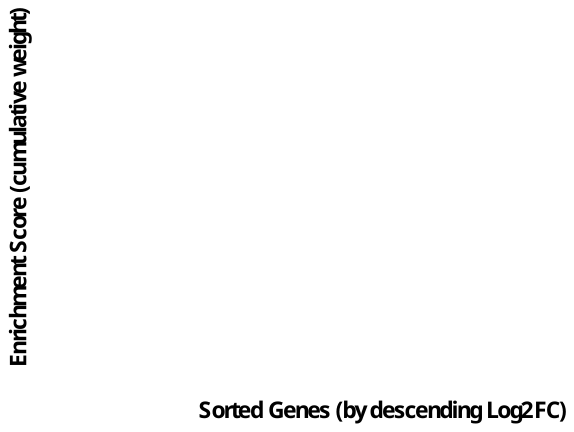

**Enrichment Score (cumulative weight)**

**Log2FC Value**

**F**

**E**

**Peptidergic Neurons (*q* = 0.034)**

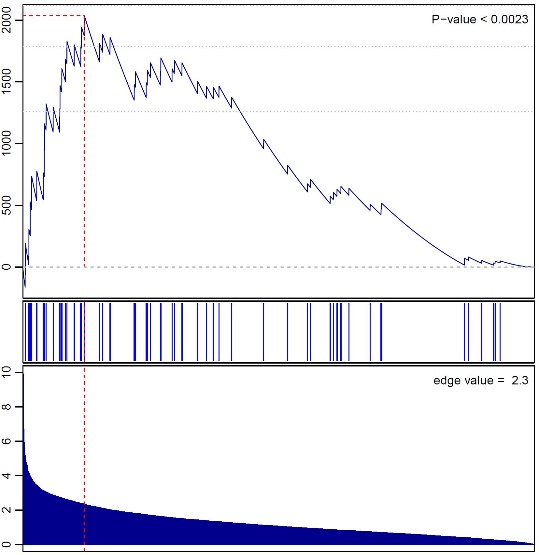

**Sorted Genes (by descending Log2FC)**

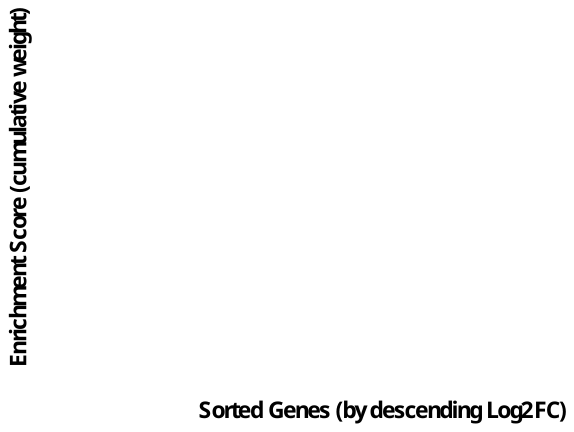

**Enrichment Score (cumulative weight)**

**Log2FC Value**

**F**

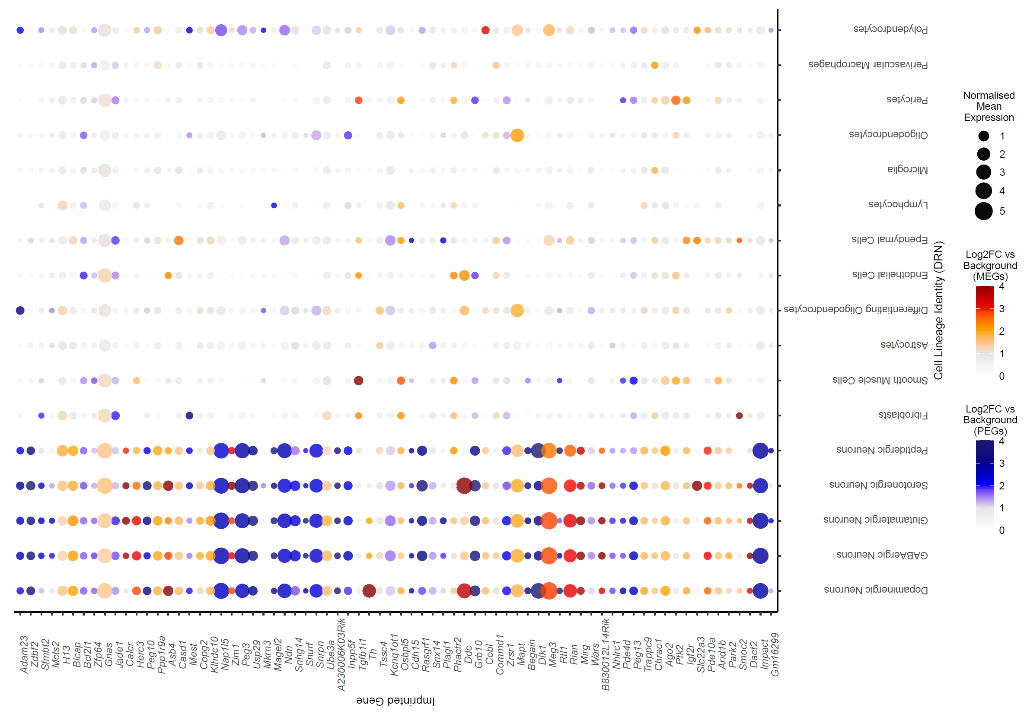

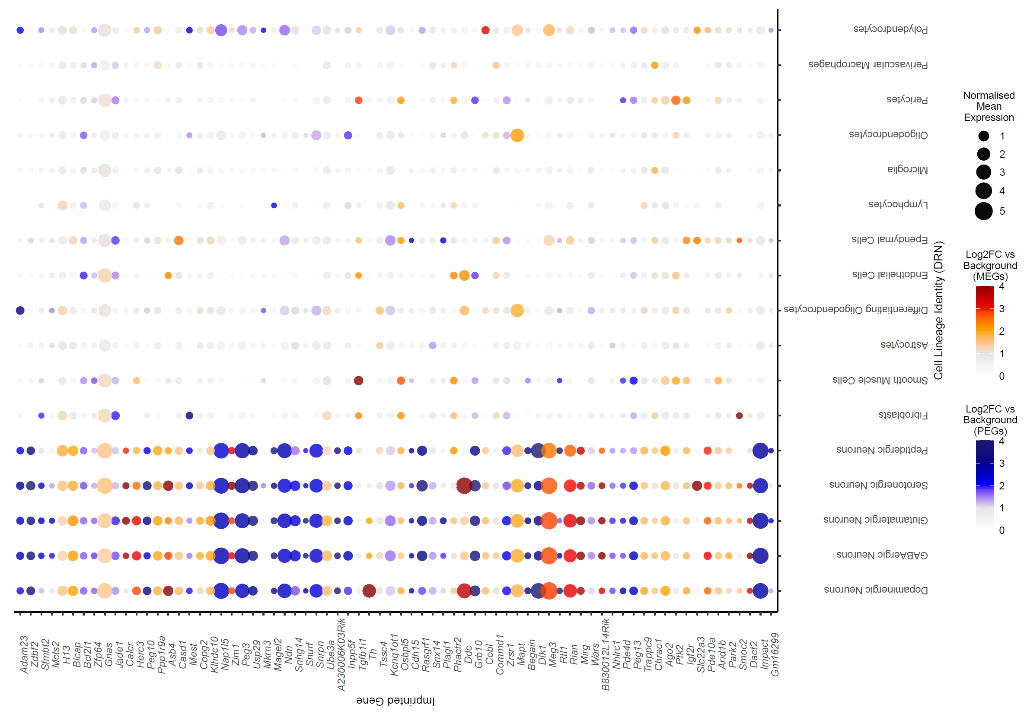

**Supplemental Figure S6.** Significant GSEA for imprinted genes in neuronal types in the Dorsal Raphe Nucleus (10) **(***A-E***)** – GSEA graphs for Serotonin neurons, Dopamine neurons, GABA neurons, Glutamatergic neurons, Peptidergic neurons. See Figure S1 legend for description on how to interpret GSEA analysis and plot. *(F*) Dot plot of the expression of all imprinted genes, upregulated in any of the 5 neuron types, plotted across all cells in the DRN. See Figure S4D legend for details of the plot.

**Supplemental Tables**

**Supplemental Table S1**

Custom List of Imprinted Genes used as the gene set in this study (‘Supplemental_Table_S1_Imprinted_Gene_List.xlsx')

**Supplemental Table S2**

The average normalised expression for all imprinted genes across the identity groups in the datasets for analyses at Levels 1 and 2. Analysis was carried out for the Mouse Cell Atlas (MCA, (1)) and *Tabula Muris* (2) datasets from Level 1 (Multi-Organ Comparison) and the Mouse Brain Atlas (3) and Ximerakis, Lipnick (4) datasets from Level 2 (Whole Brain Comparison). Tables for each dataset are presented in the same spreadsheet. *Tissue/Cell Type/Nervous System Region –* the identity groups that the cells were grouped under. *IG Mean Normalised Expression –* The average normalised expression value for all imprinted genes in cells of that identity group. *Rest Mean Normalised Expression* – The average normalised expression value for all other genes excluding the imprinted genes to provide a baseline expression value. (‘Supplemental_Table_S2_Mean_Normalised_Expression_Level_1_2.xlsx')

**Supplemental Table S3 (A & B)**

| Supplemental Table S3A, Maternally Expressed Genes (MEG) Multi-Organ Level Enrichment Analysis for the Mouse Cell Atlas (MCA, (1)) dataset. *Identity –* Tissue identities for the cells used in analysis; *Up Reg* – number of upregulated genes with *q* ≤ 0.05 and Log2FC ≥ 1 (total number of genes in the dataset in brackets); *MEG* – number of maternally expressed genes upregulated with *q* ≤ 0.05 and Log2FC ≥ 1 (total number of MEGs in the dataset in brackets); *ORA p* – *p* value from over representation analysis on groups with minimum 5% of total MEGs; *ORA q* – Bonferroni corrected *p* value from ORA; *Mean FC MEG* – mean fold change for upregulated MEGs; *Mean FC Rest* – mean fold change for all other upregulated genes; *GSEA p* – *p* value from Gene Set Enrichment Analysis for identity groups with 15+ MEGs and Mean FC MEG > Mean FC Rest; *GSEA q* – Bonferroni corrected *p* values from GSEA; *No. MEGs with highest expression* – Number of MEGs with highest mean expression value for that identity group. | | | | | | | |
| --- | --- | --- | --- | --- | --- | --- | --- |
| Identity | **Up Reg**  **(20,534)** | **MEG (53)** | **ORA *p*** | **ORA *q*** | **Mean FC MEG** | **Mean FC Rest** | **No. MEGs with highest expression** |
| Pancreas | 2737 | 24 | 1.39E-08 | **1.53E-07** | 4.54 | 10.35 | 13 |
| Bladder | 3183 | 19 | 0.0002 | **0.0025** | 4.26 | 8.50 | 5 |
| Kidney | 1714 | 10 | 0.0113 | 0.1245 | 13.76 | 182.89 | 5 |
| Uterus | 2567 | 12 | 0.0281 | 0.3090 | 4.44 | 8.45 | 4 |
| Lung | 1203 | 6 | 0.0884 | 0.9723 | 4.11 | 151.16 | 4 |
| Brain | 3401 | 13 | 0.0887 | 0.9753 | 9.04 | 124.28 | 6 |
| Liver | 1739 | 6 | 0.2909 | 1 | 3.30 | 80.42 | 1 |
| Small Intestine | 1719 | 5 | 0.4596 | 1 | 7.99 | 218.64 | 2 |
| Stomach | 1821 | 5 | 0.5116 | 1 | 4.71 | 88.50 | 2 |
| Ovary | 2219 | 4 | 0.8387 | 1 | 2.73 | 11.26 | 2 |
| Testis | 5212 | 8 | 0.9756 | 1 | 6.87 | 5052.59 | 5 |
| Mammary Gland | 902 | 2 | - | - | 5.35 | 4.02 | 0 |
| Bone Marrow | 1095 | 1 | - | - | 6.36 | 4.43 | 1 |
| Muscle | 1127 | 1 | - | - | 15.81 | 15.03 | 2 |
| Peripheral Blood | 1146 | 1 | - | - | 4.61 | 3.57 | 0 |
| Thymus | 1805 | 1 | - | - | 5.06 | 6.75 | 1 |
| Prostate | 369 | 0 | - | - | 0.00 | 478.10 | 0 |
| Spleen | 1501 | 0 | - | - | 0.00 | 4.77 | 0 |

| Supplemental Table S3B. Paternally Expressed Genes (PEG) Multi-Organ Level Enrichment Analysis for the MCA (1) dataset. *Identity –* Tissue identities for the cells used in analysis; *Up Reg* – number of upregulated genes with *q* ≤ 0.05 and Log2FC ≥ 1 (total number of genes in the dataset in brackets); *PEG* – number of paternally expressed genes upregulated with *q* ≤ 0.05 and Log2FC ≥ 1 (total number of PEGs in the dataset in brackets); *ORA p* – *p* value from over representation analysis on groups with minimum 5% of total PEGs; *ORA q* – Bonferroni corrected *p* value from ORA; *Mean FC PEG* – mean fold change for upregulated PEGs; *Mean FC Rest* – mean fold change for all other upregulated genes; *GSEA p* – *p* value from Gene Set Enrichment Analysis for identity groups with 15+ PEGs and Mean FC PEG > Mean FC Rest; *GSEA q* – Bonferroni corrected *p* values from GSEA; *Top Tissue PEGs* – *No. PEGs with highest expression* – Number of PEGs with highest mean expression value for that identity group | | | | | | | | |
| --- | --- | --- | --- | --- | --- | --- | --- | --- |
| Identity | **Up Reg**  **(20,534)** | **PEG (42)** | **ORA *p*** | **ORA *q*** | **Mean FC PEG** | **Mean FC Rest** | **GSEA *p*** | **No. PEGs with highest expression** |
| Brain | 3401 | 21 | 5.70E-07 | **4.56E-06** | 8.59 | 124.56 | - | 13 |
| Pancreas | 2737 | 18 | 2.41E-06 | **1.93E-05** | 14.33 | 10.27 | **0.0206** | 9 |
| Uterus | 2567 | 10 | 0.0312 | 0.2496 | 4.93 | 8.44 | - | 3 |
| Ovary | 2219 | 9 | 0.0328 | 0.2624 | 9.57 | 11.26 | - | 3 |
| Bladder | 3183 | 10 | 0.1049 | 0.8389 | 4.82 | 8.49 | - | 3 |
| Thymus | 1805 | 5 | 0.3082 | 1 | 2.32 | 6.76 | - | 1 |
| Muscle | 1127 | 3 | 0.4076 | 1 | 6.25 | 15.05 | - | 1 |
| Testis | 5212 | 6 | 0.9730 | 1 | 53.94 | 5050.60 | - | 5 |
| Liver | 1739 | 2 | - | - | 8.32 | 80.24 | - | 2 |
| Lung | 1203 | 2 | - | - | 2.96 | 150.68 | - | 0 |
| Mammary Gland | 902 | 2 | - | - | 2.06 | 4.03 | - | 0 |
| Peripheral Blood | 1146 | 2 | - | - | 3.37 | 3.57 | - | 0 |
| Stomach | 1821 | 2 | - | - | 3.07 | 88.37 | - | 1 |
| Bone Marrow | 1095 | 1 | - | - | 4.26 | 4.43 | - | 0 |
| Spleen | 1501 | 1 | - | - | 4.90 | 4.77 | - | 1 |
| Kidney | 1714 | 0 | - | - | 0.00 | 181.90 | - | 0 |
| Prostate | 369 | 0 | - | - | 0.00 | 478.10 | - | 0 |
| Small Intestine | 1719 | 0 | - | - | 0.00 | 218.03 | - | 0 |

**Supplemental Table S4 (A & B)**

| Supplemental Table S4A. MEG Multi-Organ Level Enrichment Analysis for the *Tabula Muris* (2). For all column descriptions, please see the legend of Supplemental Table S3A. | | | | | | | |
| --- | --- | --- | --- | --- | --- | --- | --- |
| Identity | **Up Reg**  **(20,839)** | **MEG**  **(58)** | **ORA**  ***p*** | **ORA**  ***q*** | **Mean FC**  **MEG** | **Mean FC**  **Rest** | **No. MEGs with highest expression** |
| Diaphragm | 416 | 12 | 1.33E-09 | **2.13E-08** | 5.28 | 4.89 | 3 |
| Limb Muscle | 761 | 14 | 1.52E-08 | **2.43E-07** | 8.15 | 5.16 | 4 |
| Trachea | 1979 | 15 | 0.0003 | **0.0040** | 3.22 | 4.57 | 3 |
| Bladder | 3338 | 19 | 0.0012 | **0.0198** | 2.98 | 5.30 | 10 |
| Pancreas | 4104 | 20 | 0.0059 | 0.0938 | 12.20 | 12.60 | 14 |
| Heart | 1108 | 7 | 0.0334 | 0.5346 | 3.03 | 5.13 | 0 |
| Mammary Gland | 1826 | 8 | 0.1321 | 1 | 3.36 | 5.24 | 2 |
| Fat | 1263 | 6 | 0.1378 | 1 | 2.47 | 3.69 | 0 |
| Brain (Non-Myeloid) | 3081 | 12 | 0.1401 | 1 | 4.99 | 14.19 | 4 |
| Brain (Myeloid) | 1024 | 5 | 0.1552 | 1 | 3.39 | 6.80 | 2 |
| Kidney | 584 | 3 | 0.2217 | 1 | 24.94 | 22.90 | 1 |
| Tongue | 4295 | 11 | 0.6725 | 1 | 4.42 | 7.16 | 6 |
| Aorta | 3515 | 8 | 0.7845 | 1 | 7.34 | 16.07 | 1 |
| Skin | 1612 | 3 | 0.8365 | 1 | 3.88 | 8.36 | 1 |
| Large Intestine | 4758 | 10 | 0.8822 | 1 | 6.34 | 12.22 | 4 |
| Liver | 1808 | 3 | 0.8894 | 1 | 4.88 | 54.82 | 1 |
| Lung | 914 | 2 | - | - | 2.73 | 5.41 | 0 |
| Marrow | 1957 | 2 | - | - | 10.31 | 5.25 | 2 |
| Spleen | 625 | 0 | - | - | 0.00 | 4.28 | 0 |
| Thymus | 678 | 0 | - | - | 0.00 | 7.42 | 0 |

| Supplemental Table S4B. PEG Multi-Organ Level Enrichment Analysis for the *Tabula Muris* (2). For all column descriptions, please see the legend of Supplemental Table S3B. | | | | | | | | | |
| --- | --- | --- | --- | --- | --- | --- | --- | --- | --- |
| Identity | **Up Reg**  **(20,839)** | **PEG**  **(49)** | **ORA**  ***p*** | **ORA**  ***q*** | **Mean FC**  **PEG** | **Mean FC**  **Rest** | **GSEA**  ***p*** | **GSEA**  ***q*** | **No. PEGs with highest expression** |
| Limb_Muscle | 761 | 10 | 8.98E-06 | **0.0001** | 10.25 | 5.15 | - | - | 4 |
| Pancreas | 4104 | 23 | 1.51E-05 | **0.0002** | 12.80 | 12.60 | 0.1283 | 0.2566 | 15 |
| Brain (Non-Myeloid) | 3081 | 19 | 3.42E-05 | **0.0005** | 16.69 | 14.13 | 0.3785 | 0.757 | 10 |
| Diaphragm | 416 | 7 | 5.01E-05 | **0.0007** | 8.57 | 4.84 | - | - | 1 |
| Trachea | 1979 | 10 | 0.0154 | 0.2150 | 4.69 | 4.56 | - | - | 2 |
| Fat | 1263 | 6 | 0.0744 | 1 | 4.46 | 3.68 | - | - | 1 |
| Thymus | 678 | 4 | 0.0746 | 1 | 3.46 | 7.45 | - | - | 1 |
| Bladder | 3338 | 12 | 0.0823 | 1 | 3.81 | 5.29 | - | - | 6 |
| Heart | 1108 | 3 | 0.4872 | 1 | 2.50 | 5.12 | - | - | 0 |
| Liver | 1808 | 4 | 0.6247 | 1 | 7.17 | 54.84 | - | - | 2 |
| Mammary Gland | 1826 | 4 | 0.6327 | 1 | 3.83 | 5.23 | - | - | 1 |
| Marrow | 1957 | 3 | 0.8512 | 1 | 5.88 | 5.25 | - | - | 2 |
| Aorta | 3515 | 6 | 0.8564 | 1 | 7.63 | 16.06 | - | - | 1 |
| Tongue | 4295 | 4 | 0.9949 | 1 | 3.40 | 7.16 | - | - | 2 |
| Large Intestine | 4758 | 1 | - | - | 2.11 | 12.21 | - | - | 1 |
| Skin | 1612 | 1 | - | - | 4.94 | 8.35 | - | - | 0 |
| Spleen | 625 | 1 | - | - | 4.63 | 4.28 | - | - | 0 |
| Brain (Myeloid) | 1024 | 0 | - | - | 0.00 | 6.78 | - | - | 0 |
| Kidney | 584 | 0 | - | - | 0.00 | 22.91 | - | - | 0 |
| Lung | 914 | 0 | - | - | 0.00 | 5.40 | - | - | 0 |

**Supplemental Table S5 (A & B)**

| Supplemental Table S5A. MEG over-representation in neural lineage types (4). *Identity –* Cell lineage identities for the cells used in analysis. All other column descriptions can be found in the legend of Supplemental Table S3A | | | | | | | | | |
| --- | --- | --- | --- | --- | --- | --- | --- | --- | --- |
| Identity (Abbr.) | **Up Reg**  **(14,498)** | **MEG**  **(45)** | **ORA *p*** | **ORA *q*** | **Mean FC**  **MEG** | **Mean FC**  **Rest** | **GSEA *p*** | **GSEA *q*** | **No. MEGs with highest expression** |
| Arachnoid barrier cells (ABC) | 2287 | 16 | 0.0009 | **0.0139** | 19.04 | 22.60 | - | - | 5 |
| Neuroendocrine cells (NendC) | 3868 | 21 | 0.0031 | **0.0466** | 6.30 | 5.49 | 0.3804 | 0.7608 | 10 |
| Pericytes (PC) | 1801 | 11 | 0.0194 | 0.2910 | 7.84 | 8.23 | - | - | 0 |
| Mature Neurons (all types) (mNEUR) | 2968 | 15 | 0.0301 | 0.4519 | 12.06 | 9.26 | 0.2256 | 0.4512 | 2 |
| Astrocyte-restricted precursors (ARP) | 1445 | 6 | 0.2895 | 1 | 4.47 | 5.09 | - | - | 1 |
| Choroid plexus epithelial cells (CPC) | 2602 | 12 | 0.0954 | 1 | 8.10 | 19.31 | - | - | 5 |
| Endothelial cells (EC) | 1455 | 6 | 0.2950 | 1 | 6.11 | 8.54 | - | - | 0 |
| Ependymocytes (EPC) | 3233 | 7 | 0.9022 | 1 | 4.58 | 53.18 | - | - | 2 |
| Hemoglobin-expressing vascular cells (Hb_VC) | 1798 | 7 | 0.3218 | 1 | 6.20 | 6.32 | - | - | 1 |
| Hypendymal cells (HypEPC) | 1525 | 3 | 0.8660 | 1 | 9.59 | 20.81 | - | - | 3 |
| Oligodendrocytes (OLG) | 1183 | 5 | 0.3044 | 1 | 3.82 | 12.88 | - | - | 3 |
| Oligodendrocyte precursor cells (OPC) | 1524 | 6 | 0.3339 | 1 | 3.11 | 7.15 | - | - | 1 |
| Tanycytes (TNC) | 1279 | 5 | 0.3648 | 1 | 6.08 | 11.98 | - | - | 3 |
| Vascular and leptomeningeal cells (VLMC) | 1714 | 9 | 0.0775 | 1 | 20.02 | 13.01 | - | - | 1 |
| Vascular smooth muscle cells (VSMC) | 3006 | 10 | 0.4608 | 1 | 12.77 | 6.71 | - | - | 4 |
| Astrocytes (ASC) | 1384 | 1 | - | - | 2.38 | 6.04 | - | - | 0 |
| Dendritic cells (DC) | 1209 | 1 | - | - | 3.50 | 16.02 | - | - | 1 |
| Immature Neurons (ImmN) | 652 | 1 | - | - | 3.20 | 5.78 | - | - | 0 |
| Macrophages (MAC) | 1222 | 2 | - | - | 3.47 | 21.56 | - | - | 0 |
| Microglia (MG) | 1342 | 2 | - | - | 26.89 | 19.21 | - | - | 3 |
| Monocytes (MNC) | 947 | 0 | - | - | 0.00 | 19.12 | - | - | 0 |
| Neutrophils (NEUT) | 519 | 0 | - | - | 0.00 | 61.92 | - | - | 0 |
| Neuronal-restricted precursor (NRP) | 2339 | 2 | - | - | 2.51 | 10.18 | - | - | 0 |
| Neural stem cells (NSC) | 1009 | 1 | - | - | 2.11 | 4.09 | - | - | 0 |
| Olfactory ensheathing glia (OEG) | 1086 | 2 | - | - | 12.12 | 25.90 | - | - | 0 |

| Supplemental Table S5B. PEG over-representation in neural lineage types (4). *Identity –* Cell lineage identities for the cells used in analysis. All other column descriptions can be found in the legend of Supplemental Table S3B | | | | | | | | |
| --- | --- | --- | --- | --- | --- | --- | --- | --- |
| Identity (Abbr.) | **Up Reg**  **(14,498)** | **PEG**  **(40)** | **ORA *p*** | **ORA *q*** | **Mean FC**  **PEG** | **Mean FC**  **Rest** | **GSEA *p*** | **No. PEGs with highest expression** |
| Neuroendocrine cells (NendC) | 3868 | 26 | 4.27E-07 | **8.97E-06** | 16.39 | 5.42 | 4.00E-04 | 16 |
| Mature Neurons (all types) (mNEUR) | 2968 | 17 | 0.0013 | **0.0269** | 5.92 | 9.30 | - | 0 |
| Olfactory ensheathing glia (OEG) | 1086 | 7 | 0.0274 | 0.5757 | 6.76 | 26.00 | - | 1 |
| Tanycytes (TNC) | 1279 | 7 | 0.0583 | 1 | 7.04 | 11.99 | - | 5 |
| Oligodendrocyte precursor cells (OPC) | 1524 | 7 | 0.1211 | 1 | 2.97 | 7.15 | - | 0 |
| Immature Neurons (ImmN) | 652 | 3 | 0.2677 | 1 | 3.42 | 5.79 | - | 0 |
| Neural stem cells (NSC) | 1009 | 4 | 0.3025 | 1 | 4.48 | 4.08 | - | 0 |
| Neuronal-restricted precursor (NRP) | 2339 | 8 | 0.3125 | 1 | 3.21 | 10.20 | - | 1 |
| Vascular and leptomeningeal cells (VLMC) | 1714 | 6 | 0.3332 | 1 | 7.63 | 13.07 | - | 3 |
| Ependymocytes (EPC) | 3233 | 10 | 0.3998 | 1 | 22.37 | 53.17 | - | 2 |
| Oligodendrocytes (OLG) | 1183 | 4 | 0.4141 | 1 | 3.60 | 12.87 | - | 2 |
| Neutrophils (NEUT) | 519 | 2 | 0.4220 | 1 | 9.18 | 62.13 | - | 0 |
| Monocytes (MNC) | 947 | 2 | 0.7459 | 1 | 16.49 | 19.13 | - | 1 |
| Hemoglobin-expressing vascular cells (Hb_VC) | 1798 | 4 | 0.7482 | 1 | 3.58 | 6.33 | - | 2 |
| Astrocytes (ASC) | 1384 | 3 | 0.7490 | 1 | 2.16 | 6.04 | - | 0 |
| Vascular smooth muscle cells (VSMC) | 3006 | 7 | 0.7517 | 1 | 3.46 | 6.73 | - | 1 |
| Hypendymal cells (HypEPC) | 1525 | 3 | 0.8071 | 1 | 24.89 | 20.78 | - | 2 |
| Choroid plexus epithelial cells (CPC) | 2602 | 5 | 0.8686 | 1 | 5.83 | 19.28 | - | 0 |
| Pericytes (PC) | 1801 | 3 | 0.8893 | 1 | 9.50 | 8.22 | - | 2 |
| Arachnoid barrier cells (ABC) | 2287 | 4 | 0.8954 | 1 | 8.04 | 22.60 | - | 2 |
| Astrocyte-restricted precursors (ARP) | 1445 | 2 | 0.9189 | 1 | 4.57 | 5.08 | - | 0 |
| Dendritic cells (DC) | 1209 | 0 | - | - | 0.00 | 16.01 | - | 0 |
| Endothelial cells (EC) | 1455 | 1 | - | - | 3.97 | 8.53 | - | 0 |
| Macrophages (MAC) | 1222 | 0 | - | - | 0.00 | 21.53 | - | 0 |
| Microglia (MG) | 1342 | 1 | - | - | 4.06 | 19.23 | - | 0 |

**Supplemental Table S6 (A & B)**

| Supplemental Table S6A. MEG over-representation in nervous system cell types (3). *Identity –* Cell identities for the cells used in analysis. All other column descriptions can be found in the legend of Supplemental Table S3A. | | | | | | | |
| --- | --- | --- | --- | --- | --- | --- | --- |
| Identity | **Up Reg**  **(19,547)** | **MEG**  **(59)** | **ORA *p*** | **ORA *q*** | **Mean FC**  **MEG** | **Mean FC**  **Rest** | **No. MEGs with highest expression** |
| Vascular | 2473 | 19 | 7.37E-05 | **0.0004** | 18.52 | 26.62 | 14 |
| Neurons | 5710 | 22 | 0.1122 | 0.6730 | 12.60 | 24.91 | 21 |
| Ependymal | 3683 | 13 | 0.3128 | 1 | 4.14 | 66.96 | 9 |
| Immune | 1564 | 5 | 0.5150 | 1 | 15.51 | 92.94 | 3 |
| Oligos | 1587 | 5 | 0.5283 | 1 | 4.42 | 11.45 | 5 |
| Peripheral Glia | 2820 | 8 | 0.6316 | 1 | 5.57 | 12.62 | 5 |
| Astrocytes | 1539 | 2 | - | - | 3.14 | 10.72 | 2 |

| Supplemental Table S6B. PEG over-representation in nervous system cell types (3). *Identity –* Cell identities for the cells used in analysis. All other column descriptions can be found in the legend of Supplemental Table S3B. | | | | | | | |
| --- | --- | --- | --- | --- | --- | --- | --- |
| Identity | **Up Reg**  **(19,547)** | **PEG**  **(49)** | **ORA *p*** | **ORA *q*** | **Mean FC**  **PEG** | **Mean FC**  **Rest** | **No. PEGs with highest expression** |
| Neurons | 5710 | 22 | 0.0184 | 0.0919 | 10.87 | 24.92 | 24 |
| Oligos | 1587 | 6 | 0.2175 | 1 | 4.82 | 11.45 | 7 |
| Peripheral Glia | 2820 | 8 | 0.4353 | 1 | 5.28 | 12.62 | 7 |
| Ependymal | 3683 | 7 | 0.8565 | 1 | 62.36 | 66.75 | 6 |
| Vascular | 2473 | 3 | 0.9610 | 1 | 14.06 | 26.57 | 2 |
| Astrocytes | 1539 | 2 | - | - | 2.63 | 10.72 | 1 |
| Immune | 1564 | 2 | - | - | 8.21 | 92.80 | 2 |

**Supplemental Table S7 (A & B)**

| Supplemental Table S7A. MEG over-representation in nervous system regions (3). *Identity –* Nervous system regional identities for the cells used in analysis genes. All other column descriptions can be found in the legend of Supplemental Table S3A. | | | | | | | | | |
| --- | --- | --- | --- | --- | --- | --- | --- | --- | --- |
| Identity | **Up Reg**  **(18,335)** | **MEG**  **(57)** | **ORA *p*** | **ORA *q*** | **Mean FC**  **MEG** | **Mean FC**  **Rest** | **GSEA**  ***p*** | **GSEA**  ***q*** | **No. MEGs with highest expression** |
| Medulla | 3147 | 21 | 0.0003 | **0.0048** | 5.70 | 4.01 | 0.0318 | 0.0636 | 9 |
| Pons | 3581 | 20 | 0.0042 | 0.0680 | 5.40 | 3.90 | 0.0136 | **0.0272** | 10 |
| Vent. Midbrain | 1228 | 8 | 0.0352 | 0.5631 | 5.95 | 4.98 | - | - | 1 |
| Hypothalamus | 1040 | 7 | 0.0414 | 0.6625 | 6.01 | 5.82 | - | - | 2 |
| Middle Cortex | 623 | 5 | 0.0439 | 0.7017 | 3.29 | 3.24 | - | - | 0 |
| Enteric Nervous System | 3885 | 17 | 0.0793 | 1 | 9.97 | 120.77 | - | - | 7 |
| Sympathetic Nervous System | 2804 | 12 | 0.1523 | 1 | 14.88 | 57.85 | - | - | 6 |
| Vent. Striatum | 689 | 4 | 0.1659 | 1 | 4.54 | 4.91 | - | - | 0 |
| Anterior Cortex | 979 | 5 | 0.1873 | 1 | 2.82 | 3.30 | - | - | 1 |
| Somatosensory Cortex | 2121 | 9 | 0.2085 | 1 | 4.46 | 3.70 | - | - | 7 |
| Posterior Cortex | 1090 | 5 | 0.2499 | 1 | 2.35 | 3.20 | - | - | 0 |
| Hippocampus - CA1 | 1082 | 4 | 0.4362 | 1 | 3.48 | 4.02 | - | - | 2 |
| Thalamus | 1441 | 5 | 0.4679 | 1 | 3.33 | 6.35 | - | - | 0 |
| Dors. Striatum | 1196 | 4 | 0.5149 | 1 | 4.41 | 5.43 | - | - | 2 |
| Dorsal Root Ganglion | 3607 | 11 | 0.5814 | 1 | 13.29 | 75.79 | - | - | 5 |
| Spinal Cord | 972 | 3 | 0.5882 | 1 | 5.53 | 12.34 | - | - | 0 |
| Olfactory Bulb | 445 | 2 | - | - | 3.92 | 8.25 | - | - | 1 |
| Antero-Middle Cortex | 646 | 1 | - | - | 5.97 | 4.31 | - | - | 1 |
| Dentate Gyrus | 796 | 1 | - | - | 7.62 | 4.16 | - | - | 1 |
| Hippocampus | 631 | 1 | - | - | 4.36 | 3.83 | - | - | 0 |
| Amygdala | 452 | 0 | - | - | 0.00 | 4.11 | - | - | 0 |
| Cerebellum | 240 | 0 | - | - | 0.00 | 32.30 | - | - | 0 |
| Dors. Midbrain | 1045 | 0 | - | - | 0.00 | 4.83 | - | - | 2 |

| Supplemental Table S7B. PEG over-representation in nervous system regions (3). *Identity –* Nervous system regional identities for the cells used in analysis genes. All other column descriptions can be found in the legend of Supplemental Table S3B. | | | | | | | |
| --- | --- | --- | --- | --- | --- | --- | --- |
| Identity | **Up Reg**  **(18,335)** | **PEG**  **(48)** | **ORA *p*** | **ORA *q*** | **Mean FC**  **PEG** | **Mean FC**  **Rest** | **No. PEGs with highest expression** |
| Hypothalamus | 1040 | 15 | 4.67E-08 | **6.53E-07** | 4.42 | 5.85 | 6 |
| Medulla | 3147 | 24 | 2.93E-07 | **4.10E-06** | 4.00 | 4.03 | 6 |
| Pons | 3581 | 24 | 3.32E-06 | **4.65E-05** | 3.21 | 3.91 | 12 |
| Vent. Midbrain | 1228 | 10 | 0.0013 | **0.0182** | 4.06 | 4.99 | 2 |
| Amygdala | 452 | 4 | 0.0323 | 0.4522 | 4.65 | 4.11 | 2 |
| Dors. Midbrain | 1045 | 6 | 0.0586 | 0.8206 | 2.20 | 4.85 | 1 |
| Vent. Striatum | 689 | 4 | 0.1115 | 1 | 3.31 | 4.92 | 0 |
| Hippocampus | 631 | 3 | 0.2376 | 1 | 5.03 | 3.82 | 2 |
| Posterior Cortex | 1090 | 4 | 0.3326 | 1 | 3.01 | 3.20 | 2 |
| Dentate Gyrus | 796 | 3 | 0.3585 | 1 | 2.52 | 4.17 | 1 |
| Enteric Nervous System | 3885 | 9 | 0.7386 | 1 | 7.12 | 120.55 | 4 |
| Thalamus | 1441 | 3 | 0.7515 | 1 | 2.19 | 6.35 | 0 |
| Sympathetic Nervous System | 2804 | 6 | 0.7808 | 1 | 4.37 | 57.78 | 3 |
| Dorsal Root Ganglion | 3607 | 5 | 0.9758 | 1 | 7.74 | 75.70 | 4 |
| Hippocampus - CA1 | 1082 | 2 | - | - | 2.06 | 4.02 | 0 |
| Antero-Middle Cortex | 646 | 2 | - | - | 3.91 | 4.31 | 0 |
| Olfactory Bulb | 445 | 2 | - | - | 4.12 | 8.25 | 1 |
| Spinal Cord | 972 | 2 | - | - | 3.13 | 12.33 | 1 |
| Somatosensory Cortex | 2121 | 2 | - | - | 2.45 | 3.71 | 1 |
| Dors. Striatum | 1196 | 2 | - | - | 3.26 | 5.43 | 0 |
| Anterior Cortex | 979 | 1 | - | - | 2.24 | 3.30 | 0 |
| Cerebellum | 240 | 0 | - | - | 0.00 | 32.30 | 0 |
| Middle Cortex | 623 | 0 | - | - | 0.00 | 3.24 | 0 |

**Supplemental Table S8**

| Supplemental Table S8. Imprinted gene over-representation in neuronal subpopulations in the MBA dataset (3). *Identity –* Neuronal Subpopulation identifier for the cells used in analysis genes; *Neuron Description –* more specific classification of neuronal subpopulations as described by Zeisel, Hochgerner (3); *Up Reg* – number of upregulated genes with *q* ≤ 0.05 and Log2FC ≥ 1 (total number of genes in the dataset in brackets); *IG* – number of imprinted genes upregulated with *q* ≤ 0.05 and Log2FC ≥ 1 (total number of IGs in the dataset in brackets); *ORA p* – *p* value from over representation analysis on groups with minimum 5% of total IGs; *ORA q* – Bonferroni corrected *p* value from ORA; *Mean FC IG* – mean fold change for upregulated imprinted genes; *Mean FC Rest* – mean fold change for all other upregulated genes; *GSEA p* – *p* value from Gene Set Enrichment Analysis for identity groups with 15+ IGs and Mean FC IG > Mean FC Rest; *GSEA q* – Bonferroni corrected *p* values from GSEA. | | | | | | | | | |
| --- | --- | --- | --- | --- | --- | --- | --- | --- | --- |
| Identity | **Neuron**  **Description** | **Up Reg**  **(18,335)** | **IG**  **(105)** | **ORA *p*** | **ORA *q*** | **Mean FC**  **IG** | **Mean FC**  **Rest** | **GSEA**  ***p*** | **GSEA**  ***q*** |
| HBSER5 | Serotonergic neurons, hindbrain | 3721 | 53 | 8.29E-12 | **1.19E-09** | 13.76 | 5.98 | 0.002 | **0.042** |
| DEINH5 | Peptidergic neurons, hypothalamus | 611 | 19 | 1.94E-09 | **2.79E-07** | 7.47 | 5.48 | 0.0639 | 1 |
| TEINH3 | Inhibitory neurons, telencephalon | 885 | 22 | 5.50E-09 | **7.93E-07** | 8.60 | 6.38 | 0.2076 | 1 |
| MEINH13 | Inhibitory neurons, midbrain | 1229 | 25 | 2.35E-08 | **3.38E-06** | 9.85 | 7.30 | 0.1213 | 1 |
| HBSER4 | Serotonergic neurons, hindbrain | 3510 | 45 | 2.83E-08 | **4.08E-06** | 27.93 | 6.29 | 5.00E-04 | **0.0105** |
| HBGLU2 | Excitatory neurons, hindbrain | 2533 | 37 | 3.33E-08 | **4.80E-06** | 4.08 | 4.88 | - | - |
| HBCHO3 | Afferent nuclei of cranial nerves VI-XII | 4647 | 53 | 4.03E-08 | **5.81E-06** | 7.61 | 5.64 | 0.1175 | 1 |
| HYPEP1 | Peptidergic neurons, hypothalamus | 766 | 19 | 7.34E-08 | **1.06E-05** | 9.92 | 9.46 | 0.2588 | 1 |
| DEINH6 | Peptidergic neurons, hypothalamus | 466 | 15 | 7.43E-08 | **1.07E-05** | 16.64 | 18.97 | - | - |
| HBSER3 | Serotonergic neurons, hindbrain | 2154 | 33 | 8.19E-08 | **1.18E-05** | 12.84 | 6.30 | 0.0127 | 0.2667 |
| HBGLU3 | Excitatory neurons, hindbrain | 1438 | 26 | 1.24E-07 | **1.78E-05** | 5.62 | 6.24 | - | - |
| MBDOP1 | Dopaminergic neurons, periaqueductal grey | 639 | 17 | 1.45E-07 | **2.09E-05** | 12.07 | 16.84 | - | - |
| HYPEP3 | Peptidergic neurons, hypothalamus | 814 | 19 | 1.89E-07 | **2.72E-05** | 4.25 | 7.82 | - | - |
| DEINH4 | Inhibitory neurons, thalamus | 913 | 19 | 1.08E-06 | **0.0002** | 4.20 | 10.29 | - | - |
| TEINH2 | Inhibitory neurons, septal nucleus | 511 | 14 | 1.44E-06 | **0.0002** | 6.32 | 7.45 | - | - |
| HBSER2 | Serotonergic neurons, hindbrain | 3497 | 41 | 2.00E-06 | **0.0003** | 10.24 | 5.32 | 0.0022 | **0.0462** |
| HBINH9 | Inhibitory neurons, hindbrain | 1461 | 24 | 2.34E-06 | **0.0003** | 4.62 | 7.17 | - | - |
| DEINH7 | Inhibitory neurons, hypothalamus | 259 | 10 | 2.58E-06 | **0.0004** | 7.72 | 15.76 | - | - |
| HYPEP2 | Peptidergic neurons, hypothalamus | 465 | 13 | 2.83E-06 | **0.0004** | 4.84 | 16.74 | - | - |
| HBINH4 | Inhibitory neurons, hindbrain | 1639 | 25 | 5.18E-06 | **0.0007** | 3.82 | 6.90 | - | - |
| HBINH2 | Inhibitory neurons, hindbrain | 3492 | 40 | 5.23E-06 | **0.0008** | 4.67 | 6.62 | - | - |
| HBNOR | Noradrenergic neurons of the medulla | 1611 | 24 | 1.25E-05 | **0.0018** | 10.24 | 12.90 | - | - |
| HBADR | Adrenergic cell groups of the medulla | 2574 | 32 | 1.37E-05 | **0.0020** | 8.27 | 7.82 | 0.2581 | 1 |
| HYPEP6 | Orexin-producing neurons, hypothalamus | 1542 | 23 | 1.93E-05 | **0.0028** | 6.12 | 9.58 | - | - |
| TEINH1 | Inhibitory neurons, pallidum | 991 | 17 | 5.16E-05 | **0.0074** | 3.08 | 3.74 | - | - |
| MEGLU2 | Excitatory neurons, midbrain | 1097 | 18 | 5.39E-05 | **0.0078** | 3.31 | 4.43 | - | - |
| HBINH5 | Inhibitory neurons, hindbrain | 3288 | 36 | 5.43E-05 | **0.0078** | 3.35 | 3.74 | - | - |
| DEINH8 | Inhibitory neurons, hypothalamus | 308 | 9 | 7.48E-05 | **0.0108** | 7.18 | 15.65 | - | - |
| MBDOP2 | Dopaminergic neurons, ventral midbrain (SNc, VTA) | 1494 | 21 | 0.0001 | **0.0161** | 11.13 | 9.29 | 0.1175 | 1 |
| MEGLU10 | Excitatory neurons, midbrain | 792 | 14 | 0.0002 | **0.0267** | 2.83 | 4.69 | - | - |
| DEINH3 | Inhibitory neurons, hypothalamus | 1347 | 19 | 0.0002 | **0.0345** | 3.30 | 3.53 | - | - |
| HYPEP4 | Oxytocin-producing cells, hypothalamus | 923 | 15 | 0.0003 | **0.0386** | 8.67 | 9.35 | - | - |
| MEGLU14 | Glutamatergic projection neurons of the raphe nucleus | 1832 | 23 | 0.0003 | **0.0388** | 7.76 | 15.15 | - | - |
| MEGLU9 | Excitatory neurons, midbrain | 661 | 12 | 0.0004 | 0.0626 | 2.89 | 6.26 | - | - |
| MEINH14 | Inhibitory neurons, midbrain | 571 | 11 | 0.0005 | 0.0670 | 4.42 | 13.65 | - | - |
| HBINH3 | Inhibitory neurons, hindbrain | 872 | 14 | 0.0005 | 0.0709 | 5.31 | 9.29 | - | - |
| HBGLU6 | Excitatory neurons, hindbrain | 3518 | 35 | 0.0005 | 0.0719 | 4.15 | 4.83 | - | - |
| HBINH6 | Inhibitory neurons, hindbrain | 3125 | 31 | 0.0013 | 0.1811 | 4.47 | 5.65 | - | - |
| HBGLU4 | Excitatory neurons, hindbrain | 2578 | 27 | 0.0013 | 0.1855 | 6.87 | 8.38 | - | - |
| MEINH3 | Inhibitory neurons, midbrain | 1104 | 15 | 0.0017 | 0.2426 | 2.76 | 4.22 | - | - |
| SCGLU1 | Excitatory neurons, spinal cord | 583 | 10 | 0.0020 | 0.2909 | 4.04 | 13.41 | - | - |
| HYPEP5 | Vasopressin-producing cells, hypothalamus | 820 | 12 | 0.0027 | 0.3958 | 6.43 | 7.82 | - | - |
| DECHO1 | Cholinergic neurons, septal nucleus, Meissnert and diagonal band | 2442 | 25 | 0.0028 | 0.4013 | 6.23 | 6.16 | 0.5821 | 1 |
| HBCHO2 | Cholinergic neurons, hindbrain | 3436 | 32 | 0.0029 | 0.4216 | 4.19 | 6.44 | - | - |
| HBINH1 | Inhibitory neurons, hindbrain | 549 | 9 | 0.0045 | 0.6547 | 6.10 | 23.68 | - | - |
| HBCHO4 | Afferent nuclei of cranial nerves III-V | 2967 | 28 | 0.0048 | 0.6951 | 3.70 | 6.04 | - | - |
| HBGLU7 | Excitatory neurons, hindbrain | 3277 | 30 | 0.0054 | 0.7705 | 3.76 | 4.72 | - | - |
| SCGLU3 | Excitatory neurons, spinal cord | 782 | 11 | 0.0055 | 0.7973 | 3.78 | 10.65 | - | - |
| HBSER1 | Serotonergic neurons, hindbrain | 2200 | 22 | 0.0069 | 0.9916 | 11.02 | 6.79 | 0.0218 | 0.4578 |
| MEGLU1 | Excitatory neurons, midbrain | 1425 | 16 | 0.0077 | 1 | 2.65 | 4.86 | - | - |
| HYPEP7 | Pmch neurons, hypothalamus | 1088 | 13 | 0.0099 | 1 | 4.24 | 13.35 | - | - |
| HBGLU9 | Excitatory neurons, hindbrain | 2739 | 25 | 0.0121 | 1 | 5.13 | 5.17 | - | - |
| DEGLU4 | Excitatory neurons, thalamus | 1185 | 13 | 0.0191 | 1 | 3.17 | 5.53 | - | - |
| HBGLU10 | Excitatory neurons, hindbrain | 708 | 9 | 0.0215 | 1 | 3.91 | 11.03 | - | - |
| HBINH7 | Inhibitory neurons, hindbrain | 1881 | 18 | 0.0221 | 1 | 3.90 | 7.83 | - | - |
| MEINH4 | Inhibitory neurons, midbrain | 854 | 10 | 0.0260 | 1 | 5.14 | 5.34 | - | - |
| HBGLU8 | Excitatory neurons, hindbrain | 3561 | 29 | 0.0293 | 1 | 6.04 | 6.53 | - | - |
| MEGLU3 | Excitatory neurons, midbrain | 755 | 9 | 0.0308 | 1 | 2.62 | 6.77 | - | - |
| OBDOP1 | Dopaminergic periglomerular interneuron, olfactory bulb | 415 | 6 | 0.0334 | 1 | 17.15 | 8.20 | - | - |
| SCINH10 | Inhibitory neurons, spinal cord | 811 | 9 | 0.0452 | 1 | 3.61 | 10.54 | - | - |
| SCINH3 | Inhibitory neurons, spinal cord | 703 | 8 | 0.0510 | 1 | 14.95 | 10.00 | - | - |
| MEINH12 | Inhibitory neurons, midbrain | 708 | 8 | 0.0527 | 1 | 2.91 | 8.24 | - | - |
| TEINH5 | Interneuron-selective interneurons, cortex/hippocampus | 488 | 6 | 0.0638 | 1 | 2.23 | 5.77 | - | - |
| HBCHO1 | Cholinergic neurons, hindbrain | 2586 | 21 | 0.0649 | 1 | 4.12 | 10.35 | - | - |
| DEGLU5 | Excitatory neurons, midbrain | 619 | 7 | 0.0676 | 1 | 2.69 | 4.43 | - | - |
| SCINH5 | Inhibitory neurons, spinal cord | 1354 | 12 | 0.0911 | 1 | 6.46 | 9.99 | - | - |
| MEINH2 | Inhibitory neurons, midbrain | 1511 | 13 | 0.0959 | 1 | 2.75 | 3.62 | - | - |
| HYPEP8 | Peptidergic neurons, hypothalamus | 962 | 9 | 0.1046 | 1 | 7.50 | 24.02 | - | - |
| MEGLU11 | Excitatory neurons, midbrain | 846 | 8 | 0.1166 | 1 | 5.02 | 5.56 | - | - |
| TEINH10 | R-LM border Cck interneurons, cortex/hippocampus | 1141 | 10 | 0.1235 | 1 | 4.02 | 4.23 | - | - |
| SYNOR3 | Noradrenergic neurons, sympathetic | 2667 | 20 | 0.1310 | 1 | 9.48 | 7.66 | 0.1842 | 1 |
| ENT4 | Cholinergic enteric neurons | 3465 | 25 | 0.1341 | 1 | 13.28 | 11.93 | 0.3431 | 1 |
| DEGLU3 | Excitatory neurons, thalamus | 1619 | 13 | 0.1413 | 1 | 3.41 | 5.56 | - | - |
| ENT7 | Cholinergic enteric neurons, VGLUT2 | 3011 | 22 | 0.1416 | 1 | 6.73 | 10.49 | - | - |
| MEGLU4 | Excitatory neurons, midbrain | 758 | 7 | 0.1493 | 1 | 6.03 | 6.26 | - | - |
| HBGLU1 | Excitatory neurons, hindbrain | 3042 | 22 | 0.1528 | 1 | 4.74 | 6.15 | - | - |
| SCGLU5 | Excitatory neurons, spinal cord | 801 | 7 | 0.1814 | 1 | 4.79 | 9.01 | - | - |
| SCINH8 | Inhibitory neurons, spinal cord | 811 | 7 | 0.1892 | 1 | 7.91 | 13.50 | - | - |
| MEINH10 | Inhibitory neurons, midbrain | 1107 | 9 | 0.1902 | 1 | 3.22 | 6.76 | - | - |
| ENT8 | Cholinergic enteric neurons, VGLUT2 | 3029 | 21 | 0.2132 | 1 | 5.22 | 8.83 | - | - |
| CBINH2 | Granular layer interneurons, cerebellum | 696 | 6 | 0.2151 | 1 | 17.05 | 14.64 | - | - |
| SCINH11 | Central canal neurons, spinal cord | 859 | 7 | 0.2287 | 1 | 53.31 | 29.03 | - | - |
| SYNOR5 | Noradrenergic erector muscle neurons | 2259 | 16 | 0.2293 | 1 | 20.13 | 9.80 | 0.0425 | 0.8925 |
| TEINH8 | Interneuron-selective interneurons, hippocampus | 870 | 7 | 0.2382 | 1 | 3.30 | 7.01 | - | - |
| TEGLU21 | Excitatory neurons, hippocampus CA1 | 1680 | 12 | 0.2635 | 1 | 4.09 | 4.42 | - | - |
| SCGLU4 | Excitatory neurons, spinal cord | 750 | 6 | 0.2666 | 1 | 3.39 | 8.48 | - | - |
| TEGLU12 | Excitatory neurons, cerebral cortex | 913 | 7 | 0.2763 | 1 | 4.18 | 5.95 | - | - |
| TEGLU10 | Excitatory neurons, cerebral cortex | 1394 | 10 | 0.2855 | 1 | 2.86 | 3.53 | - | - |
| TEINH19 | Hippocamposeptal projection, cortex/hippocampus | 1087 | 8 | 0.2924 | 1 | 5.18 | 5.04 | - | - |
| TEGLU20 | Excitatory neurons, cerebral cortex | 1407 | 10 | 0.2953 | 1 | 7.44 | 5.82 | - | - |
| MEGLU6 | Excitatory neurons, midbrain | 1584 | 11 | 0.3077 | 1 | 2.98 | 4.83 | - | - |
| MSN1 | D1 medium spiny neurons, striatum | 1112 | 8 | 0.3139 | 1 | 4.56 | 5.13 | - | - |
| DECHO2 | Cholinergic neurons, habenula | 966 | 7 | 0.3253 | 1 | 12.45 | 15.47 | - | - |
| ENT2 | Nitrergic enteric neurons | 3612 | 23 | 0.3385 | 1 | 10.44 | 9.57 | 0.5114 | 1 |
| TEGLU22 | Excitatory neurons, amygdala | 983 | 7 | 0.3413 | 1 | 5.19 | 4.76 | - | - |
| SYNOR1 | Noradrenergic erector muscle neurons | 2637 | 17 | 0.3532 | 1 | 8.54 | 6.94 | 0.1662 | 1 |
| ENT5 | Cholinergic enteric neurons | 2475 | 16 | 0.3558 | 1 | 10.03 | 10.01 | 0.3988 | 1 |
| PSPEP5 | Peptidergic (PEP1.2), DRG | 2145 | 14 | 0.3568 | 1 | 6.03 | 11.34 | - | - |
| TEINH12 | Non-border Cck interneurons, cortex/hippocampus | 1494 | 10 | 0.3626 | 1 | 2.89 | 4.38 | - | - |
| TEGLU8 | Excitatory neurons, cerebral cortex | 1172 | 8 | 0.3666 | 1 | 4.56 | 4.44 | - | - |
| TEGLU13 | Excitatory neurons, cerebral cortex | 1693 | 11 | 0.3889 | 1 | 3.54 | 4.75 | - | - |
| ENT3 | Nitrergic enteric neurons | 3382 | 21 | 0.3965 | 1 | 9.61 | 10.07 | - | - |
| TEGLU16 | Excitatory neurons, cerebral cortex | 1888 | 12 | 0.4098 | 1 | 6.37 | 5.30 | - | - |
| TEGLU23 | Excitatory neurons, hippocampus CA3 | 1261 | 8 | 0.4460 | 1 | 4.67 | 4.48 | - | - |
| MBCHO1 | Cholinergic neurons, midbrain red nucleus | 1624 | 10 | 0.4661 | 1 | 4.27 | 9.49 | - | - |
| SCINH6 | Inhibitory neurons, spinal cord | 1284 | 8 | 0.4665 | 1 | 5.74 | 10.16 | - | - |
| HBINH8 | Inhibitory neurons, hindbrain | 1831 | 11 | 0.4938 | 1 | 5.74 | 10.45 | - | - |
| TEGLU14 | Excitatory neurons, cerebral cortex | 2184 | 13 | 0.4999 | 1 | 2.90 | 4.91 | - | - |
| SYCHO2 | Cholinergic neurons, sympathetic | 2734 | 16 | 0.5207 | 1 | 6.36 | 7.41 | - | - |
| ENT9 | Cholinergic enteric neurons | 3653 | 21 | 0.5505 | 1 | 9.83 | 12.43 | - | - |
| TEGLU11 | Excitatory neurons, cerebral cortex | 1038 | 6 | 0.5596 | 1 | 2.71 | 3.92 | - | - |
| TEINH9 | Non-border Cck interneurons, hippocampus | 1038 | 6 | 0.5596 | 1 | 4.93 | 5.23 | - | - |
| SYNOR2 | Noradrenergic neurons, sympathetic | 2625 | 15 | 0.5619 | 1 | 9.71 | 7.59 | 0.1255 | 1 |
| SCINH1 | Inhibitory neurons, spinal cord | 1220 | 7 | 0.5640 | 1 | 4.60 | 11.37 | - | - |
| SYNOR4 | Noradrenergic erector muscle neurons | 2659 | 15 | 0.5832 | 1 | 9.83 | 6.98 | 0.114 | 1 |
| TECHO | Cholinergic interneurons, telencephalon | 1793 | 10 | 0.5959 | 1 | 2.40 | 6.80 | - | - |
| PSNF1 | Neurofilament (NF1), DRG | 2762 | 15 | 0.6453 | 1 | 6.88 | 8.27 | - | - |
| TEGLU2 | Excitatory neurons, cerebral cortex | 1158 | 6 | 0.6670 | 1 | 2.72 | 4.98 | - | - |
| PSNP4 | Non-peptidergic (NP2.1), DRG | 2820 | 15 | 0.6782 | 1 | 5.98 | 11.60 | - | - |
| TEGLU15 | Excitatory neurons, cerebral cortex | 1556 | 8 | 0.6867 | 1 | 4.18 | 4.33 | - | - |
| TEINH11 | R-LM border Cck interneurons, cortex/hippocampus | 1436 | 7 | 0.7334 | 1 | 4.96 | 4.30 | - | - |
| TEGLU7 | Excitatory neurons, cerebral cortex | 1442 | 7 | 0.7374 | 1 | 3.45 | 3.93 | - | - |
| ENT1 | Nitrergic enteric neurons | 2447 | 12 | 0.7713 | 1 | 4.84 | 10.91 | - | - |
| PSPEP6 | Peptidergic (TrpM8), DRG | 1777 | 8 | 0.8179 | 1 | 5.98 | 8.68 | - | - |
| ENT6 | Cholinergic enteric neurons | 2805 | 13 | 0.8431 | 1 | 5.48 | 9.36 | - | - |
| DEGLU2 | Excitatory neurons, hypothalamus | 1645 | 7 | 0.8490 | 1 | 3.26 | 4.75 | - | - |
| TEGLU4 | Excitatory neurons, cerebral cortex | 1471 | 6 | 0.8623 | 1 | 2.43 | 3.71 | - | - |
| PSPEP4 | Peptidergic (PEP1.1), DRG | 2683 | 12 | 0.8676 | 1 | 7.68 | 8.55 | - | - |
| HBGLU5 | Excitatory neurons, hindbrain | 2160 | 9 | 0.8902 | 1 | 4.60 | 5.96 | - | - |
| PSNP2 | Non-peptidergic (NP1.1), DRG | 3065 | 13 | 0.9182 | 1 | 15.55 | 8.94 | - | - |
| PSNP5 | Non-peptidergic (NP2.2), DRG | 3077 | 13 | 0.9208 | 1 | 6.51 | 10.15 | - | - |
| PSPEP7 | Peptidergic (TrpM8), DRG | 1656 | 6 | 0.9253 | 1 | 11.50 | 9.76 | - | - |
| SYCHO1 | Cholinergic neurons, sympathetic | 2721 | 11 | 0.9299 | 1 | 7.98 | 7.83 | - | - |
| PSPEP1 | Peptidergic (PEP2), DRG | 2924 | 12 | 0.9302 | 1 | 8.84 | 10.20 | - | - |
| PSPEP3 | Peptidergic (PEP1..4), DRG | 3131 | 13 | 0.9316 | 1 | 9.33 | 8.77 | - | - |
| PSNP1 | Non-peptidergic (Th), DRG | 3137 | 13 | 0.9328 | 1 | 7.34 | 9.26 | - | - |
| PSPEP8 | Peptidergic (TrpM8), DRG | 2537 | 10 | 0.9339 | 1 | 11.47 | 8.74 | - | - |
| PSPEP2 | Peptidergic (PEP1.3), DRG | 3163 | 13 | 0.9375 | 1 | 8.27 | 8.72 | - | - |
| PSNF3 | Neurofilament (NF2/3), DRG | 2767 | 11 | 0.9388 | 1 | 7.41 | 8.48 | - | - |
| PSNP3 | Non-peptidergic (NP1.2), DRG | 3066 | 12 | 0.9539 | 1 | 29.76 | 9.34 | - | - |
| SZNBL | Neuronal intermediate progenitor cells | 1802 | 6 | 0.9557 | 1 | 4.82 | 23.22 | - | - |
| PSNF2 | Neurofilament (NF4/5), DRG | 2688 | 10 | 0.9587 | 1 | 14.99 | 9.17 | - | - |
| PSNP6 | Non-peptidergic (NP3), DRG | 3524 | 14 | 0.9606 | 1 | 11.26 | 16.83 | - | - |
| DEGLU1 | Excitatory neurons, thalamus | 2056 | 6 | 0.9834 | 1 | 3.94 | 5.08 | - | - |
| DEINH2 | Inhibitory neurons, thalamus | 991 | 5 | - | - | 4.61 | 5.04 | - | - |
| DGNBL2 | Granule neuroblasts, dentate gyrus | 715 | 5 | - | - | 3.46 | 5.09 | - | - |
| MEGLU8 | Excitatory neurons, midbrain | 531 | 5 | - | - | 2.62 | 5.32 | - | - |
| MEINH7 | Inhibitory neurons, midbrain | 592 | 5 | - | - | 3.24 | 9.11 | - | - |
| MEINH8 | Inhibitory neurons, midbrain | 332 | 5 | - | - | 4.15 | 24.00 | - | - |
| MSN2 | D2 medium spiny neurons, striatum | 955 | 5 | - | - | 4.87 | 5.12 | - | - |
| MSN3 | D2 medium spiny neurons, striatum | 615 | 5 | - | - | 5.10 | 4.57 | - | - |
| MSN4 | D1 medium spiny neurons, striatum | 710 | 5 | - | - | 6.32 | 4.60 | - | - |
| MSN5 | Patch D1/D2 neurons, striatum | 443 | 5 | - | - | 3.51 | 7.23 | - | - |
| OBNBL1 | Neuroblasts, olfactory | 1055 | 5 | - | - | 8.07 | 11.63 | - | - |
| SCGLU2 | Excitatory neurons, spinal cord | 572 | 5 | - | - | 2.88 | 10.67 | - | - |
| SCINH7 | Inhibitory neurons, spinal cord | 701 | 5 | - | - | 10.21 | 13.98 | - | - |
| SCINH9 | Inhibitory neurons, spinal cord | 583 | 5 | - | - | 12.71 | 13.05 | - | - |
| TEGLU5 | Excitatory neurons, cerebral cortex | 1402 | 5 | - | - | 2.61 | 5.87 | - | - |
| TEINH13 | Trilaminar cells, hippocampus | 1150 | 5 | - | - | 2.79 | 4.55 | - | - |
| TEINH7 | Interneuron-selective interneurons, hippocampus | 1053 | 5 | - | - | 5.11 | 5.67 | - | - |
| DEINH1 | Inhibitory neurons, thalamus | 813 | 4 | - | - | 4.03 | 5.04 | - | - |
| DGGRC1 | Granule neuroblasts, dentate gyrus | 522 | 4 | - | - | 4.70 | 4.43 | - | - |
| DGNBL1 | Granule neuroblasts, dentate gyrus | 779 | 4 | - | - | 4.04 | 7.68 | - | - |
| MEGLU5 | Excitatory neurons, midbrain | 843 | 4 | - | - | 2.41 | 5.85 | - | - |
| MEGLU7 | Excitatory neurons, midbrain | 646 | 4 | - | - | 2.76 | 6.68 | - | - |
| MEINH5 | Inhibitory neurons, midbrain | 785 | 4 | - | - | 9.36 | 8.55 | - | - |
| MEINH6 | Inhibitory neurons, midbrain | 1128 | 4 | - | - | 3.52 | 6.91 | - | - |
| OBINH2 | Inhibitory neurons, olfactory bulb | 486 | 4 | - | - | 3.80 | 5.61 | - | - |
| OBNBL3 | Neuroblasts, olfactory bulb | 479 | 4 | - | - | 4.01 | 5.36 | - | - |
| SCGLU6 | Excitatory neurons, spinal cord | 609 | 4 | - | - | 3.94 | 12.88 | - | - |
| TEGLU19 | Excitatory neurons, cerebral cortex | 1191 | 4 | - | - | 3.29 | 5.22 | - | - |
| TEGLU9 | Excitatory neurons, cerebral cortex | 973 | 4 | - | - | 2.94 | 4.69 | - | - |
| TEINH15 | CGE-derived neuroglia form cells, cortex/hippocampus | 694 | 4 | - | - | 6.10 | 5.10 | - | - |
| TEINH16 | Ivy and MGE-derived neuroglia form cells, cortex/hippocampus | 788 | 4 | - | - | 2.49 | 4.90 | - | - |
| TEINH18 | Basket and bistratified cells, cortex/hippocampus | 1103 | 4 | - | - | 4.82 | 4.71 | - | - |
| TEINH21 | Sleep-active, long-range projection interneurons, cortex/hippocampus | 612 | 4 | - | - | 2.93 | 6.07 | - | - |
| TEINH6 | Interneuron-selective interneurons, cortex/hippocampus | 511 | 4 | - | - | 2.84 | 7.24 | - | - |
| CBNBL1 | Neuroblasts, cerebellum | 365 | 3 | - | - | 8.27 | 15.34 | - | - |
| CR | Cajal-Retzius cells, hippocampus | 395 | 3 | - | - | 20.70 | 17.25 | - | - |
| DETPH | Neuroblast-like, habenula | 877 | 3 | - | - | 21.54 | 106.51 | - | - |
| DGGRC2 | Granule neurons, dentate gyrus | 659 | 3 | - | - | 4.91 | 5.32 | - | - |
| MSN6 | Matrix D1 neurons, striatum | 181 | 3 | - | - | 4.35 | 7.06 | - | - |
| OBDOP2 | Inhibitory neurons, olfactory bulb | 263 | 3 | - | - | 5.48 | 7.32 | - | - |
| SCGLU10 | Excitatory neurons, spinal cord | 438 | 3 | - | - | 4.02 | 13.85 | - | - |
| SEPNBL | Neuroblasts, septum | 383 | 3 | - | - | 7.86 | 13.30 | - | - |
| TEGLU1 | Excitatory neurons, cerebral cortex | 586 | 3 | - | - | 5.57 | 5.88 | - | - |
| TEGLU18 | Excitatory neurons, cerebral cortex | 1067 | 3 | - | - | 27.77 | 4.84 | - | - |
| TEGLU24 | Excitatory neurons, hippocampus CA1 | 1053 | 3 | - | - | 7.01 | 4.72 | - | - |
| TEGLU3 | Excitatory neurons, cerebral cortex | 783 | 3 | - | - | 4.38 | 4.61 | - | - |
| TEGLU6 | Excitatory neurons, cerebral cortex | 464 | 3 | - | - | 2.84 | 5.12 | - | - |
| TEINH20 | Inhibitory interneurons, hippocampus | 1074 | 3 | - | - | 2.33 | 5.28 | - | - |
| MEINH11 | Inhibitory neurons, midbrain | 503 | 2 | - | - | 3.68 | 8.34 | - | - |
| OBINH3 | Inhibitory neurons, olfactory bulb | 314 | 2 | - | - | 5.82 | 5.53 | - | - |
| OBINH5 | External plexiform layer interneuron, olfactory bulb | 517 | 2 | - | - | 47.62 | 13.20 | - | - |
| OBNBL2 | Neuroblasts, olfactory bulb | 724 | 2 | - | - | 9.68 | 12.57 | - | - |
| SCGLU7 | Excitatory neurons, spinal cord | 337 | 2 | - | - | 7.15 | 23.84 | - | - |
| SCGLU8 | Excitatory neurons, spinal cord | 577 | 2 | - | - | 6.01 | 20.64 | - | - |
| SCGLU9 | Excitatory neurons, spinal cord | 483 | 2 | - | - | 18.86 | 12.43 | - | - |
| SCINH2 | Inhibitory neurons, spinal cord | 645 | 2 | - | - | 3.03 | 8.29 | - | - |
| SCINH4 | Inhibitory neurons, spinal cord | 526 | 2 | - | - | 2.15 | 12.87 | - | - |
| TEGLU17 | Excitatory neurons, cerebral cortex | 1043 | 2 | - | - | 6.49 | 3.75 | - | - |
| TEINH17 | Axo-axonic, cortex/hippocampus | 760 | 2 | - | - | 4.47 | 4.85 | - | - |
| TEINH4 | Interneuron-selective interneurons, cortex/hippocampus | 432 | 2 | - | - | 2.79 | 5.82 | - | - |
| CBNBL2 | Neuroblasts, cerebellum | 148 | 1 | - | - | 6.52 | 12.86 | - | - |
| CBPC | Purkinje cells | 511 | 1 | - | - | 2.55 | 19.71 | - | - |
| MEINH9 | Inhibitory neurons, midbrain | 689 | 1 | - | - | 2.69 | 6.51 | - | - |
| OBINH1 | Inner horizontal cell, olfactory bulb | 242 | 1 | - | - | 5.02 | 5.67 | - | - |
| OBNBL4 | Inhibitory neurons, olfactory bulb | 252 | 1 | - | - | 26.70 | 14.62 | - | - |
| OBNBL5 | Inhibitory neurons, olfactory bulb | 313 | 1 | - | - | 3.39 | 8.16 | - | - |
| TEINH14 | CGE-derived neuroglia form cells Cxcl14+, cortex/hippocampus | 462 | 1 | - | - | 4.67 | 5.33 | - | - |
| CBGRC | Granule neurons, cerebellum | 186 | 0 | - | - | 0.00 | 17.88 | - | - |
| CBINH1 | Molecular layer interneurons, cerebellum | 413 | 0 | - | - | 0.00 | 10.46 | - | - |
| MEINH1 | Inhibitory neurons, midbrain | 199 | 0 | - | - | 0.00 | 19.00 | - | - |
| OBINH4 | Inhibitory neurons, olfactory bulb | 204 | 0 | - | - | 0.00 | 13.29 | - | - |

**Supplemental Table S9 (A & B)**

| Supplemental Table S9A. Whole Hypothalamus Level Enrichment Analysis for all imprinted genes in cell lineage populations in the Chen, Wu (5) dataset. *Identity* – neural lineage identities for the cells used in this analysis. *Up Reg* – number of upregulated genes with *q* ≤ 0.05 and Log2FC > 0 (total number of genes in the dataset in brackets); *IG* – number of imprinted genes upregulated with *q* ≤ 0.05 and Log2FC > 0 (total number of IGs in the dataset in brackets); *ORA p* – *p* value from over representation analysis on groups with minimum 5% of total IGs; *ORA q* – Bonferroni corrected *p* value from ORA; *Mean FC IG* – mean fold change for upregulated imprinted genes; *Mean FC Rest* – mean fold change for all other upregulated genes; *GSEA p* – *p* value from Gene Set Enrichment Analysis for identity groups with 15+ IGs and Mean FC IG > Mean FC Rest; *GSEA q* – Bonferroni corrected *p* values from GSEA. | | | | | | | |
| --- | --- | --- | --- | --- | --- | --- | --- |
| Identity | **Up Reg**  **(14,687)** | **IG**  **(91)** | **ORA *p*** | **ORA *q*** | **Mean FC**  **IG** | **Mean FC**  **Rest** | **GSEA**  ***p*** |
| Neuron | 5787 | 54 | 9.06E-05 | **0.0010** | 4.81 | 3.45 | **0.0221** |
| Oligodendrocyte Progenitor Cell | 1602 | 13 | 0.1893 | 1 | 2.82 | 3.02 | - |
| Myelinating Oligodendrocyte | 1830 | 14 | 0.2391 | 1 | 2.65 | 3.9 | - |
| Immature Oligodendrocyte | 1577 | 11 | 0.3864 | 1 | 2.49 | 3.15 | - |
| Astrocyte | 2975 | 19 | 0.4827 | 1 | 1.87 | 3.75 | - |
| Epithelial cell 2 | 1056 | 7 | 0.4836 | 1 | 6.98 | 7.34 | - |
| Epithelial cell 1 | 2249 | 14 | 0.5373 | 1 | 3.96 | 6.67 | - |
| SCO | 1288 | 7 | 0.6962 | 1 | 6.48 | 8.64 | - |
| Macrophage | 1834 | 10 | 0.7144 | 1 | 5.3 | 10.96 | - |
| Tanycyte | 2938 | 12 | 0.9665 | 1 | 4.49 | 3.55 | - |
| Ependymocyte | 3621 | 15 | 0.9776 | 1 | 3.41 | 11.19 | - |
| Microglia | 625 | 3 | - | - | 5.24 | 10.72 | - |
| Proliferating Oligodendrocyte Progenitor Cell | 1264 | 1 | - | - | 1.65 | 17.41 | - |

| Supplemental Table S9B. Whole Hypothalamus Level Enrichment Analysis for all imprinted genes in neuronal subpopulations in the Chen, Wu (5) dataset. *Identity* – neural subpopulation identities for the cells used in this analysis. *Marker Genes* – Genes identified by Chen, Wu (5) with distinct expression in these neuronal subtypes for the purposes of identification. All other column descriptions can be found in the legend of Supplemental Table S9A. | | | | | | | |
| --- | --- | --- | --- | --- | --- | --- | --- |
| Identity | **Marker**  **Genes** | **Up Reg**  **(12,238)** | **IG**  **(80)** | **ORA *p*** | **ORA *q*** | **Mean FC**  **IG** | **Mean FC**  **Rest** |
| GABA17 | *Slc6a3* | 825 | 18 | 4.57E-06 | **1.19E-04** | 2.73 | 3.30 |
| GABA8 | *Vipr2* | 480 | 11 | 0.0003 | **0.0071** | 1.96 | 2.73 |
| GABA13 | *Slc18a2, Gal* | 569 | 12 | 0.0003 | **0.0079** | 1.89 | 2.02 |
| GABA15 | *Agrp* | 766 | 13 | 0.0013 | **0.0339** | 2.40 | 2.79 |
| GABA16 | *Cox6a2* | 258 | 6 | 0.0068 | 0.1762 | 1.68 | 2.70 |
| GABA14 | *Cbln4* | 271 | 6 | 0.0085 | 0.2220 | 1.68 | 2.54 |
| GABA2 | *Npas1* | 278 | 6 | 0.0096 | 0.2499 | 2.29 | 4.22 |
| GABA18 | *Lhx1, Klhl1* | 200 | 5 | 0.0099 | 0.2576 | 1.55 | 3.06 |
| GABA9 | *Vip* | 219 | 5 | 0.0142 | 0.3703 | 1.39 | 3.77 |
| GABA10 | *Prok2* | 605 | 9 | 0.0169 | 0.4391 | 3.03 | 4.28 |
| Glu9 | *Gng8, Samd3* | 345 | 6 | 0.0252 | 0.6556 | 1.34 | 3.66 |
| Glu1 | *Crh* | 182 | 4 | 0.0312 | 0.8120 | 1.68 | 4.38 |
| GABA11 | *Ghrh* | 392 | 6 | 0.0430 | 1 | 6.50 | 2.91 |
| GABA5 | *Lhx8* | 569 | 7 | 0.0780 | 1 | 1.28 | 3.09 |
| Glu14 | *Avp, Sim1* | 945 | 10 | 0.0875 | 1 | 2.07 | 3.30 |
| Glu15 | *Sst, Prdm8* | 650 | 7 | 0.1319 | 1 | 1.69 | 3.79 |
| Glu12 | *Vgll2* | 339 | 4 | 0.1811 | 1 | 1.70 | 3.88 |
| Glu13 | *Pomc* | 471 | 5 | 0.1945 | 1 | 3.36 | 4.04 |
| GABA12 | *Crabp1* | 354 | 4 | 0.2010 | 1 | 1.63 | 2.61 |
| Glu6 | *Tac1* | 607 | 6 | 0.2052 | 1 | 2.39 | 3.58 |
| Glu7 | *Fezf1, Lbhd2* | 609 | 6 | 0.2072 | 1 | 1.44 | 2.13 |
| Glu11 | *Kiss1* | 468 | 4 | 0.3663 | 1 | 7.78 | 3.68 |
| GABA3 | *Bcl11b* | 484 | 4 | 0.3902 | 1 | 1.41 | 3.36 |
| Glu8 | *Lbhd2, Cartpt* | 534 | 4 | 0.4641 | 1 | 3.34 | 2.97 |
| Glu4 | *Shox2* | 1690 | 8 | 0.8792 | 1 | 2.25 | 3.09 |
| Glu5 | *Foxb1* | 984 | 4 | 0.8942 | 1 | 2.20 | 3.69 |
| GABA6 | *Pax6* | 228 | 3 | - | - | 1.03 | 3.86 |
| GABA7 | *Trh* | 194 | 3 | - | - | 3.87 | 3.46 |
| Glu3 | *Fezf2, Samd3* | 203 | 3 | - | - | 1.07 | 5.35 |
| GABA1 | *Pvalb* | 738 | 2 | - | - | 3.81 | 4.57 |
| GABA4 | *Gm13498* | 254 | 2 | - | - | 3.97 | 4.43 |
| Glu10 | *Trh* | 228 | 2 | - | - | 1.17 | 4.88 |
| Hista | *Hdc* | 585 | 2 | - | - | 3.98 | 5.14 |
| Glu2 | *Sln* | 376 | 0 | - | - | 0.00 | 4.38 |

**Supplemental Table S10 (A & B)**

| Supplemental Table S10A. Whole Hypothalamus Level Enrichment Analysis for all imprinted genes in cell lineage populations in the Romanov, Zeisel (6) dataset. *Identity* – neural lineage identities for the cells used in this analysis. All other column descriptions can be found in the legend of Supplemental Table S9A. | | | | | | | |
| --- | --- | --- | --- | --- | --- | --- | --- |
| Identity | **Up Reg**  **(14,914)** | **IG**  **(94)** | **ORA *p*** | **ORA *q*** | **Mean FC**  **IG** | **Mean FC**  **Rest** | **GSEA**  ***p*** |
| neurons | 8806 | 68 | 0.0050 | **0.0202** | 2.53 | 2.07 | **0.0105** |
| endothelial | 2077 | 14 | 0.4377 | 1 | 9.84 | 7.62 | - |
| ependymal | 2232 | 13 | 0.6656 | 1 | 4.13 | 5.91 | - |
| oligos | 3047 | 16 | 0.8288 | 1 | 2.05 | 2.16 | - |
| astrocytes | 769 | 4 | - | - | 1.95 | 4.67 | - |
| vascular smooth muscle | 632 | 4 | - | - | 8.18 | 7.85 | - |
| microglia | 692 | 1 | - | - | 9.84 | 27.66 | - |

| Supplemental Table S10B. Whole Hypothalamus Level Enrichment Analysis for all imprinted genes in neuronal subpopulations in the Romanov, Zeisel (6) dataset. *Identity* – neural subpopulation identities for the cells used in this analysis. *Marker Genes* – Genes identified by Romanov, Zeisel (6) with distinct expression in these neuronal subtypes for the purposes of identification. All other column descriptions can be found in the legend of Supplemental Table S9A. | | | | | | | |
| --- | --- | --- | --- | --- | --- | --- | --- |
| Identity | **Marker**  **Genes** | **Up Reg**  **(12,243)** | **IG**  **(86)** | **ORA *p*** | **ORA *q*** | **Mean FC**  **IG** | **Mean FC**  **Rest** |
| GABA 5 | *Calcr, Lhx1* | 40 | 6 | 2.47E-07 | **1.73E-06** | 3.47 | 4.95 |
| GABA 14 | *Npy, Agrp* | 193 | 6 | 0.0019 | **0.0132** | 2.94 | 6.12 |
| *Ghrh* | *Ghrh* | 234 | 6 | 0.0049 | **0.0342** | 18.31 | 17.22 |
| Dopamine 4 | *Th, Slc6a3, Slc18a2* | 272 | 6 | 0.0100 | 0.0697 | 5.62 | 5.64 |
| *Avp* 2, high | *Avp* | 217 | 5 | 0.0154 | 0.1078 | 3.74 | 5.26 |
| GABA 11 | *Nts* | 319 | 5 | 0.0636 | 0.4450 | 2.25 | 2.82 |
| *Vglut2* 11 | *Slc17a6* | 1840 | 12 | 0.5924 | 1 | 3.25 | 2.97 |
| GABA 12 | *Nts* | 162 | 4 | - | - | 1.78 | 3.08 |
| *Qrfp* | *Qrfp* | 257 | 4 | - | - | 10.77 | 7.58 |
| Dopamine 1 | *Th, Slc18a2* | 217 | 3 | - | - | 5.46 | 4.26 |
| Dopamine 2 (low *Slc18a2*) | *Th* | 135 | 3 | - | - | 6.97 | 6.47 |
| Dopamine 3 | *Th, Slc18a2* | 335 | 3 | - | - | 7.79 | 4.26 |
| GABA 2 | *Gucy1a3* | 198 | 3 | - | - | 2.34 | 3.70 |
| *Hcrt* | *Hcrt* | 129 | 3 | - | - | 5.46 | 6.30 |
| *Hmit*+/- | *Slc2a13* | 16 | 3 | - | - | 1.88 | 5.24 |
| *Vglut2* 15 | *Hcn1,*  *6430411K18Rik* | 213 | 3 | - | - | 8.84 | 8.05 |
| circadian 2 | *Nms, Vip+/-* | 53 | 2 | - | - | 2.53 | 8.31 |
| GABA 4 | *Crh, Pgr15l* | 21 | 2 | - | - | 1.35 | 3.55 |
| Oxytocin 1 | *Oxt* | 331 | 2 | - | - | 3.02 | 4.10 |
| Oxytocin 4 | *Oxt* | 245 | 2 | - | - | 2.93 | 4.81 |
| *Pmch* | *Pmch* | 239 | 2 | - | - | 2.51 | 6.04 |
| *Trh* 3 (high) | *Trh, Cartpt* | 178 | 2 | - | - | 3.83 | 6.38 |
| *Vglut2* 1 | *Penk* | 294 | 2 | - | - | 9.13 | 15.24 |
| *Vglut2* 10 | *Morn4,Prrc2a* | 252 | 2 | - | - | 9.92 | 8.29 |
| *Vglut2* 17 | *A930013F10Rik,*  *Pou2f2* | 122 | 2 | - | - | 11.42 | 10.93 |
| *Adcyap1* 2 | *Adcyap1* | 236 | 1 | - | - | 1.81 | 3.21 |
| *Avp* 1, high | *Avp* | 141 | 1 | - | - | 3.10 | 5.71 |
| *Avp* 3, medium | *Avp* | 105 | 1 | - | - | 3.49 | 6.29 |
| circadian 3 | *Per2* | 19 | 1 | - | - | 9.08 | 6.08 |
| GABA 10 | *-* | 81 | 1 | - | - | 1.17 | 3.45 |
| GABA 13 | *Gal* | 126 | 1 | - | - | 4.40 | 3.61 |
| GABA 3 | *Crh+/-, Lhx6* | 144 | 1 | - | - | 1.66 | 3.66 |
| GABA 9 | *-* | 141 | 1 | - | - | 1.72 | 3.23 |
| GABA1 | *-* | 135 | 1 | - | - | 6.25 | 4.07 |
| *Sst* 1, low | *Sst* | 63 | 1 | - | - | 7.74 | 4.49 |
| *Sst* 3, medium | *Sst* | 202 | 1 | - | - | 6.36 | 5.05 |
| *Vglut2* 12 | *Mgat4b* | 199 | 1 | - | - | 7.09 | 6.34 |
| *Vglut2* 13 | *Ninl, Rfx5, Zfp346* | 149 | 1 | - | - | 6.90 | 6.59 |
| *Vglut2* 16 | *Zfp975, Tnr* | 371 | 1 | - | - | 9.30 | 11.02 |
| *Vglut2* 18 | *Zfp458, Ppp1r12b,* | 191 | 1 | - | - | 27.13 | 22.64 |
| *Vglut2* 2 | *Crh*+/- | 117 | 1 | - | - | 1.89 | 4.32 |
| *Vglut2* 5 | *Myt1, Lhx9* | 109 | 1 | - | - | 7.33 | 12.30 |
| *Vglut2* 6 | *Prmt8, Ugdh* | 46 | 1 | - | - | 5.98 | 5.29 |
| *Vglut2* 7 | Pgam, Snx12 | 93 | 1 | - | - | 7.17 | 10.26 |
| *Vglut2* 8 | *-* | 47 | 1 | - | - | 2.49 | 5.01 |
| *Adcyap1* 1 | *Adcyap1, Tac1* | 183 | 0 | - | - | 0.00 | 3.91 |
| circadian 1 | *Vip, Grp*+/- | 30 | 0 | - | - | 0.00 | 12.44 |
| GABA 15 | *Npy* | 67 | 0 | - | - | 0.00 | 10.03 |
| GABA 6 (Otof, Lhx1) | *Otof, Lhx1* | 33 | 0 | - | - | 0.00 | 3.54 |
| GABA 7 | *Pomc+/-* | 16 | 0 | - | - | 0.00 | 5.59 |
| GABA 8 | *-* | 74 | 0 | - | - | 0.00 | 4.10 |
| Gad-low, *Gnrh*-/+ | *Gnrh-/+* | 44 | 0 | - | - | 0.00 | 9.53 |
| *Npvf* | *Npvf* | 147 | 0 | - | - | 0.00 | 6.95 |
| Oxytocin 2 | *Oxt* | 254 | 0 | - | - | 0.00 | 5.14 |
| Oxytocin 3 | *Oxt* | 166 | 0 | - | - | 0.00 | 5.37 |
| *Sst* 2, high | *Sst* | 215 | 0 | - | - | 0.00 | 5.16 |
| *Trh* 1 (low) | *Trh* | 31 | 0 | - | - | 0.00 | 5.35 |
| *Trh* 2 (medium) | *Trh* | 48 | 0 | - | - | 0.00 | 4.80 |
| *Vglut2* 14 | *Col9a2* | 101 | 0 | - | - | 0.00 | 8.21 |
| *Vglut2* 3 | *Crh-+/-, low* | 1 | 0 | - | - | 0.00 | 5.19 |
| *Vglut2* 4 | *-* | 218 | 0 | - | - | 0.00 | 7.70 |
| *Vglut2* 9 | *Gpr149* | 61 | 0 | - | - | 0.00 | 8.77 |

**Supplemental Table S11**

| Supplemental Table S11. Comparison of Upregulated Genes for GABA5 in Romanov, Zeisel (6) and GABA17 in Chen, Wu (5) . 21 genes were upregulated in both subgroups which is over 50% of the upregulated genes found in GABA5 including the top 5 marker genes. Position of genes organised by *p* value is included for both subtypes alongside the *p* values, Bonferroni corrected *q* values, and the fold change (FC) values for expression in this neuronal subpopulation vs. background. *Prlr* did not have a significant q-value in GABA5 but is included due to its significance to TIDA neurons | | | | | | | | |
| --- | --- | --- | --- | --- | --- | --- | --- | --- |
| Gene | **Position in GABA5 sorted by *p* (38 upreg genes)** | **GABA5**  ***p*** | **GABA5**  ***q*** | **GABA5**  **FC** | **Position in GABA17 sorted by *p* (825 upreg genes)** | **GABA17 *p*** | **GABA17 *q*** | **GABA17 FC** |
| *Calcr* | 1 | 2.09E-38 | 1.30E-36 | 8.13 | 41 | 4.12E-13 | 4.67E-12 | 4.96 |
| *Lhx1* | 2 | 1.82E-10 | 1.13E-08 | 9.56 | 29 | 1.35E-16 | 1.53E-15 | 4.74 |
| *Slc6a3* | 3 | 6.44E-09 | 1.33E-07 | 15.71 | 1 | 2.20E-111 | 7.47E-110 | 69.29 |
| *Asb4* | 4 | 6.76E-08 | 4.19E-06 | 2.82 | 445 | 0.000408 | 0.002311 | 1.99 |
| *Lhx1os* | 5 | 1.13E-07 | 3.50E-06 | 4.74 | 23 | 1.25E-19 | 1.42E-18 | 4.61 |
| *Npbwr1* | 7 | 8.22E-06 | 0.000510 | 9.02 | 819 | 0.008319 | 0.047138 | 3.55 |
| *Diap3* | 9 | 4.79E-05 | 0.002967 | 7.51 | 12 | 1.73E-27 | 5.87E-26 | 9.97 |
| *Gpr83* | 11 | 0.000118 | 0.003456 | 4.27 | 14 | 3.49E-25 | 1.19E-23 | 6.27 |
| *Slc18a2* | 13 | 0.000158 | 0.001961 | 3.24 | 3 | 9.52E-44 | 1.62E-42 | 9.43 |
| *Peg10* | 16 | 0.000240 | 0.004955 | 2.25 | 40 | 9.95E-14 | 3.38E-12 | 2.46 |
| *Dlk1* | 20 | 0.000324 | 0.009853 | 1.89 | 124 | 1.24E-07 | 1.05E-06 | 1.93 |
| *Fam159b* | 21 | 0.000371 | 0.007659 | 4.98 | 72 | 5.46E-10 | 6.19E-09 | 4.19 |
| *Rab3b* | 24 | 0.000462 | 0.009550 | 1.85 | 30 | 1.39E-16 | 4.72E-15 | 1.96 |
| *Six6* | 25 | 0.000526 | 0.008157 | 4.35 | 11 | 1.01E-27 | 3.45E-26 | 6.41 |
| *AA388235* | 30 | 0.000863 | 0.036304 | 3.79 | 83 | 2.36E-09 | 8.01E-08 | 10.61 |
| *Hnrnpul1* | 33 | 0.001399 | 0.043371 | 3.87 | 602 | 0.001031 | 0.035041 | 2.20 |
| *Arhgap36* | 36 | 0.001821 | 0.028222 | 2.19 | 46 | 2.61E-12 | 4.44E-11 | 3.50 |
| *Th* | 37 | 0.002050 | 0.014126 | 2.24 | 8 | 1.45E-35 | 2.46E-34 | 8.29 |
| *Dlx2* | 38 | 0.002801 | 0.034730 | 4.09 | 139 | 4.42E-07 | 3.01E-06 | 3.23 |
| *Dlx6* | 39 | 0.003047 | 0.047231 | 9.07 | 193 | 5.23E-06 | 5.93E-05 | 3.53 |
| *Dlx1* | 40 | 0.006400 | 0.036074 | 2.07 | 34 | 1.23E-15 | 2.08E-14 | 3.60 |
| *Prlr* | 66 | 0.011087 | 0.114386 | 1.86 | 13 | 3.21E-26 | 1.09E-24 | 4.36 |

**Supplemental Table S12**

| Supplemental Table S12. Specific Hypothalamic Nuclei Level Enrichment Analysis for all imprinted genes in neuronal subpopulation in the Arcuate Nucleus dataset (7). *Identity* – neural lineage identities for the cells used in this analysis. All other column descriptions can be found in the legend of Supplemental Table S9A. | | | | | | | | |
| --- | --- | --- | --- | --- | --- | --- | --- | --- |
| Identity | **Up Reg**  **(16,098)** | **IG**  **(95)** | **ORA *p*** | **ORA *q*** | **Mean FC**  **IG** | **Mean FC**  **Rest** | **GSEA**  ***p*** | **GSEA**  ***q*** |
| *Th/Slc6a3* | 1113 | 26 | 7.49E-10 | **1.72E-08** | 2.14 | 4.04 | - | - |
| *Arx/Nr5a2* | 599 | 16 | 3.98E-07 | **9.16E-06** | 1.33 | 3.15 | - | - |
| *Agrp/Sst* | 1704 | 23 | 0.0001 | **0.0026** | 1.61 | 1.78 | - | - |
| *Pomc/Anxa2* | 3162 | 34 | 0.0002 | **0.0039** | 1.70 | 1.82 | - | - |
| *Ghrh* | 1265 | 18 | 0.0004 | **0.0088** | 2.02 | 2.82 | - | - |
| *Gpr50* | 578 | 11 | 0.0006 | **0.0136** | 2.52 | 5.02 | - | - |
| *Pomc/Glipr1* | 1536 | 19 | 0.0014 | **0.0328** | 1.91 | 1.90 | 0.2851 | 0.8553 |
| *Slc17a6/Trhr* | 1547 | 19 | 0.0016 | **0.0357** | 3.22 | 2.35 | 0.1143 | 0.3429 |
| *Th/Cxcl12* | 818 | 12 | 0.0031 | 0.0710 | 1.95 | 3.40 | - | - |
| *Agrp/Gm8773* | 3629 | 33 | 0.0044 | 0.1021 | 2.08 | 1.84 | 0.1032 | 0.3096 |
| *Fam19a2* | 1237 | 15 | 0.0057 | 0.1321 | 1.62 | 3.42 | - | - |
| *Th/Lef1* | 600 | 9 | 0.0089 | 0.2052 | 1.54 | 3.98 | - | - |
| *Tbx19* | 584 | 8 | 0.0220 | 0.5063 | 1.36 | 3.71 | - | - |
| *Pomc/Ttr* | 1114 | 11 | 0.0634 | 1 | 1.92 | 2.21 | - | - |
| *Gm8773/Tac1* | 1628 | 14 | 0.0963 | 1 | 1.51 | 3.14 | - | - |
| *Kiss1/Tac2* | 1087 | 9 | 0.1912 | 1 | 1.55 | 3.86 | - | - |
| *Nfix/Htr2c* | 1719 | 13 | 0.2112 | 1 | 2.93 | 4.36 | - | - |
| *Sst/Unc13c* | 832 | 7 | 0.2200 | 1 | 1.39 | 2.35 | - | - |
| *Th/Sst* | 1605 | 12 | 0.2357 | 1 | 3.00 | 7.62 | - | - |
| *Qrfp* | 751 | 6 | 0.2828 | 1 | 4.58 | 14.58 | - | - |
| *Htr3b* | 1514 | 10 | 0.4034 | 1 | 1.29 | 2.17 | - | - |
| *Sst/Nts* | 3861 | 23 | 0.5191 | 1 | 1.62 | 2.31 | - | - |
| *Sst/Pthlh* | 1609 | 7 | 0.8493 | 1 | 1.81 | 2.38 | - | - |
| *Tmem215* | 498 | 2 | - | - | 1.08 | 2.18 | - | - |

**Supplemental Table S13**

| Supplemental Table S13. Specific Hypothalamic Nuclei Level Enrichment Analysis for all imprinted genes in neuronal subpopulation in the Suprachiasmatic Nucleus (9). *Identity* – neural lineage identities for the cells used in this analysis. All other column descriptions can be found in the legend of Supplemental Table S9A. | | | | | | | |
| --- | --- | --- | --- | --- | --- | --- | --- |
| Identity | **Up Reg**  **(11,391)** | **IG**  **(77)** | **ORA *p*** | **ORA *q*** | **Mean FC**  **IG** | **Mean FC**  **Rest** | **GSEA *p*** |
| *Avp/Nms* | 909 | 24 | 3.01E-09 | **1.51E-08** | 3.12 | 1.94 | **0.004** |
| *Cck/Bdnf* | 831 | 11 | 0.0233 | 0.1167 | 2.17 | 3.09 | - |
| *Vip/Nms* | 816 | 10 | 0.0473 | 0.2367 | 2.54 | 1.95 | - |
| *Cck/C1ql3* | 1287 | 12 | 0.1554 | 0.7769 | 1.69 | 1.97 | - |
| *Vip/Grp* | 2026 | 12 | 0.7385 | 1 | 1.67 | 2.43 | - |

**Supplemental Table S14**

| Supplemental Table S14. Monoaminergic Nuclei Level Enrichment Analysis for all imprinted genes in dopaminergic subpopulations located throughout the mouse brain at P7 and E15.5 (11). *Identity* – neural lineage identities for the cells used in this analysis. All other column descriptions can be found in the legend of Supplemental Table S9A. | | | | | | |  | |
| --- | --- | --- | --- | --- | --- | --- | --- | --- |
| Identity | **Up Reg**  **(13,095)** | **IG**  **(87)** | **ORA *p*** | **ORA *q*** | **Mean FC IG** | **Mean FC Rest** | | **GSEA *p*** |
| Arcuate Nucleus (*Th, Slc6a3, Prlr*) (P7) | 931 | 25 | 8.82E-10 | **1.15E-08** | 1.97 | 1.97 | | 0.5795 |
| Arcuate Nucleus (*Th, Ghrh, Gal*) (P7) | 560 | 15 | 3.69E-06 | **4.79E-05** | 2.96 | 3.32 | | - |
| Periaqueductal Gray (P7) | 667 | 14 | 0.0001 | **0.0015** | 2.67 | 3.54 | | - |
| Midbrain Neuroblast (P7) | 1100 | 18 | 0.0003 | **0.0036** | 2.26 | 2.66 | | - |
| Ventral Tegmental Area (P7) | 1872 | 25 | 0.0004 | **0.0046** | 1.88 | 2.33 | | - |
| Olfactory Bulb Mature Th+ (P7) | 691 | 7 | 0.1750 | 1 | 3.54 | 3.79 | | - |
| Olfactory Bulb immature Th+ (P7) | 128 | 2 | 0.2090 | 1 | 3.06 | 4.18 | | - |
| Olfactory Bulb Maturing Th+ (P7) | 360 | 4 | 0.2173 | 1 | 4.19 | 4.42 | | - |
| Substantia Nigra (P7) | 2553 | 20 | 0.2409 | 1 | 2.19 | 2.51 | | - |
| Post-mitotic midbrain neuron (E15.5) | 1362 | 11 | 0.2929 | 1 | 2.30 | 2.06 | | - |
| Forebrain neuroblast (E15.5) | 3192 | 19 | 0.7474 | 1 | 1.51 | 1.79 | | - |
| Midbrain Neuroblast (E15.5) | 5085 | 29 | 0.8789 | 1 | 1.70 | 2.16 | | - |
| Post-mitotic forebrain Th+ (E15.5) | 1177 | 5 | 0.9013 | 1 | 1.84 | 2.36 | | - |

**Supplemental Table S15**

| Supplemental Table S15. Monoaminergic Nuclei Level Enrichment Analysis for all imprinted genes in cell populations isolated from E11.5- E18.5 ventral midbrain (12) *Identity* – cell identities for the cells used in this analysis. All other column descriptions can be found in the legend of Supplemental Table S9A. | | | | | | | | |
| --- | --- | --- | --- | --- | --- | --- | --- | --- |
| Identity | **Up Reg**  **(14,008)** | **IG**  **(93)** | **ORA *p*** | **ORA *q*** | **Mean FC**  **IG** | **Mean FC**  **Rest** | **GSEA *p*** | **GSEA *q*** |
| Serotonergic Neurons | 1781 | 33 | 1.47E-08 | **3.09E-07** | 4.78 | 2.36 | 0.0272 | 0.0544 |
| Pericytes | 1409 | 22 | 0.0001 | **0.0022** | 8.57 | 8.04 | 0.2209 | 0.4418 |
| Dopaminergic 1 (*Th*) | 347 | 9 | 0.0005 | **0.0103** | 3.17 | 2.84 | - | - |
| Dopaminergic 0 (Immature) | 218 | 7 | 0.0006 | **0.0129** | 4.03 | 2.90 | - | - |
| Lateral Neuroblasts 2 | 2158 | 26 | 0.0014 | **0.0290** | 2.11 | 2.48 | - | - |
| Oculomotor and trochlear nucleus | 1994 | 22 | 0.0101 | 0.2116 | 1.83 | 2.28 | - | - |
| Mediolateral Neuroblasts 3 | 707 | 10 | 0.0187 | 0.3918 | 2.25 | 2.27 | - | - |
| GABAergic neurons 1b | 943 | 12 | 0.0218 | 0.4585 | 1.84 | 2.03 | - | - |
| GABAergic neurons 1a | 743 | 10 | 0.0253 | 0.5317 | 2.05 | 2.12 | - | - |
| Dopaminergic 2 (*Aldh1a1*) | 463 | 7 | 0.0342 | 0.7187 | 3.22 | 3.42 | - | - |
| Endothelial cell | 2554 | 24 | 0.0432 | 0.9075 | 7.38 | 8.31 | - | - |
| Mediolateral Neuroblasts 1 | 302 | 5 | 0.0505 | 1 | 1.76 | 2.24 | - | - |
| GABAergic neurons 2 | 5711 | 46 | 0.0550 | 1 | 1.89 | 1.95 | - | - |
| Radial Glia-like Cells 3 | 1141 | 9 | 0.3455 | 1 | 3.43 | 5.44 | - | - |
| Red Nucleus | 1021 | 8 | 0.3668 | 1 | 2.21 | 2.01 | - | - |
| Mediolateral Neuroblasts 4 | 811 | 6 | 0.4533 | 1 | 1.54 | 2.42 | - | - |
| Microglia | 1283 | 7 | 0.7594 | 1 | 6.06 | 20.70 | - | - |
| Radial Glia-like Cells 1 | 1309 | 7 | 0.7777 | 1 | 3.08 | 2.79 | - | - |
| Ependymal | 1388 | 7 | 0.8270 | 1 | 3.49 | 12.31 | - | - |
| Neuronal Progenitor | 1774 | 9 | 0.8485 | 1 | 2.32 | 2.74 | - | - |
| Radial Glia-like Cells 2 | 1726 | 8 | 0.9002 | 1 | 2.89 | 3.41 | - | - |
| Lateral Neuroblasts 1 | 529 | 4 | - | - | 1.76 | 2.93 | - | - |
| Mediolateral Neuroblasts 5 | 125 | 4 | - | - | 1.86 | 1.92 | - | - |
| Neuroblast Dopaminergic | 196 | 3 | - | - | 3.55 | 2.14 | - | - |
| Medial Neuroblast | 313 | 2 | - | - | 1.48 | 2.34 | - | - |
| Mediolateral Neuroblasts 2 | 206 | 1 | - | - | 1.13 | 1.82 | - | - |

**Supplemental Table S16 (A, B & C)**

| Supplemental Table S16A. Monoaminergic Nuclei Level Enrichment Analysis for all imprinted genes in cell populations isolated from the Dorsal Raphe Nucleus (10). *Identity* – major cell identities for the cells used in this analysis. All other column descriptions can be found in the legend of Supplemental Table S9A. | | | | | | | | |
| --- | --- | --- | --- | --- | --- | --- | --- | --- |
| Identity | **Up Reg**  **(14,391)** | **IG**  **(85)** | **ORA *p*** | **ORA *q*** | **Mean FC**  **IG** | **Mean FC**  **Rest** | **GSEA *p*** | **GSEA *q*** |
| Dopaminergic Neurons | 9408 | 65 | 0.0006 | **0.0089** | 11.23 | 3.02 | 1.00E-04 | **7.00E-04** |
| Serotonergic Neurons | 9496 | 65 | 0.0008 | 0.0125 | 5.83 | 2.92 | 1.00E-04 | **7.00E-04** |
| Glutamatergic Neurons | 10828 | 68 | 0.0087 | 0.1398 | 5.59 | 3.14 | 1.00E-04 | **7.00E-04** |
| GABAergic Neurons | 10588 | 66 | 0.0145 | 0.2323 | 5.10 | 3.06 | 1.00E-04 | **7.00E-04** |
| Peptidergic Neurons | 7066 | 56 | 0.0000 | **0.0008** | 4.91 | 3.60 | 0.0023 | **0.0161** |
| Fibroblasts | 2430 | 19 | 0.0468 | 0.7481 | 9.56 | 8.20 | 0.2697 | 1 |
| Smooth Muscle Cells | 2242 | 16 | 0.1241 | 1 | 7.69 | 7.46 | 0.5246 | 1 |
| Polydendrocytes | 4880 | 38 | 0.0036 | 0.0574 | 2.11 | 2.64 | - | - |
| Pericytes | 1226 | 14 | 0.0046 | 0.0742 | 7.28 | 7.67 | - | - |
| Oligodendrocytes | 1812 | 13 | 0.1530 | 1 | 2.14 | 4.42 | - | - |
| Ependymal Cells | 6214 | 32 | 0.5970 | 1 | 5.14 | 7.43 | - | - |
| Endothelial Cells | 3147 | 15 | 0.7042 | 1 | 3.06 | 6.80 | - | - |
| Differentiating Oligodendrocytes | 6484 | 31 | 0.7812 | 1 | 1.92 | 1.99 | - | - |
| Astrocytes | 2688 | 9 | 0.9560 | 1 | 1.87 | 2.94 | - | - |
| Microglia | 3003 | 9 | 0.9841 | 1 | 3.08 | 6.14 | - | - |
| Perivascular Macrophages | 3354 | 7 | 0.9995 | 1 | 2.23 | 7.45 | - | - |
| Lymphocytes | 1121 | 0 | - | - | 0.00 | 14.87 | - | - |

| Supplemental Table S16B. Monoaminergic Nuclei Level Enrichment Analysis for all imprinted genes in neuronal major populations isolated from the Dorsal Raphe Nucleus (10). *Identity* – neurotransmitter neuron identities for the cells used in this analysis. All other column descriptions can be found in the legend of Supplemental Table S9A. | | | | | | | | |
| --- | --- | --- | --- | --- | --- | --- | --- | --- |
| Identity | **Up Reg**  **(14,391)** | **IG**  **(85)** | **ORA *p*** | **ORA *q*** | **Mean FC**  **IG** | **Mean FC**  **Rest** | **GSEA *p*** | **GSEA *q*** |
| Serotonergic Neurons | 1274 | 18 | 0.0004 | **0.0019** | 4.03 | 2.66 | 0.1603 | 0.3206 |
| Dopaminergic Neurons | 1519 | 15 | 0.0316 | 0.1579 | 4.93 | 2.93 | 0.1004 | 0.2008 |
| Glutamatergic Neurons | 3458 | 28 | 0.0391 | 0.1955 | 1.67 | 1.82 | - | - |
| Peptidergic Neurons | 1298 | 11 | 0.1416 | 0.7082 | 1.83 | 6.42 | - | - |
| GABAergic Neurons | 2796 | 19 | 0.2860 | 1 | 1.34 | 1.76 | - | - |

| Supplemental Table S16C. Monoaminergic Nuclei Level Enrichment Analysis for all imprinted genes in neuronal subpopulations populations isolated from the Dorsal Raphe Nucleus (10). *Identity* – neurotransmitter neuron subpopulation identities for the cells used in this analysis. All other column descriptions can be found in the legend of Supplemental Table S9A. | | | | | | |
| --- | --- | --- | --- | --- | --- | --- |
| Identity | **Up Reg**  **(14,391)** | **IG**  **(85)** | **ORA *p*** | **ORA *q*** | **Mean FC**  **IG** | **Mean FC**  **Rest** |
| Neuron_GABA_*Sst_Asb4*_vlPAG_dl | 1001 | 20 | 1.06E-06 | **1.70E-05** | 1.66 | 2.25 |
| Neuron_Glu_*Vsx2_Pnoc*_vlPAG_vl | 633 | 13 | 8.14E-05 | **0.0013** | 2.23 | 2.56 |
| Neuron_5-HT_*Hcrtr1_Asb4_*DRN_vl | 464 | 11 | 8.62E-05 | **0.0014** | 2.06 | 2.10 |
| Neuron_DA_*Cck_Npw*_DRN_rm | 556 | 12 | 9.85E-05 | **0.0016** | 2.28 | 2.29 |
| Neuron_5-HT_*Prkcq_Trh*_DRN_dl | 848 | 14 | 0.0004 | **0.0066** | 1.89 | 2.10 |
| Neuron_5-HT_*Met_Trpc3*_DRN_cm | 1249 | 15 | 0.0061 | 0.0968 | 3.42 | 5.30 |
| Neuron_DA_*Vip_Prkcd*_DRN_dl | 959 | 12 | 0.0104 | 0.1667 | 2.64 | 2.84 |
| Neuron_5-HT_*Pdyn_Nos1*_DRN_dm | 541 | 8 | 0.0145 | 0.2313 | 2.29 | 2.26 |
| Neuron_Peptide_*Cartpt_Ucn*_EW | 963 | 11 | 0.0259 | 0.4139 | 1.83 | 8.09 |
| Neuron_Glu_*Pax6_Penk*_vlPAG | 1905 | 17 | 0.0517 | 0.8279 | 1.53 | 2.06 |
| Neuron_Glu_*Fign_Pdyn*_vlPAG_cd | 2063 | 17 | 0.0940 | 1 | 1.73 | 2.32 |
| Neuron_GABA_*Kit_Ebf3*_vlPAG | 925 | 9 | 0.0949 | 1 | 1.53 | 2.23 |
| Neuron_5-HT_*Slc17a8_Cbln2*_DRN_vm | 641 | 6 | 0.1777 | 1 | 1.69 | 2.01 |
| Neuron_DA_*Chrna4_Chrnb3*_CLi | 808 | 7 | 0.1998 | 1 | 2.72 | 2.45 |
| Neuron_Glu_*Slc17a8_Gata3*_DRN_dm | 1744 | 12 | 0.3320 | 1 | 1.96 | 3.01 |
| Neuron_GABA_*Asic4_Calb2*_DRN_dl | 1116 | 8 | 0.3379 | 1 | 1.18 | 2.06 |
| Neuron_GABA/Glu_*Crhbp_Ret*_CLi | 888 | 1 | - | - | 2.91 | 3.19 |
| Neuron_Glu_*Nifa_Nfix*_vlPAG_dl | 524 | 3 | - | - | 3.84 | 2.66 |

**Supplemental Table S17 (A & B)**

| Supplemental Table 17A. Imprinted gene over-representation in Anterior Pituitary cells – 10x (13). *Identity –* Cell identities for the cells used in analysis; All other column descriptions can be found in the legend of Supplemental Table S9A. | | | | | | |
| --- | --- | --- | --- | --- | --- | --- |
| Cell Population Identity | **Up Reg**  **(11,175)** | **IG**  **(70)** | **ORA**  ***p*** | **ORA**  ***q*** | **Mean FC**  **IG** | **Mean FC**  **Rest** |
| Somatotrope | 617 | 14 | 2.30E-05 | **0.000184** | 2.97 | 2.92 |
| Lactotrope | 1179 | 15 | 0.005571 | **0.044565** | 1.78 | 2.40 |
| Melanotrope | 1678 | 18 | 0.013409 | 0.107273 | 2.19 | 3.54 |
| Multi-Hormone Cluster | 1233 | 9 | 0.366262 | 1 | 3.11 | 3.11 |
| Stem Cell (*Mki67*+) | 1003 | 6 | 0.608454 | 1 | 4.12 | 9.80 |
| Gonadotrope | 1755 | 9 | 0.790887 | 1 | 3.97 | 4.53 |
| Stem Cell (*Sox2*+) | 1776 | 9 | 0.802769 | 1 | 11.59 | 9.00 |
| Macrophage | 2186 | 8 | 0.975755 | 1 | 51.60 | 75.60 |
| Corticotrope | 320 | 4 | - | - | 2.29 | 5.71 |
| Folliculo-stellate cells | 498 | 3 | - | - | 7.51 | 16.96 |

**Supplemental Table S18**

| Supplemental Table 17B. Imprinted gene over-representation in Anterior Pituitary cells – Dropseq (13). *Identity –* Cell identities for the cells used in analysis; All other column descriptions can be found in the legend of Supplemental Table S9A. | | | | | | | |
| --- | --- | --- | --- | --- | --- | --- | --- |
| Cell Population Identity | **Up Reg**  **(11,444)** | **IG**  **(77)** | **ORA**  ***p*** | **ORA**  ***q*** | **Mean FC**  **IG** | **Mean FC**  **Rest** | **GSEA**  ***p*** |
| Lactotrope | 401 | 11 | 8.71E-06 | **7.84E-05** | 1.72 | 2.14 | - |
| Somatotrope | 1036 | 17 | 2.58E-05 | **0.000232** | 1.97 | 1.87 | 0.4987 |
| Corticotrope | 3291 | 28 | 0.004954 | **0.044589** | 2.75 | 3.44 | - |
| Melanotrope | 2926 | 23 | 0.029694 | 0.267250 | 2.03 | 2.90 | - |
| Gonadotrope | 2604 | 19 | 0.090364 | 0.813277 | 2.78 | 3.28 | - |
| Stem Cell (*Sox2*+) | 2773 | 18 | 0.215493 | 1 | 3.24 | 4.84 | - |
| Folliculo-stellate cells | 1290 | 9 | 0.250136 | 1 | 32.62 | 6.96 | - |
| Multi-Hormone Cluster | 1312 | 9 | 0.265892 | 1 | 1.99 | 2.38 | - |
| Macrophage | 2666 | 6 | 0.997677 | 1 | 75.17 | 46.03 | - |
| Stem Cell (*Mki67*+) | 1280 | 1 | - | - | 3.19 | 8.86 | - |

| Supplemental Table S18. Imprinted gene over-representation in pituitary cells (14). *Identity –* Cell identities for the cells used in analysis; All other column descriptions can be found in the legend of Supplemental Table S9A. | | | | | | | | |
| --- | --- | --- | --- | --- | --- | --- | --- | --- |
| Cell Population Identity | **Up Reg**  **(15,777)** | **IG**  **(92)** | **ORA**  ***p*** | **ORA**  ***q*** | **Mean FC**  **IG** | **Mean FC**  **Rest** | **GSEA**  ***p*** | **GSEA**  ***q*** |
| Somatotropes | 1277 | 22 | 2.98E-06 | **3.57E-05** | 1.99 | 2.60 | - | - |
| Thyrotropes | 486 | 9 | 0.002094 | **0.025123** | 4.39 | 7.32 | - | - |
| Posterior pituitary | 3010 | 26 | 0.020724 | 0.248685 | 14.38 | 14.15 | 0.3636 | 1 |
| Melanotropes | 1759 | 17 | 0.024650 | 0.295798 | 1.85 | 4.47 | - | - |
| Corticotropes | 2728 | 23 | 0.038664 | 0.463974 | 2.37 | 4.66 | - | - |
| Connective tissue | 2832 | 22 | 0.090352 | 1 | 18.95 | 28.50 | - | - |
| Lactotropes | 4535 | 30 | 0.237678 | 1 | 1.98 | 2.02 | - | - |
| Gonadotropes | 8039 | 49 | 0.367498 | 1 | 8.11 | 3.70 | 0.0319 | 0.0957 |
| WBCs | 1620 | 8 | 0.740475 | 1 | 26.02 | 137.56 | - | - |
| Endothelia | 4603 | 22 | 0.892176 | 1 | 20.80 | 15.85 | 0.4886 | 1 |
| Proliferating Pou1f1-cells | 5103 | 22 | 0.970235 | 1 | 1.53 | 5.08 | - | - |
| Pituitary stem cells | 10000 | 49 | 0.982304 | 1 | 3.40 | 7.36 | - | - |

**Supplemental Table S19**

Output of Level 1 Analysis performed using imprinted gene lists excluding imprinted genes thus far only shown to be imprinted in brain tissue. (‘Supplemental_Table_S19_Level_1_Analyses_Non_Brain_Only.xlsx')

**Supplemental Table S20**

| Supplemental Table S20. Dataset specific sequencing and processing information for all datasets analysed. Datasets are organised by level of analysis and compared for single-cell sequencing protocol, Animal and Tissue processing, Cell Quality Filters used, No. of Cells in final dataset and Data Normalisation procedure followed. | | | | | | |
| --- | --- | --- | --- | --- | --- | --- |
| Dataset | **Level** | **Protocol** | **Animal/Tissue** | **Cell Filter** | **Cell No.** | **Normalisation** |
| *Mouse Cell Atlas (1)* | Multi-Organ | Microwell-seq | C57BL/6J , M+F, 6-10 weeks, E14.5 and neonatal | 1500 highest quality cells per tissue | 61,637 | 100,000 transcripts log transformed |
| *Tabula Muris (2)* | Multi-Organ | Smart-seq2 | C57BL/6J 7 mice (4F/3M), 10-15 weeks, virgin | reads>50,000  genes>500 | 44,879 | ln(CPM+1) |
| *Mouse Brain Atlas (3)* | Whole Brain | 10X chromium | CD1, M+F, p12 – 30, week 6 and week 8, | 600<UMI  1.2 UMI:gene | 160,796 | 5,000 reads per cell log transformed |
| *Whole Brain (4)* | Whole Brain | 10X chromium | C57BL/6J, 8 mice 2-3 months, whole brain minus hindbrain | 200<UMI<30,000  250<gene<6,000 | 16,028 | 10,000 reads per cell log transformed |
| *Whole Hypothalamus (6)* | Whole Hypothalamus | STRT-seq (Fluidigm C1) | C57BL/6J, M+F, 14-28 days | Molecules > 1500 (excluding rRNA and mitochondrial RNA) | 2,882 | 10,000 reads per cell log transformed |
| *Whole Hypothalamus (5)* | Whole Hypothalamus | Drop-seq | B6D2F1 mice (C57B6 female × DBA2 male) - 7 Female, 8-10 weeks | Genes>2000 | 3319 | log(TPM+1) |
| *Arcuate Nucleus (ARC) (7)* | Specific Hypothalamic Nucleus | Drop-seq | C57BL/6J – 53mice - 4-12 weeks, virgin, M+F | genes>800 | 20,921 | 10,000 reads per cell, log transformed |
| *Suprachiasmatic Nucleus (SCN) (9)* | Specific Hypothalamic Nucleus | 10x chromium | C57BL/6J at timepoint ZT8 | None stated (but prefiltered cells provided) | 1,251 | 5000 transcripts per cell, log transformed |
| *Dorsal Raphe Nucleus (DRN) (10)* | Monoaminergic Nuclei | inDrop | C57BL/6J -8 mice (4M, 4F), 8-10 weeks | 18,000>UMI>500  6000>gene>200  mito<0.1 | 39,411 | 10,000 UMI per cell log transformed |
| *E11.5 - E18.5 ventral midbrain (12)* | Monoaminergic Nuclei | STRT-seq (Fluidigm C1) | CD1, E11.5 - E18.5 - 271 embryos | 2000>Molecules >26,000 | 1,907 | 10,000 reads per cell log transformed |
| *Whole Brain Dopamine (11)* | Monoaminergic Nuclei | Smart-seq2 | C57BL/6J at E15.5 and P7 | 2000<genes <10,000  1000>RNA >40,000 | 396 | log2(RPC+1) |
| *Pituitary Gland (14)* | Pituitary Gland | 10X chromium | C57BL/6 6 mice, M, 7-week-old | Genes >= 200 | 13,620 | 10,000 UMI per cell log transformed |
| *Anterior Pituitary Gland – 10x*  *(13)* | Pituitary Gland | 10X chromium | CD1, 2M+2F, 7-8 weeks old | Mito < 0.1  200 < Genes < 3000 | 2,780 | 10,000 UMI per cell log transformed |
| *Anterior Pituitary Gland - Dropseq*  *(13)* | Pituitary Gland | Drop-Seq | CD1 1M+1F, 8-week-old | Mito < 0.1  UMI > 300  100 < Genes < 4,000 | 4,663 | 10,000 UMI per cell log transformed |
